# CAM emerges in a leaf metabolic model under water-saving constraints in different environments

**DOI:** 10.1101/2020.01.20.912782

**Authors:** Nadine Töpfer, Thomas Braam, Sanu Shameer, R. George Ratcliffe, Lee J. Sweetlove

## Abstract

Crassulacean Acid Metabolism (CAM) evolved in arid environments as a water-saving alternative to C_3_ photosynthesis. There is great interest in engineering more drought-resistant crop species by introducing CAM into C_3_ plants. However, one of the open questions is whether full CAM or alternative water-saving flux modes would be more productive in the environments typically experienced by C_3_ crops. To study the effect of temperature and relative humidity on plant metabolism we coupled a time-resolved diel model of leaf metabolism to an environment-dependent gas-exchange model. This model allowed us to study the emergence of CAM or CAM-like behaviour as a result of a trade-off between leaf productivity and water-saving. We show that vacuolar storage capacity in the leaf is a major determinant of the extent of CAM and shapes the occurrence of phase II and IV of the CAM cycle. Moreover, the model allows us to study alternative flux routes and we identify mitochondrial isocitrate dehydrogenase (ICDH) and an isocitrate-citrate-proline-2OG cycle as a potential contributor to initial carbon fixation at night. Simulations across a wide range of environmental parameters show that the water-saving potential of CAM strongly depends on the environment and that the additional water-saving effect of carbon fixation by ICDH can reach up to 4% for the conditions tested.

## Introduction

Increasing aridity threatens agricultural productivity not only in hot and dry climates but also in temperate regions where extreme weather conditions are becoming more frequent ^1^. Thus, the development of water-use efficient crop varieties is of utmost importance to maintain food security ^2^. Several plant lineages living in arid environments have evolved CAM photosynthesis, a water-saving mode of C-fixation in which CO_2_ uptake into the mesophyll cell and fixation by RuBisCO are temporally separated ^3^. In CAM photosynthesis the stomata open at night and CO_2_ is fixed and stored in the vacuole in the form of a carboxylic acid such as malate or (iso-)citrate ^4–7^. During the day the stored CO_2_ is remobilized for fixation by RuBisCO in the chloroplast, accompanied by the accumulation of storage carbohydrates. This cycle is considered to be an energetically expensive, but water-use-efficient, alternative to direct day-time CO_2_-fixation by RuBisCO (C_3_ photosynthesis) ^8,9^.

The implementation of CAM photosynthesis into a C_3_ crop plant is a promising engineering target for two reasons: first, all enzymes required for the CAM cycle are already present in C_3_ plants (although specific isoforms with different regulatory properties are required ^10^); and second, some facultative CAM species, such as ice plant (*Mesembryanthemum crystallinum*), can be induced to switch from C_3_ to CAM photosynthesis by a number of environmental factors such as drought or high-salinity ^8,11^ suggesting that it should be possible to engineer CAM into a C_3_ leaf. However, CAM photosynthesis is usually considered to be advantageous in hot and arid climates where water-use efficiency (WUE) is a strong determinant for plant growth and where the suppression of photorespiration through carbon concentration behind closed stomata becomes a considerable factor that balances the additional cost of running the expensive CAM cycle ^12^. To test this hypothesis, in a previous study, we investigated the energetics and productivity of CAM and found that despite 3-fold higher energy consumption at night, the additional cost of running a CAM cycle can be balanced by the carbon-concentrating effect of carboxylic acid decarboxylation behind closed stomata during the day ^13^.

Here we addressed a different question: what are the metabolic and morphological limitations to implementing a water-saving CAM or CAM-like mechanism in a C_3_ leaf and what is the extent of the water-saving effect in different environments? To address this question, we constructed a time-resolved, large-scale metabolic leaf model and coupled it to a gas-exchange model that includes the two main determinants of water-loss through the stomata — the temperature (T) and the relative humidity (RH). This environment-coupled model was used to investigate emergent flux modes when water-saving constraints are applied in addition to high productivity.

## Results

### Model construction

Light availability and gas exchange (CO_2_, water vapour) are major determinants of the metabolic behaviour of a plant leaf. To model the interplay between leaf productivity and water-loss through transpiration we extended a previous diel flux-balance modelling framework ^13^ in two ways. First, we increased the temporal resolution from a binary day-night scenario to modelling a 24 time-step diel cycle, where each interval represents one hour of the day. The time-resolution in the models was achieved by coupling 24 copies of the model in way that a range of metabolites (starch, sugars, amino acids, carboxylic acids, and nitrate) were allowed to accumulate and subsequently be degraded in the plastid and vacuole, respectively. For each of these metabolites we introduced *linker fluxes* which transfer the accumulated metabolite from one time interval to the next. Upper bounds were placed on the quantity of carboxylic acids and other compounds that were allowed to accumulate in the vacuole based on vacuole size and leaf anatomy (leaf thickness and porosity) for average C_3_ and CAM leaves. Secondly, we coupled the metabolic model to a simplified gas-diffusion model which allows us to compute water-loss through the stomata according to the CO_2_ demand of the metabolic system and the environmental conditions (T and RH). A detailed description of the model construction and the exchange constraints is given in the Materials and Methods section. The resulting time-resolved, environment-coupled model enabled us to simulate the effect of the diel light curve, T and RH on leaf metabolism and was used to study the trade-offs between leaf productivity and water-use efficiency (Figure 1).

**Figure 1:**
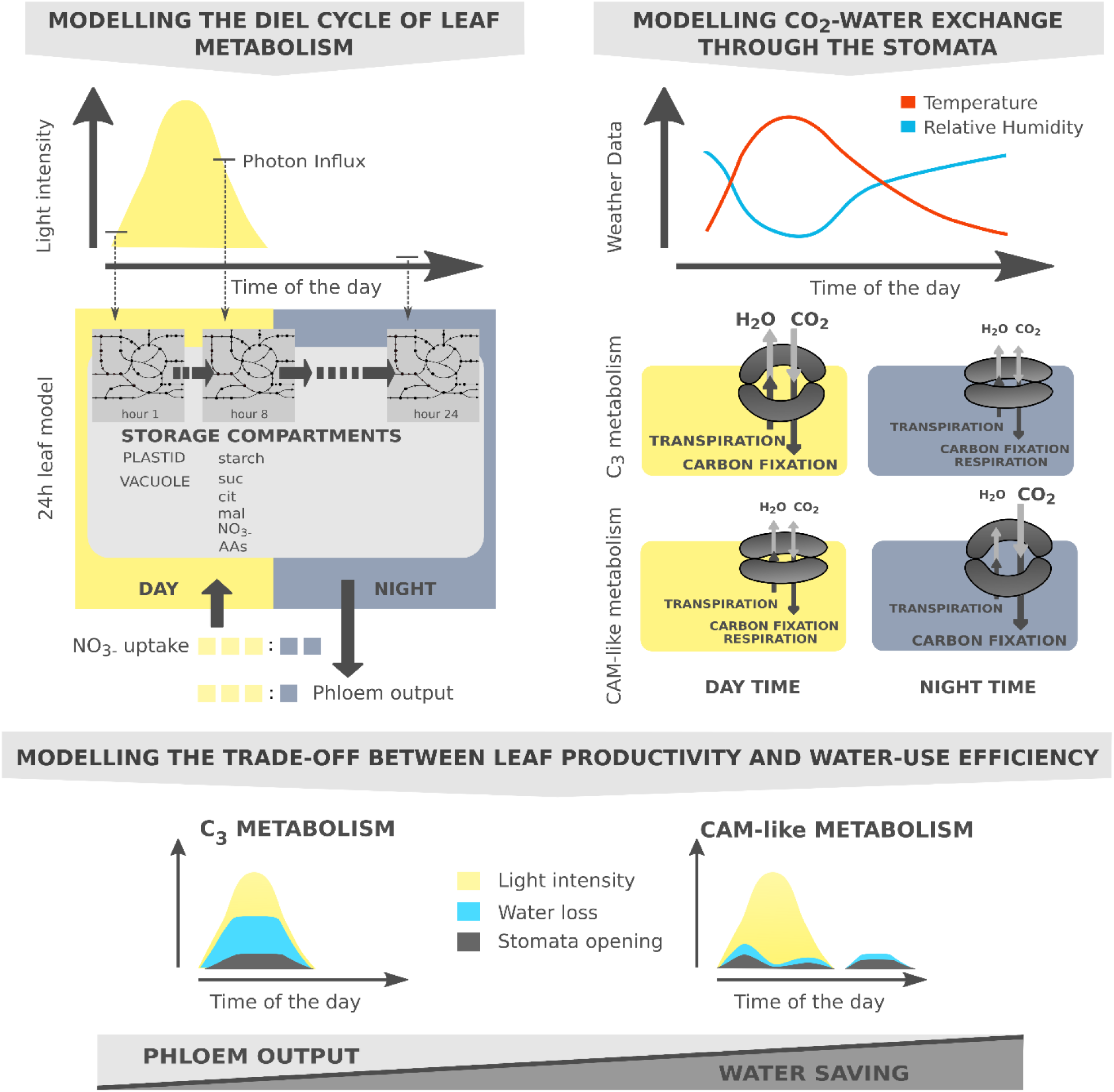
Modelling water-saving flux modes in an environment-coupled model of leaf metabolism. Upper left: A 24h-leaf model was constructed by concatenating copies of a core model of plant metabolism ^13^. The individual models were connected via ‘linker fluxes’ which allowed the transfer of storage compounds in the vacuole and the plastid between successive models. Light uptake was constrained by the diel light curve. The day:night ratios of phloem output and maintenance were set to 3:1 for each hour of the diel cycle, and N uptake was constrained to a ratio of 3:2 based on previous estimates ^50^. Upper right: The effect of temperature T and relative humidity RH on stomatal water loss was modelled by a simplified gas-diffusion equation. T and RH data determined the relationship between CO_2_ uptake and water loss. In C_3_ plants stomata open during the day and CO_2_ uptake and water loss are high due to high T and low RH. At night the stomata are closed and gas/water exchange is minimized. In CAM plants stomata remain closed during the day and open at night allowing high CO_2_ influx while having a low transpiration stream due to low T and high RH. Bottom: Combining metabolic and gas-exchange models allows the trade-off between productivity and water-loss to be studied as competing objectives on a Pareto frontier and reveals alternative water-saving C-fixation mechanisms.

In this study we considered the metabolism of a mature leaf and started the analysis by using the maximization of phloem output over the course of the day as the primary objective. This optimality criterion led the metabolic system to synthesize storage compounds in the light which were then used to sustain night-time metabolic processes such as phloem output, maintenance, and nitrogen assimilation in an overall optimal manner. In accordance with the metabolic mode of the system, the model predicted changing CO_2_-demand and — depending on T and RH — water-loss by transpiration over the course of the day. Conversely, we could fix phloem output to a given value (equal to or less than the maximum value) and used minimization of water-loss as a driving force to act on the metabolic system. These constraints led to the prediction of water-saving flux modes while maintaining high productivity.

### Model Analysis & Biological Implications

#### A time-resolved diel model simulates dynamics of C_3_ metabolism

To establish a reference model for a mature leaf operating under optimal conditions we began our analysis by simulating an energy-limited scenario (due to the maximisation of phloem output for a fixed light input) without water-saving constraints. We simulated a typical summer day in a temperate climate with a maximum T of 26°C and a maximum RH of 0.8 (Figure 2A). We used a light curve that peaks at a moderate intensity of 250 μmol m^-2^ s^-1^ and follows a normal distribution with a day length of 12 hours. As a second optimization criterion we minimized the metabolic flux sum ^14^. This objective was used as a proxy for the cost of providing the enzymes for the active reactions. Applying it as a second optimality criterion left the objective value, here the phloem output, unaltered, but chose the flux distribution with the least enzymatic cost from a set of alternatives.

**Figure 2:**
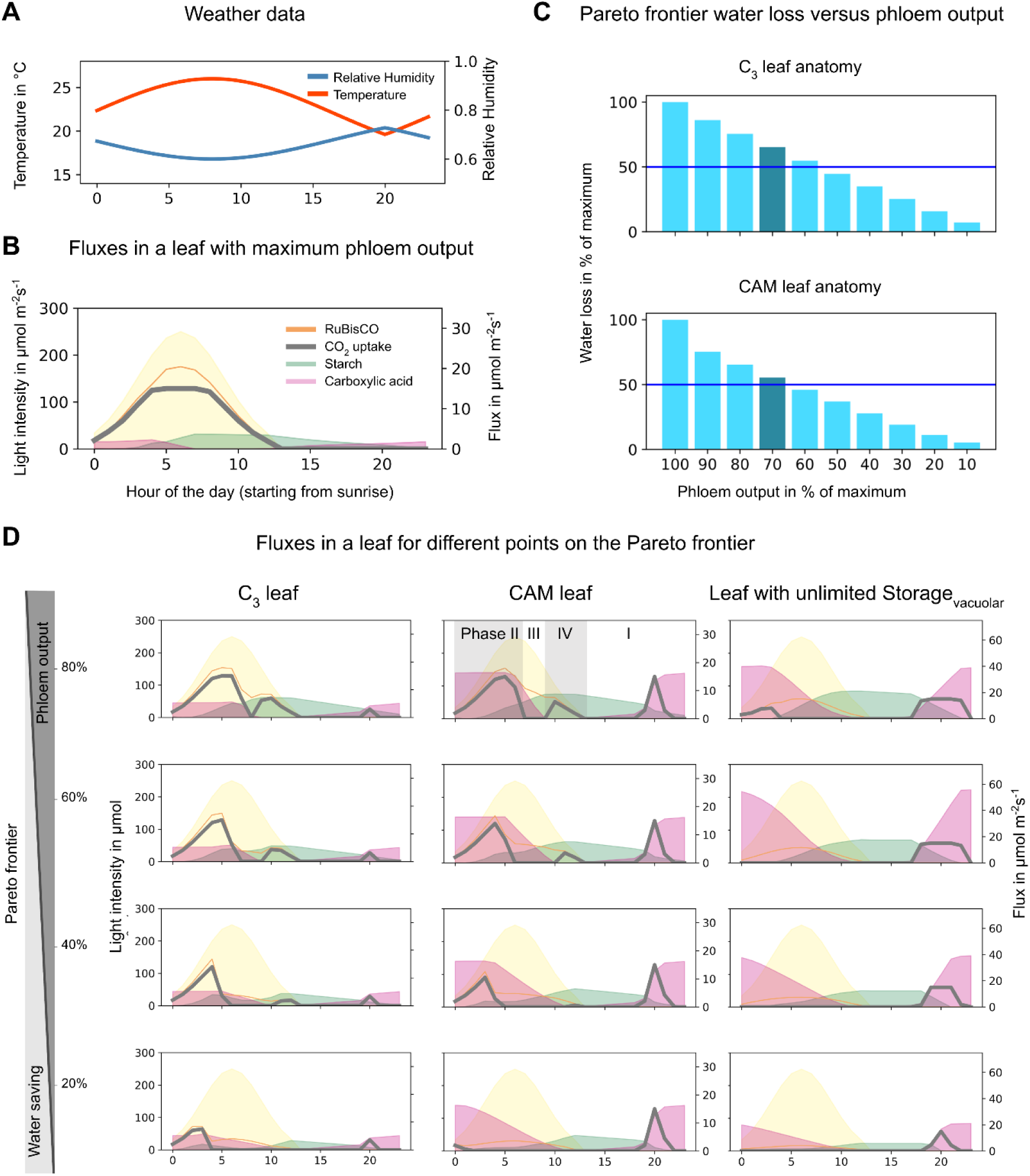
Metabolic fluxes and water loss for different modelling scenarios. (A) Example weather data used throughout the simulations. (B) Metabolic flux profiles in a C_3_ leaf optimized towards phloem output (100% phloem output). The diel light curve is indicated in yellow and peaks at a maximum intensity of 250 μmol m^-2^ s^-1^. Selected fluxes for reactions and linker fluxes demonstrate that the model is operating in C_3_ photosynthesis mode. (C) Pareto analysis of phloem output versus water-loss in a C_3_ leaf (top) and a CAM leaf (bottom). The CAM leaf enabled a better trade-off between the two competing objectives. (D) Metabolic flux profiles for (left) a C_3_ leaf, (middle) a CAM leaf, and (right) a leaf with unlimited vacuolar storage capacity for different Pareto steps (80, 60, 40, and 20% of the max phloem output (shown for a C_3_ leaf in A). The C_3_ leaf exhibits ‘daytime depression’ behaviour, the CAM leaf exhibits all 4 phases of the CAM cycle (phase I: open stomata at night, phase II: open stomata in the early morning hours, phase III: closed stomata during the day, phase IV: open stomata during evening hours). The leaf with unlimited storage exhibits a full CAM cycle below 80% productivity. Note the different flux scales on the right plot axis for the C_3_ and CAM leaf and the leaf with unlimited storage capacity.

Using this setup, the model predicted a total phloem output of 41.3 mmol m^-2^ leaf d^-1^. Daily total water loss was predicted to be 116.4 mol m^-2^ leaf which is 2.1 L m^-2^ leaf and CO_2_ uptake was 446.4 mmol m^-2^ leaf. This resulted in 260.7 mol H_2_O lost per CO_2_ fixed. For extreme conditions with a maximum T of 40°C and a maximum RH of 0.4 this value increases to 955.3 mol H_2_O lost per CO_2_ fixed. Comparing this value to the rule of thumb for C_3_ plants “900-1200 moles H_2_O per mol CO_2_ fixed” ^15^ we found that our model is in broad agreement with experimental observations. This was reassuring, given the simple nature of the gas-exchange model and the fact that the water-loss was predicted from the model’s demand for CO_2_ to maximize phloem output — thereby approving the choice of the objective function.

To get a better overview of the metabolic behaviour over the course of the diel cycle we examined the CO_2_ uptake, RuBisCO activity and the linker fluxes for starch and carboxylic acids, respectively as shown in Figure 2B. The magnitude of a linker flux corresponds to the amount of the stored metabolite, i.e. a flux of 1 μmol m^-2^ s^-1^ means that 3.6 mmol are available for utilisation in subsequent time intervals in the model. Both, CO_2_ uptake (grey line) and RuBisCO flux (orange) were predicted to follow the light curve and peaked at midday coinciding with light availability. Carboxylic acid levels (magenta area) peaked before midday and remained low from before sunset to dawn. Starch (green area) accumulated during day-time hours and was subsequently degraded to sustain metabolism at night. Overall, the described flux patterns were characteristic for C_3_ leaf metabolism. From this starting point we then asked the question: how will the metabolic fluxes change if we change the optimality criterion from maximizing phloem output to minimizing water loss?

#### An optimality study reveals trade-offs between productivity and WUE

Computationally, the question of how a system’s behaviour changes when operating between competing objectives can be tackled by performing a Pareto analysis ^16–19^. In our case phloem output and water-saving represented two competing driving forces. We started the Pareto analysis from the above described scenario of a mature leaf optimized for maximum phloem output (i.e. 100% phloem output, here termed Pareto step 1). We then subsequently reduced the required phloem output in 10%-steps and used minimization of water loss as the primary optimization objective. Given this setup, we saw an almost linear decrease in water loss with decreasing phloem output, hence we did not observe any significant water-saving mechanism in our model (Figure 2C top). Inspection of the CO_2_ uptake and RuBisCO reaction flux in the model showed that with decreasing phloem output, the model closed the stomata during the warmest and driest hours of the day, a phenomenon known as midday depression of photosynthesis (Figure 2D left column), which was accompanied by a very minor peak of CO_2_ uptake at night.

One possible explanation for the lack of water-saving metabolic modes in the model was that the model was limited by the constraints we applied to mimic C_3_ leaf anatomy (e.g. total vacuolar volume per unit of leaf). To test this, we examined the differences between C_3_ and CAM leaf anatomy and adjusted the vacuolar storage constraints accordingly. Using morphological data for an average CAM leaf resulted in a 3.1-times larger vacuolar storage capacity per unit leaf compared to a C_3_ leaf. (see Supplementary Text). When repeating the Pareto analysis using this CAM-morphology, a non-linear relationship between productivity and water loss emerged and the model predicted almost 50% water-saving at 70% of the maximum phloem output (Figure 2C bottom). It is worth noticing that the upper limit for the vacuolar storage capacity had only a very minor impact on the maximum phloem output of the model. The output at Pareto step 1 for the C_3_ leaf model was 99.7% of the phloem output of the CAM leaf. Therefore, in subsequent analyses we directly compared between the two sets of simulations.

What was causing the non-linearity in the relationship between productivity and water loss? As in the C_3_-anatomy-constrained model, we observed a closure of the stomata and a reduction of RuBisCO activity during the hottest and driest hours of the day. However, in addition to these day-time changes to reduce water loss, we also observed a substantial peak of CO_2_ uptake at night which was accompanied by an accumulation of carboxylic acids in the vacuole at night and a greater amount of starch stored during the day and degraded at night. (Figure 2D middle columns). These observations suggested that the model was performing a CAM or CAM-like cycle in which CO_2_ was initially fixed at night and stored in the form of carboxylic acids. During the day, when sufficient light energy was available, CO_2_ was released from its intermediate storage and re-fixed for triose phosphate synthesis during the day using RuBisCO and the Calvin Benson cycle. This was confirmed by inspection of the complete set of predicted fluxes in the model (see Supplementary Text).

When inspecting the Pareto frontier, we observed the steepest slope (i.e. the largest increase in water-saving) between 90% and 100% of the maximum phloem output. At 90% maximum phloem output the model already switched to CAM-like behaviour and partially closed the stomata during the day and reopened them for a short period at night. Comparing this to the almost linear Pareto frontier for the C_3_ leaf indicated that the additional effect of night-time CO_2_-fixation contributed largely to the water-saving.

#### Vacuolar storage capacity limits WUE and influences the extent of phases II and IV of the CAM cycle

When investigating the CO_2_ uptake at different steps along the Pareto frontier it became apparent that the model did not exhibit a full CAM-cycle (Figure 2D middle columns). The stomata closed only for a few hours in the day and re-opened only for a short period at night. The sharp CO_2_ uptake peak at night-time can be explained by three factors. First, the applied RH and T curves have a sharp local maximum and minimum (see Figure 2A). This occurs where the lower and upper ends of the normal curves meet to close a diel cycle. Secondly, our model is anticipating: i.e., the solution for time-point t depends on the environmental parameters to be encountered at time-point t+1. Thirdly, the upper-bound on the rate of CO_2_ uptake was based on the largest CO_2_ uptake rates measured. Therefore, very high rates over a short period were permitted, whereas in reality, under most conditions and in most species, CO_2_ diffusion constraints would cause that CO_2_ is fixed at lower rates but for a longer duration. These factors led to the observed behaviour where CO_2_ uptake showed a short burst when water loss through transpiration was the lowest.

During the day, the stomata remained open for CO_2_ exchange during the early hours of the day and re-opened in the evening hours before sunset. This behaviour was exhibited for all Pareto steps meaning that night-time CO_2_-fixation alone was not sufficient to sustain the required phloem output. The observed opening and re-opening of the stomata during the day occurs in certain CAM species and is known as phase II and IV of the CAM cycle ^3^. Night-time stomata opening for CO_2_-uptake and day-time stomata closure are referred to as phases I and III, respectively. Some CAM species show a remarkable plasticity with respect to these four phases and the reasons for the occurrence and extent of these distinct patterns is still debated ^20–22^. Given the indication that vacuolar storage capacity had a major impact on the night-time CO_2_ uptake pattern in the model and the fact that some CAM species exhibit a bi-phasic CAM cycle we wondered if we might underestimate the vacuolar storage capacity of an average CAM leaf. We therefore repeated the Pareto analysis using the same model but without any vacuolar storage constraints. The results of this analysis are shown in Figure 2D, right column. Without any limitation of the vacuolar storage capacity the model performed a bi-phasic full CAM cycle at 70% of the maximum productivity and below i.e. the stomata remained closed during the day and were open throughout the night, without the appearance of phase II and IV of the CAM cycle. *Therefore, our model suggested that keeping the stomata open for at least a portion of the day was necessary to sustain a high productivity when vacuolar storage capacity is limiting.*

#### Night-time C-fixation by ICDH contributes to WUE

The occurrence of the four phases of CAM in our model raised the question of how metabolic fluxes were distributed during these metabolically distinct phases. To analyse the underlying flux modes in more detail we focused the analysis on a model with a vacuolar storage capacity of a CAM leaf at 70% of maximum productivity (phloem output) optimized for water-saving. We chose this value, as a yield penalty of 30% would be an acceptable trade-off if water usage could be reduced by almost half. We followed the flux of CO_2_ (including bicarbonate) from the stomata through the metabolic system by plotting time-resolved fluxes of all reactions that use CO_2_ or bicarbonate as either a reactant or product (Figure 3A left). During the day, RuBisCO fixed the majority of CO_2_ available from gas-exchange and released by metabolic processes. Cytosolic isocitrate dehydrogenase (ICDH), glycine oxidation in the photorespiratory pathway (glycine decarboxylase), carbamate kinase in N metabolism and NADP-malic enzyme in the cytosol were the main CO_2_-releasing processes during the day. To our surprise, we found that night-time CO_2_ fixation in the model was shared between two enzymes — PEP-carboxylase (PEPC) in the cytosol and ICDH in the mitochondria. While PEPC’s role in CAM photosynthesis is well established, mitochondrial ICDH activity has not been previously linked to this metabolic cycle. In order for ICDH to be used for CO_2_ fixation it has to operate in the reverse direction to its conventional direction in the TCA cycle. This is possible given an appropriate mass action ratio (e.g. due to a high 2OG concentration) and indeed this reaction has been shown to operate in the reverse direction in several *in vivo* metabolic flux studies in developing rapeseed and soybean embryos ^23–25^. ICDH has also been suggested as a kinetically acceptable option for synthetic carbon fixation pathways (ΔG = 21 kJ mol^-1^ at pH 7, ionic strength of 0.1 M, and reactant concentrations of 1 mM ^26,27^). For convenience, we refer to this reaction as *ICDH*_*rev*_.

**Figure 3:**
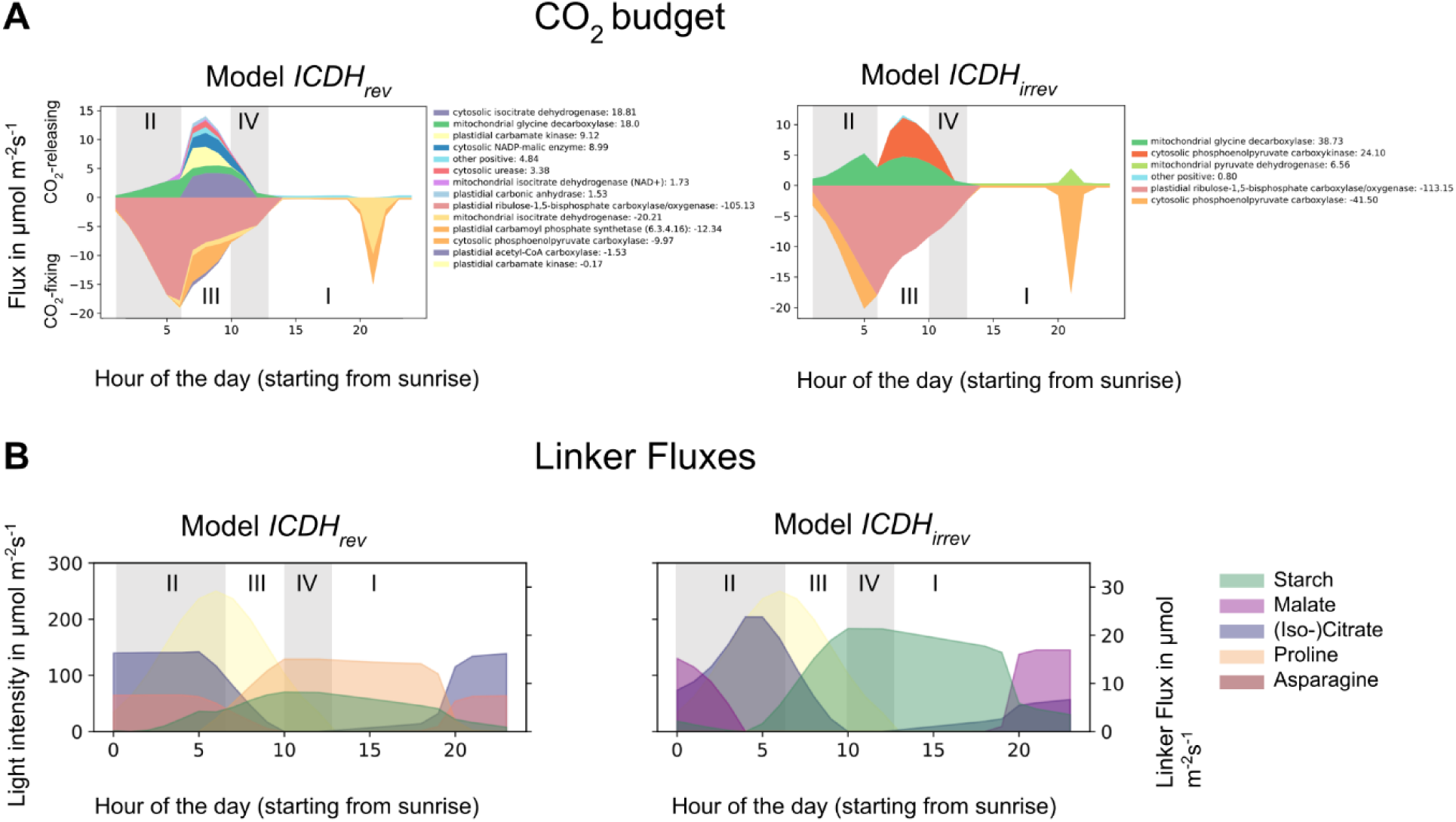
Different flux distributions in a water-saving CAM leaf at 70% productivity with (model *ICDH*_*rev*_) and without (model *ICDH*_*irrev*_) reversible mitochondrial IDCH. (A) CO_2_ budget for the two models reveals different CO_2_ turnover fluxes over the course of the day. Shown are all reactions with a flux > 0.5 μmol m^-2^ s^-1^ for at least one time point. I-IV indicates the 4 phases of the CAM cycle. The values for the cumulative contribution are given next to the reaction name. (B) Significant linker fluxes for both models. Model *ICDH*_*rev*_ accumulated (iso-)citrate as carboxylic acids and additionally Pro and Asp. Model *ICDH*_*irrev*_ accumulated both malate and (iso-)citrate but no amino acids. Starch levels in model *ICDH*_*irrev*_ were almost three-fold higher than in model *ICDH*_*rev*_.

Analysis of the linker fluxes revealed that citrate and/or isocitrate were the sole carboxylic acid to accumulate at night (Accumulation of either citrate or isocitrate or of both carboxylic acids resulted in the same phloem output and water-saving.). Additionally, two amino acids accumulated — Asn during the night and Pro during the day (Figure 3B left). None of the other linker reactions in the vacuole carried a significant flux. Closer inspection of the metabolic fluxes revealed an alternative CO_2_ fixation pathway where both PEPC and ICDH contribute to night-time CO_2_-fixation. An overview of the reactions involved is shown in Figure 4A.

**Figure 4:**
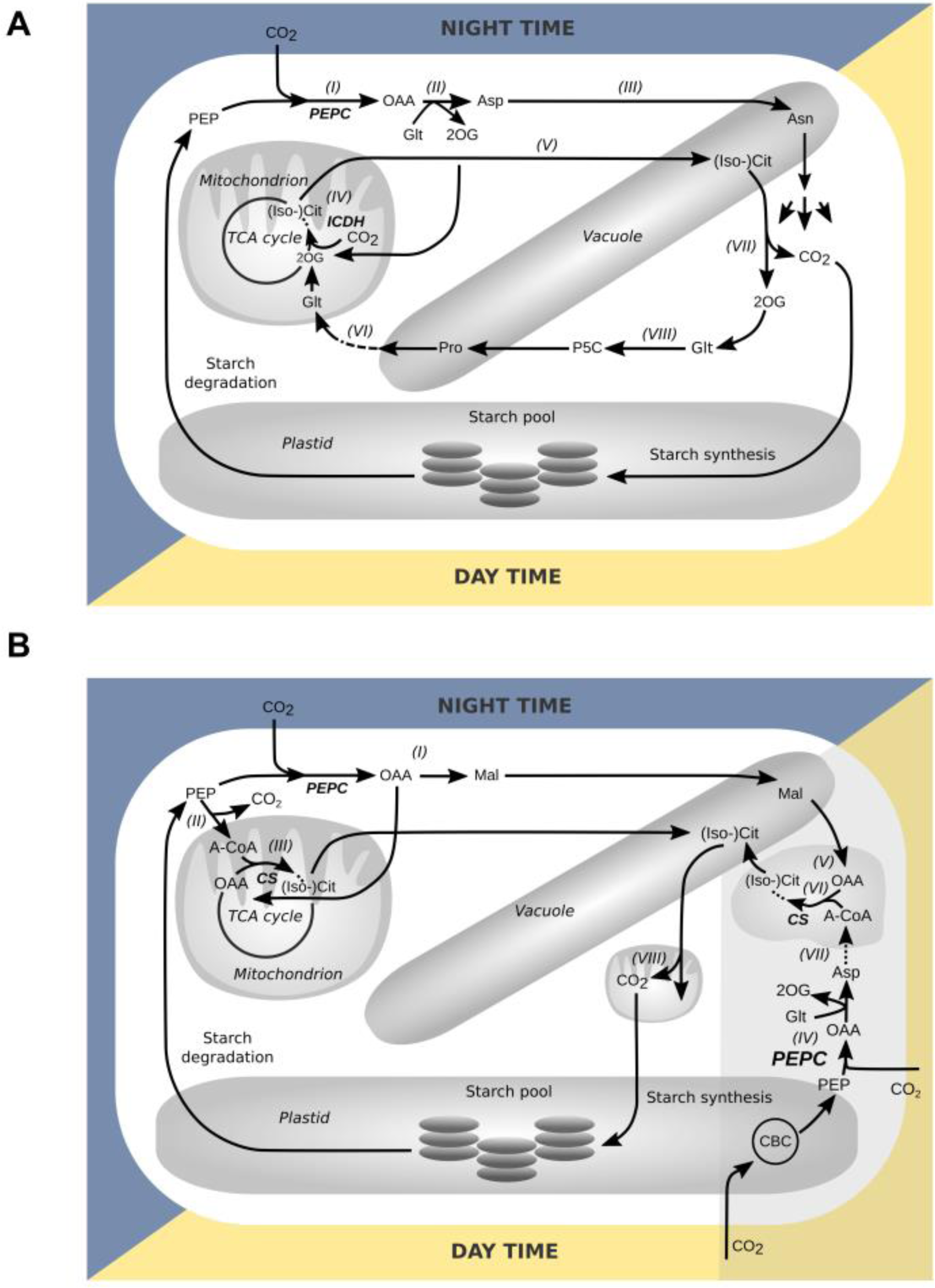
Major flux routes involved in the CAM-like temporally-separated C-fixation mechanism in a water-saving CAM leaf at 70% productivity (A) with (model *ICDH*_*irrev*_) and B) without (model *ICDH*_*irrev*_) reversible mitochondrial IDCH. The two models used different pathways to fix and release C. CS, citrate synthase; ICDH, isocitrate dehydrogenase; PEPC, PEP-carboxylase; 2OG, 2-oxoglutarate; A-CoA, acetyl-coenzyme A; Asn, asparagine; Asp, aspartate; (Iso-)Cit, (Iso-)citrate; Glt, glutamate; Mal, malate; OAA, oxaloacetate; P5C, 1-pyrroline-5-carboxylic acid; PEP, phosphoenolpyruvate; Pro, proline; TCA cycle, tricarboxylic acid cycle; CBC, Calvin-Benson-Bassham cycle. The grey area in B) highlights those reactions that are active in phase II.

At night, when the stomata are open and CO_2_ can enter the leaf, PEPC catalyses the fixation of CO_2_ to PEP (marked as *(I)* in Figure 4A). The resulting oxaloacetate (OAA) together with glutamate was converted to 2OG and Asp by aspartate aminotransferase *(II)*. Asp was converted to Asn and stored in the vacuole *(III)*. 2OG was translocated to the mitochondria as a substrate for *ICDH*_*rev*_ which catalyses the carboxylation of 2OG to isocitrate *(IV)* which was either directly stored in the vacuole or further converted to citrate and then stored in the vacuole for day-time usage *(V)*. Additionally, conversion of the vacuolar pool of Pro to 2OG supported the flux through ICDH in the mitochondria *(VI). Net, this pathway — starting from PEP to the carboxylic acids stored in the vacuole — can fix one mol of CO*_*2*_ *per mol stored (iso-)citrate or Asn.*

During the day, (iso-)citrate from the vacuole was converted to 2OG and CO_2_ by cytosolic ICDH and CO_2_ was refixed in the Calvin-Benson-Bassham (CBB) cycle *(VII)* and ultimately stored as starch to then support night-time metabolism. 2OG was converted to Pro and stored in the vacuole for the next night-time period *(VIII)*.

Does the shared night-time carbon fixation between PEPC and ICDH represent an advantage and if so to what extent is it more beneficial than using PEPC alone? To answer this question, we set ICDH to be irreversible in the conventional forward direction (*ICDH*_*irrev*_) and re-ran the simulations. We found that the additional carboxylating activity of ICDH increased water-saving by 2% (for this particular set of parameters). *From these observations we concluded that night-time C-fixation by ICDH in combination with day-time storage of Pro as a precursor for 2OG might act as an additional water-saving mechanism by adding to the temporal separation of initial CO*_*2*_ *fixation and the activity of the CBB cycle.*

#### PEPC also fixes CO_2_ in the early hours of the day when ICDH is irreversible

How do the metabolic fluxes in our model differ when ICDH_rev_ is not available for C-fixation and how does it affect WUE? A first inspection of the metabolic fluxes for this scenario revealed that making ICDH irreversible had a major impact on the accumulation pattern of both carboxylic acids and starch (Figure 3B right). While the overall diel pattern of carboxylic acid accumulation and degradation was very similar in the two scenarios, the individual patterns for malate and (iso-)citrate were different. When ICDH was irreversible, we observed both malate and (iso-)citrate accumulation at night and a drop of malate levels in the early hours of the day together with an increase of (iso-)citrate levels in the morning hours. Day-time starch levels were more than twice as high as in the scenario where ICDH is reversible and the onset of starch accumulation was shifted towards the later hours of the morning. Pro and Asn did not accumulate. Closer inspection of the flux routes involved in the C-fixation cycle revealed differences between the two scenarios (Figure 4B). At night, PEPC fixed CO_2_ to PEP and formed OAA which was partially converted to malate and stored in the vacuole *(I)* and partially transported to the mitochondria. Additionally, PEP was converted to acetyl-CoA which released one mol CO_2_ per mol acetyl-CoA *(II)*. This loss of CO_2_ was compensated by the activity of citrate synthase in the mitochondrion which converted acetyl-CoA and OAA to citrate *(III)* which was then stored in the vacuole as citrate or isocitrate. Therefore, the synthesis of (iso-)citrate from PEP is C-neutral. *Net, this pathway — starting from PEP to the carboxylic acids in the vacuole — can store 1 mol of CO*_*2*_ *per mol malate and it refixes one mol of CO*_*2*_ *(which was lost during PEP to acetyl-CoA conversion) and stores it as (iso-)citrate. In contrast — when both ICDH and PEPC were active — both, malate and Asn contributed to C-fixation. Therefore, in terms of CO*_*2*_*-fixation capacity the pathway was less efficient than using both PEPC and ICDH.*

During the day the two models showed marked differences in the flux routes between phase II and phase III of the CAM cycle. While model *ICDH*_*rev*_ predicted that CO_2_ fixation through PEPC was limited to night-time, model *ICDH*_*irrev*_ showed additional PEPC activity during phase II in parallel with RuBisCO (Figure 3A right). This early morning PEPC activity increased the amount of CO_2_ that could be transiently stored in the vacuole until sufficient light-energy was available for starch synthesis. PEPC used PEP delivered from the CBB cycle as a substrate *(IV)* to generate OAA. At the same time malate was released from the vacuole and converted to OAA by malate dehydrogenase in the peroxisome *(V)*. A part of the OAA pool and acetyl-CoA were substrates for citrate synthase in the peroxisome *(VI).* The other part of the OAA pool, together with GLT, was used by an aminotransferase to generate 2OG and Asp in the cytosol *(VII)*. Asp was further metabolized and the downstream product acetyl-CoA (see sequence of reactions in Supplementary Text) acted as a precursor for the synthesis of citrate by citrate synthase. (Iso-)Citrate replaced malate in the vacuole. PEPC is known to be active in phase II ^28,29^, however subsequent metabolic flux modes in the model were different from the canonical CAM cycle where malate is decarboxylated to PEP or pyruvate by phosphoenolpyruvate carboxykinase or malic enzyme. Later during phase III, (iso-)citrate was released from the vacuole and supplied CO_2_ for the CBB cycle via a degradation route that involved the glycine decarboxylase system in the mitochondria *(VIII)* (reaction sequence in Supplementary Text). To test if the observed malate to (iso-)citrate exchange in the vacuole during phase II of the CAM cycle was indeed a water-saving advantage, we simulated a scenario where (iso-)citrate uptake into the vacuole was blocked during the day (termed *model ICDH*_*irrev, Cit_night*_). This constraint increased water usage at 70% productivity by 1.1%.

#### High enzyme costs might outweigh the water-saving effect of alternative flux routes

The occurrence of specific metabolic patterns was not only determined by water-use efficiency but also by the cost for enzyme synthesis. In our analysis, this metabolic investment was only indirectly considered by minimizing the metabolic flux sum after the leaf productivity and water loss had been determined. Therefore, flux minimization did not represent a competing objective on the Pareto frontier and a slightly more water-efficient solution with a high enzymatic investment (high flux sum) would always be preferred over a slightly worse performing mechanism with less enzyme investment. To account for this bias, we considered the metabolic flux sum for the three models — *ICDH*_*rev*_, *ICDH*_*irrev*_, and *ICDH*_*irrev, Cit_night*_: the values were 11,455, 13,001, and 11,350 μmol m^-2^ s^-1^, respectively. As a second indicator for metabolic efficiency we considered the overall ATP budget, that is all ATP produced and consumed over the course of the day (Supplementary Text, Supplementary Figure 1). These values indicated the highest ATP turnover of 882 μmol m^-2^ s^-1^ for model *ICDH*_*irrev*_ and lower values of 806 and 796 μmol m^-2^ s^-1^ for models — *ICDH*_*rev*_ and *ICDH*_*irrev, Cit_night*_, respectively. From these observations we conclude that the additional metabolic cost of exchanging malate for (iso-)citrate in the phase II of the CAM cycle would very likely outweigh the water-saving effect of this mechanism. On the other hand, the other two modelling scenarios — model *ICDH*_*rev*_ and model *ICDH*_*irrev,Cit_night*_ had similar metabolic flux sums as well as ATP budgets, indicating that ICDH’s contribution to CO_2_-fixation might indeed be a feasible prediction with respect to enzyme cost.

#### CAM’s WUE depends on the environment

So far, we have focused our analysis on one particular environmental scenario. In the next step we used the model to study the impact of different environments on the trade-off between productivity and WUE focusing on two questions: in which environments is the introduction of a CAM-cycle of the greatest advantage? and how does the contribution of ICDH to C-fixation impact WUE in different environments? To systematically scan the space of possible environments we chose three alternative light regimes with a maximum of 250, 500 and 1000 μmol m^-2^ s^-1^ and three different day:night-time ratios, i.e., 16:8, 12:12 and 8:16. For each of these nine combinations we then scanned for combinations of RH_min_ and RH_max_ between 0.3 and 0.9 and across a temperature regime with night-time temperatures between 0 and 30°C and day-time temperatures between 0 and 50°C (Figure 5A, Supplementary Table 2). For each of these conditions we simulated model *ICDH*_*irrev*_ and model *ICDH*_*rev*_ and analysed the resulting CAM-like behaviour at 70% of the maximum phloem output (100% phloem output corresponds to C_3_ metabolism). We determined the absolute water loss, the relative water loss with respect to the C_3_ model, and the difference between model *ICDH*_*rev*_ and model *ICDH*_*irrev*_. From these analyses we made the following observations: Overall, we found that for the investigated environmental conditions, the relative water loss at 70% productivity with respect to a C_3_ leaf ranged between 31 and 67 % and the additional water-saving effect of ICDH_rev_ reached up to 4%. As an example, the results for a set of simulations for a maximum light intensity of 250 μmol m^-2^ s^-1^, a 12:12 day:night-time ratio and a fixed RH_max_ of 0.9 are shown in Figure 5B and will be analysed further.

**Fig 5:**
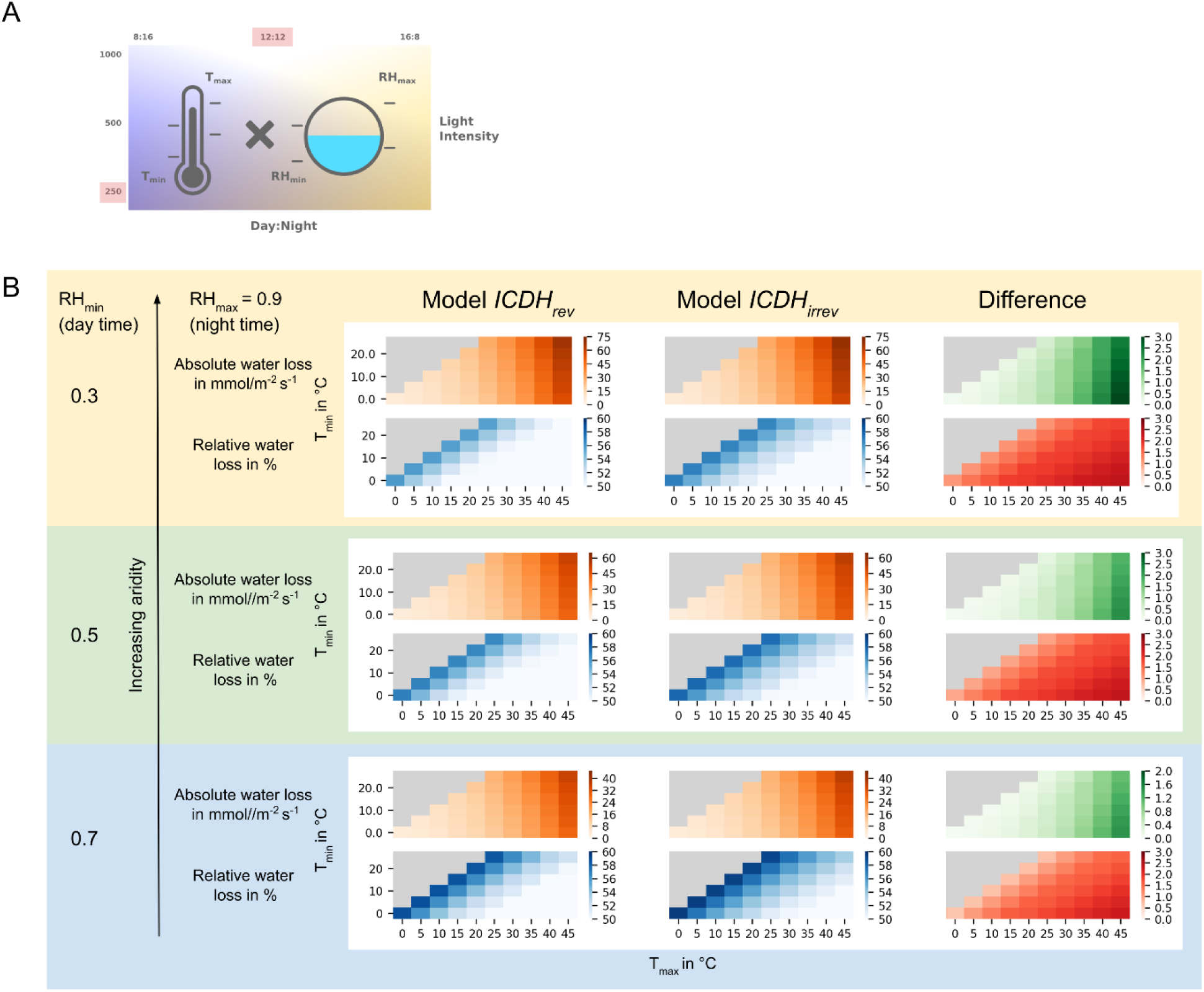
Absolute and relative water loss of a CAM-like leaf at 70% productivity with respect to a C_3_ leaf for model *ICDH*_*rev*_ and model *ICDH*_*irrev*_ for different temperature T and relative humidity RH regimes. (A) Overview of the environments analysed. (B) Simulation results are structured as follows: All simulations are performed for the same RH_min_ of 0.3 and three different values of RH_max_ (0.5, 0.7, 0.9) as indicated by the three blocks with different background colours. Within each block two output values are shown for different combinations of T_min_ and T_max_ values represented by the heatmaps: The upper row shows the absolute water loss for model *ICDH*_*rev*_ (orange heatmap, left) and model *ICDH*_*irrev*_ (orange heatmap, right) and the difference between model *ICDH*_*irrev*_ and *ICDH*_*rev*_ (green heatmap). The lower row shows the water loss in % with respect to the C_3_ model, again for model *ICDH*_*rev*_ (blue heatmap, left) and model *ICDH*_*irrev*_ (blue heatmap, right) and the difference (red heatmap). Darker colours represent a higher value. The results presented here were calculated for a maximum light intensity of 250 μmol m^-2^ s^-1^ and a 12:12 day:night cycle. All other results are presented and discussed in the Supplementary Text.

#### Absolute water loss is the highest for high T and low RH values

The model predicted the highest absolute water loss for low RH and high T. This behaviour was to be expected from the gas-diffusion relationship between the system’s demand for CO_2_ and the resulting water loss through transpiration. In Figure 5B this relationship is illustrated by the orange heatmaps, which represent the absolute water loss experienced by both models — *ICDH*_*rev*_ (left) and *ICDH*_*irrev*_ (middle) for all analysed combinations of T_min_ (y-axis) and T_max_ (x-axis) values for three different RH_min_ values (0.3, 0.5, 0.7). The darker the colours the higher absolute water loss. For example, at RH between 0.5 and 0.9, T_min_ = 10°C and T_max_ between 20 and 30°C water loss increased by 60 and 61%, respectively. Furthermore, we observed that the day-time temperature T_max_ was the main driver for water loss as the colour gradient changed more along the x-axis than along the y-axis. This could be explained by the occurrence of phase II and IV of the CAM cycle in our model. During these two phases the stomata opened during the day and water-loss was much higher than at night in phase I. This was also reflected in the effect of relative humidity on water loss. We found that the effect of the day-time humidity RH_min_ on the absolute water loss was stronger than the effect of RH_max_ (A comparative plot for changing RH_max_ is shown in the Supplementary Text, Figure 2). For example, water loss was 1.9-fold higher at RH_min_ = 0.3 as compared to RH_min_ = 0.7 at T_min_ = 10°C, T_max_ = 30°C and RH_max_= 0.9. The difference in the absolute water loss between model *ICDH*_*rev*_ and model *ICDH*_*irrev*_ was largest for combinations of low T_min_ and high T_max_ (darkest areas in the green heatmap in Figure 5B) and for the highest ΔRH value, i.e. RH_min_ = 0.3 and RH_max_ = 0.9. Therefore, with respect to absolute water-saving, model *ICDH*_*rev*_ outperformed model *ICDH*_*irrev*_ the most for high day- and low night-time temperatures and large RH differences between day and night.

#### Relative water loss is the highest for small diel temperature (ΔT) and high RH changes (ΔRH)

Next, we considered the relative water loss with respect to the C_3_ leaf which is illustrated in the blue heatmaps (Figure 5B). Here, we found that the relative water loss was the lowest for combinations of low night-time temperatures and high day-time temperatures. This observation could be explained by the fact that low night-time temperatures benefited the water-saving for a CAM-like leaf but had little impact on the water-loss of a C_3_ leaf which closed the stomata during the night. Conversely, this holds true for high day-time temperatures. We also observed that high ΔRH values drove high water loss. Finally, the red heatmaps show the difference in relative water loss with respect to the C_3_ model between model *ICDH*_*rev*_ and *ICDH*_*irrev*_. The plot reveals differences in the contribution of ICDH_rev_ to the water-saving potential of the CAM-cycle. As with the absolute water loss, the contribution was the highest for environments with high day-time and low night-time temperatures and environments with low RH values. However, in contrast to the difference in absolute water-saving, the difference in relative water-saving was high across a large range of conditions (as can be seen by the dark colours for most parameter combinations). This indicated that for most of the encountered conditions ICDH_rev_ can have a significant contribution to relative water-saving with respect to a C_3_ plant.

#### Day length and light intensity impact water loss

In addition to T and RH, we investigated the influence of different light intensities and day-light hours on water loss. We found a strong effect of light intensity on the additional relative water-saving potential of running the isocitrate-citrate-proline-2OG cycle and we observed that for higher light intensities this effect diminishes. Changing day-time hours also had an effect on the relative water-saving potential of ICDH_rev_ and can make a difference of up to 4% for short days where it only reached 2% for long days. Plots showing the water loss as dependent on light intensity and day length are shown in Supplementary Figure 3 and 4, respectively.

## Discussion

The time-resolved, environment-coupled model of leaf metabolism allowed us to study the trade-offs between productivity and water-saving for different network configurations and across different environmental conditions in a systematic manner. Our analysis led to three main conclusions. First, the leaf’s vacuolar storage capacity is a major determinant of the extent of a CAM cycle and without engineering a higher vacuole to cytoplasm ratio it will be unlikely that a full CAM cycle can be engineered into a C_3_ leaf. Secondly, the reversibility of mitochondrial ICDH might contribute to initial carbon fixation at night-time. This operational mode of the TCA cycle was previously demonstrated by metabolic flux analysis in rapeseed and soybean embryos ^23,24^ but is a novel prediction with respect to nocturnal CO_2_ assimilation. Thirdly, the water-saving effect of CAM strongly depends on the environment and the additional water-saving effect of carbon fixation by ICDH can range between 0.1 and 4% for the environmental conditions tested here. The additional water-saving contribution is largest at lower light intensities and for broad ranges of temperature and relative humidity which makes it an interesting candidate for metabolic engineering approaches as these should be beneficial for many weather conditions typically encountered by C_3_ crops.

### Reduced photorespiration due to daytime stomata closure can increase the water-saving potential of CAM leaves

Our previous study on CAM photosynthesis investigated the energetics and productivity of metabolic networks operating in C_3_ and CAM. It was found that — depending on the rates of the carboxylase and oxygenase activities of RuBisCO — the productivity of a CAM network could reach between 74 - 100% of the C_3_ network ^13^. In the analysis presented here, we focused on the water-saving potential of CAM without considering the potentially positive effect of carbon concentration behind closed stomata during the day. As we do not know how the carboxylation to oxygenation ratio changes as we move along the Pareto frontier from open stomata to partial and full closure during the daytime we used a constant value of 3:1. Therefore the implications of our analysis can be regarded as a conservative estimate. Due to the suppression of photorespiration in a leaf operating in CAM mode the actual water-saving potential at the same productivity level is expected to be higher than calculated here.

### ICDH might play a role in facultative CAM photosynthesis

Diel cycles of Pro accumulation have been previously observed in ice plant exposed to CAM-inducing salt stress. Under this stress condition Pro is known to act as an osmoprotectant. It has been reported that Pro accumulation proceeded in an oscillating manner in which high levels of Pro accumulated during the day (up to 16 μmol gFW^-1^), followed by a partially degradation at night which led to steadily increasing Pro levels during the CAM-induction phase ^30^. The increase in Pro levels was accompanied by an increase in PEPCase mRNA up to 10 days after stress exposure when PEPC mRNA has reached a full CAM level. This oscillatory behaviour led the authors to the following statement “Changes of proline in light and darkness suggested that proline plays an important role in addition to serving as an osmolyte.” but they offered no further explanation of what this role could be. We suggest that in addition to its function as an osmoprotectant during the day, Pro degradation at night might support C-fixation by supplying the substrate 2OG for citrate synthesis through ICDH in the mitochondria. Once PEPC capacity has been induced to the level required for full CAM the initial CO_2_-fixation proceeds via this enzyme, as it is kinetically superior, catalysing a thermodynamically favourable reaction compared to *ICDH*_*rev*_ (ΔG = −40kJ mol^-1^ at pH 7, ionic strength of 0.1 M, and reactant concentrations of 1 mM ^27^).

This conclusion is further supported by another study in ice plant in which malate, citrate and isocitrate levels and CAM-relevant enzyme activities were measured for the same CAM-inducing conditions ^5^. The authors reported malate and citrate levels of up to 27.5 mM and 29.4mM, respectively. Isocitrate levels were about 10 times lower than citrate levels. These observations are in line with our model’s predictions of carboxylic acid storage. Moreover, in accordance with previous reports ^31^ the authors found citrate accumulation to precede malate accumulation throughout the CAM induction period. Enzymes essays revealed an increased night-time activity of mitochondrial citrate synthase towards the end of the induction period. These findings support the prediction that (iso-)citrate synthesis through citrate synthase increases as PEPC concentrations reach full CAM level.

### Considerations of enzyme cost could improve flux mode predictions

Our analysis explored the water-saving potential of alternative flux routes without accounting for the enzymatic cost of different flux routes. Previous modelling approaches in bacteria and cyanobacteria have successfully included a cost for enzyme synthesis in the form of a ‘resource allocation’ or ‘resource balance’ problem ^32–35^. However, these or similar modelling approaches rely on comprehensive knowledge of *k*_*cat*_ and enzyme turnover rates or large datasets to estimate such parameters and therefore are still of limited applicability for large-scale plant metabolic models.

### Implications for engineering CAM into a C_3_ species in temperate climates

Strategies for introducing CAM into crop plants to make them more resilient to hotter and drier conditions have been largely discussed in the context of arid or marginal lands ^36–38^; ^236–38^. Less attention has been given to the question of whether CAM could benefit the productivity of C_3_ species typically grown in temperate climates, such as wheat or barley, where hot and dry periods are becoming increasingly frequent ^39,40^. In this context a flexible CAM i.e. a ‘C3+CAM’ phenotype could be beneficial as it combines high productivity in C_3_ mode with increased WUE in CAM mode. Besides the challenge of engineering a CAM cycle into a C_3_ crop, other factors such as the trade-off between improved WUE and potential constraints on leaf productivity due to anatomical changes or the response of CAM to increasing CO_2_ levels need consideration ^41^. However, naturally occurring CAM has two characteristics that make it a suitable target for engineering approaches for crops grown in temperate regions. First, the CAM syndrome is extremely flexible. It has been shown that the contribution of CAM to diel CO_2_ uptake patterns can range between 0 and 100%, particularly in plants with either ontogenetically or environmentally induced transition from C_3_ to CAM ^41^. Secondly, CAM has evolved many times independently and it is believed to be present in well over 5% of vascular plant species ^42,43^. These observations can be attributed to the fact that CAM most likely evolved on a ‘biochemistry first, anatomy second’ trajectory ^44^ in which ‘C_3_+CAM’ is an evolutionarily accessible phenotype on the trajectory to strong CAM ^45,46^. Our model demonstrates the water-saving potential of a partial CAM or CAM-like mode for plants grown in temperate climates, while maintaining a high net metabolic output. In particular, we predict that the relative additional water-saving of operating an isocitrate-citrate-proline-2OG nocturnal carbon fixation cycle is significant across a wide range of conditions. To summarize, our model supports the potential productivity benefits of flexible CAM as it can enable temperate crops plant to better cope with and survive periods of heat and drought with only minimal impact upon metabolic productivity.

### Overcoming vacuolar storage constraints by increasing cell size

Our study identified the vacuolar storage capacity as a limiting factor for introducing CAM into a C_3_ plant. Comparison of models with different vacuolar storage capacities revealed that shifting from C_3_ to CAM leaf anatomy increased water-saving by 6.6 %, given a 3.1-fold increase in the vacuolar storage capacity. Given this observation, engineering leaf anatomical parameters of a C_3_ leaf towards CAM architecture will be key to increasing water-saving. How could this be achieved? Several anatomical traits (cell size (as a proxy for vacuole size), % intercellular airspace (IAS), tissue thickness) have been discussed in this context ^44^. However, altering the latter two parameters could potentially limit photosynthesis in C_3_ mode due to reduced CO_2_ diffusion through the mesophyll and a lower *C*_*i*_ ^47^ and therefore disadvantage flexible CAM. Barrera Zambrano et al. proposed that Clusia might overcome this discrepancy by having a high % IAS in the spongy mesophyll for efficient CO_2_ diffusion in C_3_ mode, and large palisade cells for carboxylic acid storage in CAM mode and suggested this as a potential engineering strategy for transferring inducible CAM into a C_3_ plant ^48^. Of particular relevance is a study which attempted to increase leaf cell size by overexpressing a grape berry transcription factor (*VvCEB1*_opt_) in *Arabidopsis thaliana* and *Nicotiana sylvestris* ^49^. This approach increased cell size — but not number — in both species with a 1.8-to 2.3-fold increase in the palisade mesophyll and 2.0-to 2.5-fold increase in the spongy mesophyll in *A. thaliana.* Assuming that the larger cell size is mainly driven by increased vacuolar volume the reported increase would suffice to enable partial CAM with high water-saving potential. Besides increased cell size, the authors also reported a significant decrease in cell wall thickness — another feature typically observed in CAM plants ^36^. Thus, (tissue-specifically) overexpressing *VvCEB1*_op_ in the context of engineering CAM could indeed be a promising strategy to engineer more drought-resistant crops.

## Materials and Methods

### Developing a time-resolved environment-coupled model of leaf metabolism

The model building process was divided into two parts: developing a time-resolved model of leaf metabolism and modelling the gas-exchange through the stomata. The time-resolved diel model is an extension of our previously published diel modelling framework ^13,50^. Starting from the latest version of a charge and proton balanced generic core model of plant metabolism (PlantCoreMetabolism_v1_2_3.xml) we concatenated 24 copies (each representing one hour of the day) of the model by allowing a range of metabolites to accumulate and to be transferred from one time-point to the other via so-called *‘linker fluxes’*. Starch was allowed to accumulate freely in the plastid. The sugars glucose, sucrose, and fructose, the carboxylic acids malate, citrate and isocitrate, the proteinogenic amino acids, and nitrate were allowed to accumulate in the vacuole. The last time interval of the optimization routine was coupled to the first interval to form a closed diel cycle. The overall vacuolar storage capacity was based on estimates for the vacuolar malate storage capacity and the vacuolar volume of an average C_3_ plant (see Supplementary Text for detailed calculations). The specificity of the metabolic network at each hour of the day was achieved by setting the light input according to the diel light curve and by constraining the export of sugars and amino acids to the phloem (phloem output) to a day:night ratio of 3:1 and nitrogen uptake to a day:night ratio of 3:2 according to previous estimates ^50^. Maintenance cost was modelled in a light-dependent manner, where the day-time cost depends on the average daytime light intensity (Supplementary Text). The day:night ratio was assumed to be 3:1 and the ratio of ATP maintenance cost to NADPH maintenance cost was assumed to be 3:1 (Cheung et al., 2013). RuBisCO was only activated during day-light hours. Due to the lack of knowledge about the RuBisCO carboxylation:oxygenation ratio we used a value of 3:1 as estimated form flux measurements in Arabidopsis ^51^. The uptake rate for CO_2_ was limited to a value of 15 μmol m^-2^ s^-1^ based on values for different C_3_ and CAM species at a light intensity of 250 μmol m^-2^ s^-1^ ^52^. All other fluxes were unconstrained. To avoid any bias, we used both a phloem composition of a C_3_ crop plant (tomato, Arabidopsis) and a CAM species (Opuntia) and found no qualitative differences in our analysis. The phloem composition and output values for the Opuntia and Arabidopsis data-constrained model are listed in the Supplementary Table 2.

Gas-exchange through the stomata was described by a linearized diffusion model which predicts the water-loss depending on the metabolic model’s demand for CO_2_, at a particular T and RH. The input curves for T and RH are based on a normal distribution function and allowed us to systematically scan a multidimensional parameter space by adjusting the function parameters accordingly.

The model equations for optimizing phloem output and water-saving were solved as a linear optimization problem (L_1_ norm). The subsequent minimization of the metabolic flux sum was solved as a quadratic optimization problem (L_2_ norm) to select from a possible set of multiple solutions, the one with the least variation in fluxes between time points. To exemplify this, consider a three-time-step model with the flux sequence [1, 1, 1], [2, 0, 1] and [3, 0, 0]. When applying the L_1_ measure all three cases will be weighted with 3 although in the second and third case more enzyme needs to be synthesized and degraded and therefore would be costlier. The L_2_ distance yields values of 3, 5, and 9 and would therefore prefer the flux distribution in which fluxes are equally split between the 3 phases. A derivation of the gas-water exchange relationship through the stomata, any further modelling assumptions, parameter derivations, and auxiliary calculations are detailed in the Supplementary Text.

## Supplementary Information

Supplementary Text: Supplementary methods and results

Supplementary Table 1: Leaf parameters for different C_3_ and CAM species

Supplementary Table 2: Phloem sap compositions for different species and modelling results for different species and constraints.

## Abbreviations

Asn: asparagine
Asp: asparagine
PEP: phosphoenolpyruvate
Pro: proline
RuBisCO: ribulose-1,5-bisphosphate carboxylase/oxygenase
2OG: 2-oxoglutarate

