## Supplemental Text for "CAM emerges in a leaf metabolic model under water-saving constraints in different environments"

### Supplementary Methods and Results: CAM emerges in a leaf metabolic model under water-saving constraints in different environments

Nadine Töpfer, Thomas Braam, Sanu Shameer,
R. George Ratcliffe, Lee J. Sweetlove

#### 1 Deriving of a simplified gas-diffusion equation

To model gas-exchange through the stomata in dependence of the metabolic system's demand for $CO_2$ we derived a simplified form of the gas-diffusion equation presented in [1] which is only dependent on the temperature ($T$) and relative humidity ($RH$) of the environment. As a starting point, we considered the one-dimensional form of Fick's first law of diffusion, Eq.8.4b on page 374 of the book, which describes the diffusion flux across a stomata pore:

$$J_j = D_j \cdot \frac{\Delta c_j^{\mathrm{st}}}{\Delta x^{\mathrm{st}}} \cdot \frac{A^{\mathrm{st}}}{A} = D_j \cdot \frac{n \cdot a^{\mathrm{st}}}{\delta x^{\mathrm{st}}} \cdot \Delta c_j^{\mathrm{st}}, \tag{1}$$

where $J_j$ is the flux density (short flux) of compound $j$ through the stomata pore, $D_j$ is the diffusion coefficient of compound $j$, $\Delta c_j^{\mathrm{st}}$ is the concentration gradient of compound $j$ and $\Delta x^{\mathrm{st}}$ is distance across the pore. $A^{\mathrm{st}}$ is the area occupied by open stomata pores and $A$ the total leaf area under consideration. Moreover, $n$ is the number of stomata per unit area of the leaf and $a^{\mathrm{st}}$ is the average area per stomata pore. Thus $na^{\mathrm{st}}$ equals the fraction of the leaf surface area occupied by stomata pores. Assuming that open stomata have a circular shape, we obtain:

$$J_j = D_j \cdot \frac{n \cdot \pi \cdot R^2}{l + R_{\mathrm{mean}}} \cdot (c_{\mathrm{in},j} - c_{\mathrm{out},j}), \tag{2}$$

where $\pi \cdot R^2$ is the approximate area of the stomata opening, $l$ is the actual depth of the pore, $R_{\mathrm{mean}}$ is a correction factor, about the mean "radius" of the pore , which accounts for the fact that the effective depth of the pore is greater than the actual pore depth [1]. The values $c_{\mathrm{in},j}$ and $c_{\mathrm{out},j}$ denote the inside and outside concentration of $j$, respectively. Here, we assume $R_{\mathrm{mean}}$ to be constant and assign it a value of $5 \cdot 10^{-6}$ m [1] (This value will only be important if we want to calculate the stomata radius).

For the diffusion flux of $CO_2$ across the stomata pore this equations reads as follows:

$$J_{\mathrm{CO_2}} = D_{\mathrm{CO_2}} \cdot \frac{n \cdot \pi \cdot R^2}{l + R_{\mathrm{mean}}} \cdot (c_{\mathrm{in,CO_2}} - c_{\mathrm{out,CO_2}}), \tag{3}$$

and for $H_2O$:

$$J_{\mathrm{H_2O}} = D_{\mathrm{H_2O}} \cdot \frac{n \cdot \pi \cdot R^2}{l + R_{\mathrm{mean}}} \cdot (c_{\mathrm{in,H_2O}} - c_{\mathrm{out,H_2O}}), \tag{4}$$

Solving Eq.3 for $R$ results in:

$$R = \sqrt{\frac{J_{\mathrm{CO_2}} \cdot (l + R_{\mathrm{mean}})}{D_{\mathrm{CO_2}} \cdot n \cdot \pi \cdot (c_{\mathrm{in,CO_2}} - c_{\mathrm{out,CO_2}})}}. \tag{5}$$

Substituting $R$ in Eq.4 yields:

$$J_{\mathrm{H_2O}} = \frac{D_{\mathrm{H_2O}} \cdot n \cdot \pi \cdot (l + R_{\mathrm{mean}}) \cdot (c_{\mathrm{in,H_2O}} - c_{\mathrm{out,H_2O}})}{D_{\mathrm{CO_2}} \cdot n \cdot \pi \cdot (l + R_{\mathrm{mean}}) \cdot (c_{\mathrm{in,CO_2}} - c_{\mathrm{out,CO_2}})} \cdot J_{\mathrm{CO_2}} \tag{6}$$

or

$$J_{\mathrm{H_2O}} = \frac{D_{\mathrm{H_2O}} \cdot (c_{\mathrm{in,H_2O}} - c_{\mathrm{out,H_2O}})}{D_{\mathrm{CO_2}} \cdot (c_{\mathrm{in,CO_2}} - c_{\mathrm{out,CO_2}})} \cdot J_{\mathrm{CO_2}}. \tag{7}$$

According to Eq.8.9 on page 379 of [1] the diffusion coefficient $D_j$ is a function of the temperature $T$, with:

$$D_j \cong D_j^0 \cdot (T/273)^{1.8}, \tag{8}$$

where $D_j^0$ is the standard diffusion coefficient. This cancels out in Eq.7 to:

$$J_{\mathrm{H_2O}} = \frac{D^0_{\mathrm{H_2O}} \cdot (c_{\mathrm{in,H_2O}} - c_{\mathrm{out,H_2O}})}{D^0_{\mathrm{CO_2}} \cdot (c_{\mathrm{in,CO_2}} - c_{\mathrm{out,CO_2}})} \cdot J_{\mathrm{CO_2}}, \tag{9}$$

According to [1], we can calculate $c_{\mathrm{H_2O}}$ at a given temperature $T$ by multiplying the saturation concentration of water vapour, $c^*_{\mathrm{H_2O}}$ by the relative humidity $RH$ (as a decimal) at this temperature. The values for $c^*_{\mathrm{H_2O}}$ for temperatures from -30° to +60° are listed in Appendix I, pages 552-554 in [1]. Thus, we can rewrite Eq.9:

$$J_{\mathrm{H_2O}} = \frac{D^0_{\mathrm{H_2O}}}{D^0_{\mathrm{CO_2}}} \cdot \frac{(c^*_{\mathrm{in,H_2O}} \cdot RH_{\mathrm{in}} - c^*_{\mathrm{out,H_2O}} \cdot RH_{\mathrm{out}})}{(c_{\mathrm{in,CO_2}} - c_{\mathrm{out,CO_2}})} \cdot J_{\mathrm{CO_2}}. \tag{10}$$

We assume $RH_{\mathrm{in}}$ to be 100% and can approximate $c_{\mathrm{in,CO_2}}$ and $c_{\mathrm{out,CO_2}}$ from *e.g.* [2]. This leads to:

$$J_{\mathrm{H_2O}} = K \cdot (c(T)^*_{\mathrm{in,H_2O}} - c(T)^*_{\mathrm{out,H_2O}} \cdot RH_{\mathrm{out}}) \cdot J_{\mathrm{CO_2}}, \tag{11}$$

with:

$$K = \frac{D^0_{\mathrm{H_2O}}}{D^0_{\mathrm{CO_2}}} \cdot \frac{1}{(c_{\mathrm{in,CO_2}} - c_{\mathrm{out,CO_2}})}. \tag{12}$$

This demonstrates that water loss through the stomata can be approximated by the system's demand for $CO_2$, $T$ and $RH$, only. Leaf parameters such as the number of stomata per area, stomata size, and the actual and effective pore depth are not necessary. Finally, looking at the standard units of Eq.1:

$$J_j[\frac{\mathrm{mol}}{\mathrm{m^2 \cdot s}}] = D_j[\frac{\mathrm{m^2}}{\mathrm{s}}] \cdot \Delta c_j[\frac{\mathrm{mol}}{\mathrm{m^3}}] \cdot \frac{1}{\Delta x_j}[\frac{1}{\mathrm{m}}] \cdot \frac{A^{st}}{A}[\frac{\mathrm{m^2}}{\mathrm{m^2}}] \tag{13}$$

$$[\frac{\mathrm{mol}}{\mathrm{m^2 \cdot s}}] = [\frac{\mathrm{m^2}}{\mathrm{s}}] \cdot [\frac{\mathrm{mol}}{\mathrm{m^3}}] \cdot [\frac{1}{\mathrm{m}}] \cdot [\frac{\cancel{m^2}}{\cancel{m^2}}]; \tag{14}$$

we find that $\Delta c_j$ should be given in mol/m$^3$.

### 2 Model construction

#### 2.1 Light, temperature, and relative humidity input

The model can be either analyzed for generated or measured light, temperature, and humidity data.

**Generated data:** Light, $T$, and $RH$ curves can be generated using a Gaussian function with parameters for peak height, width, shift, and base level (e.g., file 'Temperature-and-Humidity-Curves.csv'). Based on a given day length and a maximum light intensity the respective Gaussian curve is pre-calculated and used as input for the model. For reasons of comparability the light zenith is fixed to phase 7 in the 24 phase model.

**Measured temperature and relative humidity data:** Alternatively, the model can be analysed using measured temperature and humidity data and a given maximum light intensity that is balanced to take a bell shaped curve around the zenith at phase 7 and simulates a 12/12 day-night cycle.

#### 2.2 Maintenance cost (ATP and NADPH consumption)

Maintenance costs in a leaf are dependent on the incident light intensity. To compute an appropriate value for ATP maintenance cost for the light intensity used in this study we collected experimental data for dark respiration and starch content for $C_3$ leaves at different light intensities [3, 4, 5, 6]. We then set up a two phase diel leaf model [7] and varied the value of the ATPase reaction in the model representing ATP maintenance costs until, in each case, the model predicted values for starch and/or dark respiration that matched the experimental data. This yielded a linear relationship between ATP maintenance costs ($v_{\text{ATPase}}$ in $\mu$mol m$^{-2}s^{-1}$) and light intensity $\mu$mol m$^{-2}s^{-1}$):

$$v_{\text{ATPase}} = 0.0049 \cdot \overline{\text{light intensity}} + 2.7851 \tag{15}$$

Based on this value the system's maintenance demand for reducing power, represented by NADPH oxidases, is set to a $v_{\text{ATPase}}$:$v_{\text{NADPHoxidases}}$ ratio of 3:1 [8].

#### 2.3 Malate storage capacity of the vacuole

To include the average vacuolar storage capacity of a $C_3$ or CAM leaf in our model we need to link concentrations to fluxes. In our model the vacuolar amount of metabolite $X$ per m$^2$ at a given time-interval (1 hour) equals to the linker flux of metabolite $X$ in $\mu$mol m$^{-2}$s$^{-1}$ multiplied by 3600 (conversion from second to hour). This relationship can be used to define an upper limit for the sum of all vacuolar linker fluxes at a given time-interval. As most CAM species accumulate large amount of malate we use the vacuolar malate storage capacity $C_{\text{Mal}}$ as a proxy for the total vacuolar storage capacity. This value is assumed to be 200 mol m$^{-3}$ (200 mM) based on measurements in *K. daigremontiana* [9]. Additionally, we need to know the vacuolar volume of 1 m$^2$ leaf which can be calculated from values for the mesophyll thickness, the porosity of the leaf, and the vacuolar volume for several $C_3$ and CAM species as given in [10]. From these data we can calculate the maximum amount of malate that can be transferred by the the liker flux (units in $\mu$mol m$^{-2}$s$^{-1}$).

Vacuolar Volume $V_V$ per m$^2$ leaf:

$$V_V = \text{Mesophyll thickness} \cdot (1 - \text{Porosity}) \cdot \text{Relative vacuolar volume} \tag{16}$$

Relative vacuolar volume (fraction of cell that is vacuole) = 0.95 for CAM species and 0.9 for $C_3$ species [11, 12].

Amount of malate $MAL$ per m$^2$ leaf:

$$MAL = c_{Mal} \cdot V_V. \tag{17}$$

This amount is transferred to the next hour interval at a constant linker flux $v_{\text{L,Mal}}$ and we obtain:

$$v_{\text{L,Mal,max}} = MAL/3600 \tag{18}$$

or

$$v_{\text{L,Mal,max}} = c_{\text{Mal}} \cdot \text{Mesophyll thickness} \cdot (1 - \text{Porosity}) \cdot \frac{\text{Relative vacuolar volume}}{3600} \tag{19}$$

For the values listed in the Supplementary Table 1 we calculate the maximum linker flux:

- for an average $C_3$ leaf $v_{\text{L,Mal,max,C}_3\text{,average}} = 7.68$ $\mu$mol m$^{-2}$s$^{-1}$
- for an average CAM leaf $v_{\text{L,Mal,max,CAM,average}} = 24.04$ $\mu$mol m$^{-2}$s$^{-1}$

We assume that only green mesophyll, has malate storage capacity that contributes to CAM [13] as the transport to non-green water-tissue is considered to be too slow and ineffective for CAM [14].

#### 2.4 Concentration gradient between the leaf and the environment

We assume an average $c_i : c_a$ ratio of 0.75 which is a typical mean ratio for $C_3$ plants under field conditions at ambient $CO_2$ concentrations ($360\mu$mol mol air$^{-1}$) [15], where $c_i$ is the leaf internal and $c_a$ is the external concentration of $CO_2$. This leads to an average $c_i$ of $270\mu$ mol molair$^{-1}$ and $\Delta CO_2$ of $90\mu$ mol molair$^{-1}$ or 0.004 mol m$^{-3}$ air.

$\Delta CO_2 = 0.004$

#### 2.5 Diffusion coefficients for water and $CO_2$

In air at 1 atm and 0° Celsius (as in Appendix I in [1]):
Following Eq 8.9 and Appendix I in [1] the temperature-dependent diffusion coefficient under atmospheric pressure of 1 atm can be approximated by:

$$D_j = D_{j,0} \cdot T/273 \cdot 10^{1.8}, \tag{20}$$

with

$D_{\mathrm{H2O}_0} = 2.13 \cdot 10^{-5} \mathrm{m}^2\ \mathrm{s}^{-1}$,

$D_{\mathrm{CO2}_0} = 1.33 \cdot 10^{-5} \mathrm{m}^2\ \mathrm{s}^{-1}$.

#### 2.6 Temperature difference between leaf outside and inside - $\Delta$ T

We assume a value of 2 degrees which is a typical value from the literature (De Boeck et al. 2012).

$\Delta T = 2°\mathrm{C}$

#### 2.7 Dark-light threshold approximated by light-compensation point

When using an artificial normal distributed light curves we need to define a cutoff value below which the given light intensity is considered as dark. To approximate this value we use the light-compensation point measured for *Kalanchoe daigremontiana*[16].

Light-compensation point $= 30\mu$mol m$^{-2}$s$^{-1}$

### 3 Modelling results

#### 3.1 Sequence of reactions involved in C-fixation pathways

It shall be noted that the reaction sequences here only describe the major flux routes observed in our model. Depending one the exact hour of day these flux routes might be supported by metabolites from other pathways or they might branch off into other pathways.

```
#############################################
# Starch degradation Pathway to PEP
#############################################

RXN_1826_p Pi_p + STARCH_p --> GLC_1_P_p

PHOSPHOGLUCMUT_RXN_p GLC_1_P_p <=> GLC_6_P_p

G6P_Pi_pc GLC_6_P_p + 0.7 Pi_c + 0.3 aPi_c <=> GLC_6_P_c + 0.3 PROTON_c + Pi_p
```

```
GLUCISOM_RXN_c GLC_6_P_c <=> FRUCTOSE_6P_c

6PFRUCTPHOS_RXN_c 0.65 ATP_c + FRUCTOSE_6P_c + 0.35 aATP_c --> 0.5 ADP_c +
FRUCTOSE_16_DIPHOSPHATE_c + 0.85 PROTON_c + 0.5 aADP_c

F16ALDOLASE_RXN_c FRUCTOSE_16_DIPHOSPHATE_c <=> DIHYDROXY_ACETONE_PHOSPHATE_c +
GAP_c

GAPOXNPHOSPHN_RXN_c GAP_c + NAD_c + 0.7 Pi_c + 0.3 aPi_c <=> DPG_c + NADH_c +
1.3 PROTON_c

PHOSGLYPHOS_RXN_c 0.65 ATP_c + G3P_c + 0.15 PROTON_c + 0.35 aATP_c <=> 0.5 ADP_c
+ DPG_c + 0.5 aADP_c

3PGAREARR_RXN_c G3P_c <=> 2_PG_c

2PGADEHYDRAT_RXN_c  2_PG_c <=> PHOSPHO_ENOL_PYRUVATE_c + WATER_c

############################################
# Reactions involved in PEPC cycle at night
############################################

### starting from PEP and CO2
##
#

RXN0_5224_c CARBON_DIOXIDE_c + WATER_c <=> HCO3_c + PROTON_c

PEPCARBOX_RXN_c HCO3_c + PHOSPHO_ENOL_PYRUVATE_c + 0.3 PROTON_c -->
OXALACETIC_ACID_c + 0.7 Pi_c + 0.3 aPi_c

### part of the OAA is converted to MAL and stored in the vacuole
##
#

MALATE_DEH_RXN_c MAL_c + NAD_c <=> NADH_c + OXALACETIC_ACID_c + PROTON_c

MAL_PROTON_vc MAL_c + 0.3 PROTON_v --> 0.7 MAL_v + 0.3 aMAL_v

### part of the OAA is involved in a MAL/OAA cycle between cytosol and mitochondria
or converted to CIT in the mitochondria and then exported for storage
##
#

OAA_MAL_mc MAL_c + OXALACETIC_ACID_m <=> MAL_m + OXALACETIC_ACID_c

MALATE_DEH_RXN_m MAL_m + NAD_m <=> NADH_m + OXALACETIC_ACID_m + PROTON_m

# or

CITSYN_RXN_m ACETYL_COA_m + OXALACETIC_ACID_m + WATER_m --> CIT_m + CO_A_m +
PROTON_m

### ACETYL-COA from PEP for CIT synthesis with OAA
```

```
##
#

PEPDEPHOS_RXN_c 0.5 ADP_c + PHOSPHO_ENOL_PYRUVATE_c + 0.85 PROTON_c +
0.5 aADP_c --> 0.65 ATP_c + PYRUVATE_c + 0.35 aATP_c

PYRUVATE_PROTON_mc PROTON_m + PYRUVATE_m <=> PROTON_c + PYRUVATE_c

PYRUVDEH_RXN_m CO_A_m + NAD_m + PYRUVATE_m --> ACETYL_COA_m + CARBON_DIOXIDE_m +
NADH_m

### Transporter
##
#

MAL_CIT_mc_01 CIT_c + MAL_m + PROTON_c <=> CIT_m + MAL_c + PROTON_m

CIT_PROTON_vc CIT_c + 0.5 PROTON_v --> 0.5 CIT_v + 0.5 aCIT_v


##################################################
# Reactions involved in PEPC cycle during the day
##################################################

#################
# First conversion
#################

### MAL to CIT conversion at start of day
##
#

MAL_PROTON_rev_vc 0.7 MAL_v + 1.7 PROTON_v + 0.3 aMAL_v --> MAL_c + 2.0 PROTON_c

MAL_xc MAL_x <=> MAL_c

MALATE_DEH_RXN_x MAL_x + NAD_x <=> NADH_x + OXALACETIC_ACID_x + PROTON_x

CITSYN_RXN_x ACETYL_COA_x + OXALACETIC_ACID_x + WATER_x --> CIT_x + CO_A_x +
PROTON_x

### CIT is exported back to the vacuole and then takes the route described further below


### Acetyl-CoA from C-fixation by RuBisCO and PEPC via PEP and OAA
##
#

### PEP from C-fixation by RuBisCO


RIBULOSE_BISPHOSPHATE_CARBOXYLASE_RXN_p CARBON_DIOXIDE_p + D_RIBULOSE_15_P2_p +
WATER_p --> 2.0 G3P_p + 2.0 PROTON_p

3PGAREARR_RXN_p G3P_p <=> 2_PG_p
```

```
2PGADEHYDRAT_RXN_p 2_PG_p <=> PHOSPHO_ENOL_PYRUVATE_p + WATER_p

PEP_Pi_pc PHOSPHO_ENOL_PYRUVATE_p + 0.7 Pi_c + 0.3 aPi_c <=>
PHOSPHO_ENOL_PYRUVATE_c + 0.3 PROTON_c + Pi_p


#### PEP to Acetyl-CoA with additional C-fixation by PEPC

PEPCARBOX_RXN_c HCO3_c + PHOSPHO_ENOL_PYRUVATE_c + 0.3 PROTON_c --> OXALACETIC_ACID_c +
0.7 Pi_c + 0.3 aPi_c

ASPAMINOTRANS_RXN_c 2_KETOGLUTARATE_c + L_ASPARTATE_c <=> GLT_c + OXALACETIC_ACID_c

# Transport c to p

ASPARTATEKIN_RXN_p 0.9 ATP_p + L_ASPARTATE_p + 0.1 PROTON_p + 0.1 aATP_p -->
0.8 ADP_p + L_BETA_ASPARTYL_P_p + 0.2 aADP_p

ASPARTATE_SEMIALDEHYDE_DEHYDROGENASE_RXN_p L_BETA_ASPARTYL_P_p + NADPH_p +
PROTON_p --> L_ASPARTATE_SEMIALDEHYDE_p + NADP_p + Pi_p

HOMOSERDEHYDROG_RXN_NADP_p L_ASPARTATE_SEMIALDEHYDE_p +
NADPH_p + PROTON_p --> HOMO_SER_p + NADP_p

HOMOSERKIN_RXN_p 0.9 ATP_p + HOMO_SER_p + 0.1 aATP_p -->
0.8 ADP_p + O_PHOSPHO_L_HOMOSERINE_p + 0.9 PROTON_p + 0.2 aADP_p

THRESYN_RXN_p O_PHOSPHO_L_HOMOSERINE_p + WATER_p --> Pi_p + THR_p

# Transport p to c

THREONINE_ALDOLASE_RXN_c THR_c --> ACETALD_c + GLY_c

# Transport c to m

RXN66_3_m ACETALD_m + NAD_m + WATER_m --> ACET_m + NADH_m + 2.0 PROTON_m

# Transport m to x

ACETATE_COA_LIGASE_RXN_x ACET_x + 0.65 ATP_x + CO_A_x + 0.35 aATP_x -->
ACETYL_COA_x + AMP_x + 0.65 PPI_x + 0.35 aPPI_x

###################
# second conversion
###################

### CO2-release starting from CIT
##
#

CIT_PROTON_rev_vc 0.5 CIT_v + 2.5 PROTON_v + 0.5 aCIT_v --> CIT_c + 3.0 PROTON_c

ACONITATEDEHYDR_RXN_c CIT_c <=> CIS_ACONITATE_c + WATER_c
```

```
ACONITATEHYDR_RXN_c CIS_ACONITATE_c + WATER_c <=> THREO_DS_ISO_CITRATE_c

THREO_DS_ISO_CITRATE_xc THREO_DS_ISO_CITRATE_x <=> THREO_DS_ISO_CITRATE_c

ISOCIT_CLEAV_RXN_x THREO_DS_ISO_CITRATE_x --> GLYOX_x + SUC_x

GLYCINE_AMINOTRANSFERASE_RXN_x GLT_x + GLYOX_x <=> 2_KETOGLUTARATE_x + GLY_x

GLY_xc GLY_x <=> GLY_c

GLY_mc GLY_m <=> GLY_c

GCVMULTI_RXN_m GLY_m + NAD_m + THF_m --> AMMONIUM_m + CARBON_DIOXIDE_m + METHYLENE_THF_m +
NADH_m

and

GLYOHMETRANS_RXN_m SER_m + THF_m <=> GLY_m + METHYLENE_THF_m + WATER_m



#####################################################
# Reactions involved in the PEPC/ICDH cycle at night
#####################################################

### PEPC branch from PEP to ASN
##
#

RXN0_5224_c CARBON_DIOXIDE_c + WATER_c <=> HCO3_c + PROTON_c

PEPCARBOX_RXN_c HCO3_c + PHOSPHO_ENOL_PYRUVATE_c + 0.3 PROTON_c --> OXALACETIC_ACID_c +
0.7 Pi_c + 0.3 aPi_c

ASPAMINOTRANS_RXN_c 2_KETOGLUTARATE_c + L_ASPARTATE_c <=> GLT_c + OXALACETIC_ACID_c

ASNSYNA_RXN_c AMMONIUM_c + 0.65 ATP_c + L_ASPARTATE_c + 0.35 aATP_c --> AMP_c + ASN_c +
0.65 PPI_c + PROTON_c + 0.35 aPPI_c

ASN_PROTON_vc ASN_c + PROTON_v --> ASN_v + PROTON_c

### ICDH branch from PRO to CIT
##
#

PRO_PROTON_rev_vc PROTON_v + PRO_v --> PROTON_c + PRO_c

PRO_GLU_mc GLT_c + PRO_m <=> GLT_m + PRO_c

PRO_mc PRO_m <=> PRO_c

RXN_14903_mi PRO_m + UBIQUINONE_mi --> L_DELTA1_PYRROLINE_5_CARBOXYLATE_m + PROTON_m +
UBIQUINOL_mi

RXN_7183_NADP_m L_DELTA1_PYRROLINE_5_CARBOXYLATE_m + NADP_m + 2.0 WATER_m --> GLT_m +
```

```
NADPH_m + PROTON_m

GLUTAMATE_DEHYDROGENASE_RXN_m GLT_m + NAD_m + WATER_m --> 2_KETOGLUTARATE_m +
AMMONIUM_m + NADH_m + PROTON_m

ISOCITRATE_DEHYDROGENASE_NAD_RXN_m NAD_m + THREO_DS_ISO_CITRATE_m <-> 2_KETOGLUTARATE_m +
CARBON_DIOXIDE_m + NADH_m

ACONITATEHYDR_RXN_m CIS_ACONITATE_m + WATER_m <=> THREO_DS_ISO_CITRATE_m

ACONITATEDEHYDR_RXN_m CIT_m <=> CIS_ACONITATE_m + WATER_m

CIT_PROTON_vc CIT_c + 0.5 PROTON_v --> 0.5 CIT_v + 0.5 aCIT_v

### CIT transporters c to m
##
#

MAL_CIT_mc_01 CIT_c_01 + MAL_m_01 + PROTON_c_01 <=> CIT_m_01 + MAL_c_01 + PROTON_m_01

SUC_CIT_mc_01 CIT_c_01 + PROTON_c_01 + SUC_m_01 <=> CIT_m_01 + PROTON_m_01 + SUC_c_01

2KG_CIT_mc_01 2_KETOGLUTARATE_m_01 + CIT_c_01 + PROTON_c_01 <=> 2_KETOGLUTARATE_c_01 +
CIT_m_01 + PROTON_m_01

#######################################################
# Reactions involved in ICDH/PEPC cycle during the day
#######################################################

### starting from CIT to PRO
##
#

ACONITATEDEHYDR_RXN_c CIT_c <=> CIS_ACONITATE_c + WATER_c

ACONITATEHYDR_RXN_c CIS_ACONITATE_c + WATER_c <=> THREO_DS_ISO_CITRATE_c

ISOCITDEH_RXN_c NADP_c + THREO_DS_ISO_CITRATE_c <=> 2_KETOGLUTARATE_c + CARBON_DIOXIDE_c +
NADPH_c

2KG_MAL_pc 2_KETOGLUTARATE_p + MAL_c <=> 2_KETOGLUTARATE_c + MAL_p

GLUTAMATE_SYNTHASE_FERREDOXIN_RXN_p 2_KETOGLUTARATE_p + GLN_p + 2.0 PROTON_p +
2.0 Reduced_ferredoxins_p --> 2.0 GLT_p + 2.0 Oxidized_ferredoxins_p

####################
# a part goes here:
####################

GLUTAMINESYN_RXN_p AMMONIUM_p + 0.9 ATP_p + GLT_p + 0.1 aATP_p --> 0.8 ADP_p + GLN_p
+ 0.9 PROTON_p + Pi_p + 0.2 aADP_p

#############################
# and another part goes here:
#############################
```

```
GLT_MAL_pc GLT_p + MAL_c <=> GLT_c + MAL_p

GLUTKIN_RXN_c 0.65 ATP_c + GLT_c + 0.35 aATP_c --> 0.5 ADP_c + 0.91 L_GLUTAMATE_5_P_c +
0.76 PROTON_c + 0.5 aADP_c + 0.09 aL_GLUTAMATE_5_P_c

GLUTSEMIALDEHYDRROG_RXN_c 0.91 L_GLUTAMATE_5_P_c + NADPH_c + 2.21 PROTON_c +
0.09 aL_GLUTAMATE_5_P_c --> L_GLUTAMATE_GAMMA_SEMIALDEHYDE_c + NADP_c + 0.7 Pi_c +
0.3 aPi_c

SPONTPRO_RXN_c L_GLUTAMATE_GAMMA_SEMIALDEHYDE_c <=> L_DELTA1_PYRROLINE_5_CARBOXYLATE_c +
WATER_c

PYRROLINECARBREDUCT_RXN_NADP_c L_DELTA1_PYRROLINE_5_CARBOXYLATE_c + NADPH_c + PROTON_c -->
NADP_c + PRO_c

# and

PYRROLINECARBREDUCT_RXN_NAD_c L_DELTA1_PYRROLINE_5_CARBOXYLATE_c + NADH_c + PROTON_c -->
NAD_c + PRO_c

### starting from Asn
##
#

ASPARAGHYD_RXN_c ASN_c + WATER_c --> AMMONIUM_c + L_ASPARTATE_c

### from here it diverges


#####################################################
# Reactions involved in starch synthesis during the
# day (both models)
#####################################################


### PEP from C-fixation by RubisCO

RIBULOSE_BISPHOSPHATE_CARBOXYLASE_RXN_p CARBON_DIOXIDE_p + D_RIBULOSE_15_P2_p +
WATER_p --> 2.0 G3P_p + 2.0 PROTON_p

### branching of G3P fluxes towards PEP and transport to p

3PGAREARR_RXN_c G3P_p <=> 2_PG_p

2PGADEHYDRAT_RXN_p 2_PG_p <=> PHOSPHO_ENOL_PYRUVATE_p + WATER_p

PEP_Pi_pc   PHOSPHO_ENOL_PYRUVATE_p + 0.7 Pi_c + 0.3 aPi_c <=> PHOSPHO_ENOL_PYRUVATE_c
+ 0.3 PROTON_c + Pi_p

2PGADEHYDRAT_RXN_c  2_PG_c <=> PHOSPHO_ENOL_PYRUVATE_c + WATER_c


### three reactions that generate G3P in c either via transport from p of from 2_PG
```

```
# this

DHAP_3PGA_pc_01    DIHYDROXY_ACETONE_PHOSPHATE_p + G3P_c <=> DIHYDROXY_ACETONE_PHOSPHATE_c
+ G3P_p

# and

3PGA_Pi_pc_12    G3P_p + 0.7 Pi_c + 0.3 aPi_c <=> G3P_c + 0.3 PROTON_c + Pi_p

# and

3PGAREARR_RXN_c G3P_c <=> 2_PG_c


PHOSGLYPHOS_RXN_c  0.65 ATP_c + G3P_c + 0.15 PROTON_c + 0.35 aATP_c <=>
0.5 ADP_c + DPG_c + 0.5 aADP_c

GAPOXNPHOSPHN_RXN_c    GAP_c + NAD_c + 0.7 Pi_c + 0.3 aPi_c <=> DPG_c + NADH_c +
1.3 PROTON_c

TRIOSEPISOMERIZATION_RXN_c  GAP_c <=> DIHYDROXY_ACETONE_PHOSPHATE_c

F16ALDOLASE_RXN_c   FRUCTOSE_16_DIPHOSPHATE_c <=> DIHYDROXY_ACETONE_PHOSPHATE_c + GAP_c

# one part

F16BDEPHOS_RXN_c    FRUCTOSE_16_DIPHOSPHATE_c + 0.3 PROTON_c + WATER_c -->
FRUCTOSE_6P_c + 0.7 Pi_c + 0.3 aPi_c

# another part

Reaction 2_PERIOD_7_PERIOD_1_PERIOD_90_RXN_c    FRUCTOSE_6P_c + 0.65 PPI_c +
0.35 aPPI_c <=> FRUCTOSE_16_DIPHOSPHATE_c + 1.05 PROTON_c + 0.7 Pi_c + 0.3 aPi_c

PGLUCISOM_RXN_c GLC_6_P_c <=> FRUCTOSE_6P_c

PHOSPHOGLUCMUT_RXN_c    GLC_1_P_c <=> GLC_6_P_c

G6P_Pi_pc   GLC_6_P_p + 0.7 Pi_c + 0.3 aPi_c <=> GLC_6_P_c + 0.3 PROTON_c + Pi_p

PHOSPHOGLUCMUT_RXN_p    GLC_1_P_p <=> GLC_6_P_p

GLUC1PADENYLTRANS_RXN_p    0.9 ATP_p + GLC_1_P_p + 0.45 PROTON_p + 0.1 aATP_p -->
ADP_D_GLUCOSE_p + 0.55 PPI_p + 0.45 bPPI_p

GLYCOGENSYN_RXN_p   ADP_D_GLUCOSE_p --> 0.8 ADP_p + 0.8 PROTON_p + STARCH_p + 0.2 aADP_p
```

#### 3.2 ATP budgets

Plotted is the cumulative flux sum over all ATP-generating reactions.

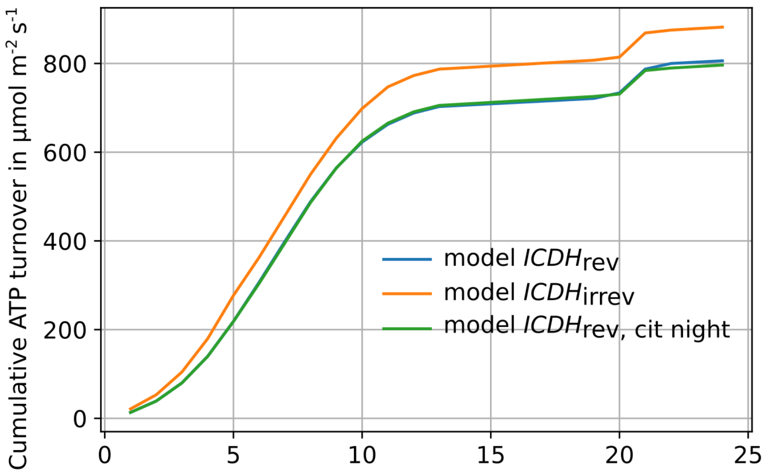


**Figure 1: Cumulative ATP turnover for the three models considered.**

#### 3.3 Environmental parameters scan

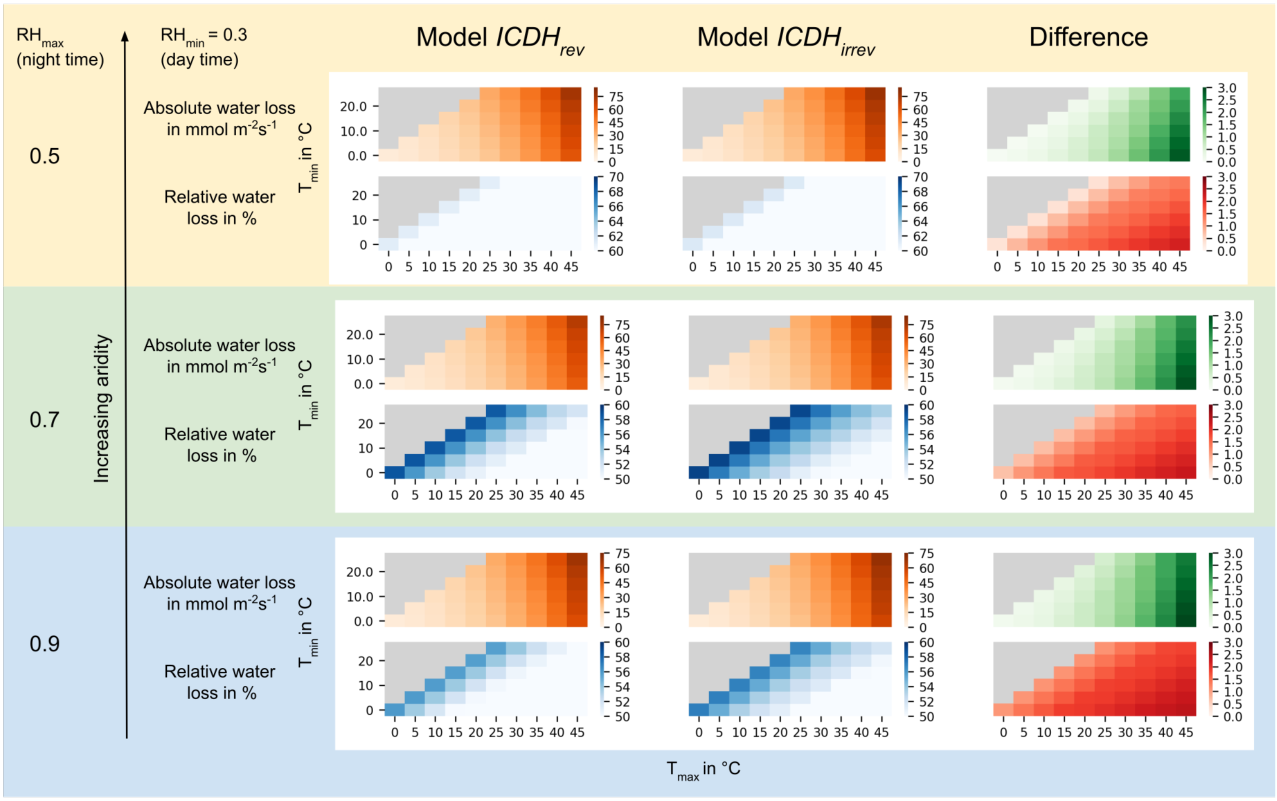


**Figure 2: $RH_{max}$-dependent water loss for model $ICDH_{rev}$ and model $ICDH_{irrev}$.** Shown is the absolute and relative water loss in dependence of $RH_{max}$ for 12 hours daylight, a light intensity of $250\mu$mol $m^{-2}s^{-1}$ and a daytime relative humidity of $RH_{min} = 0.3$.

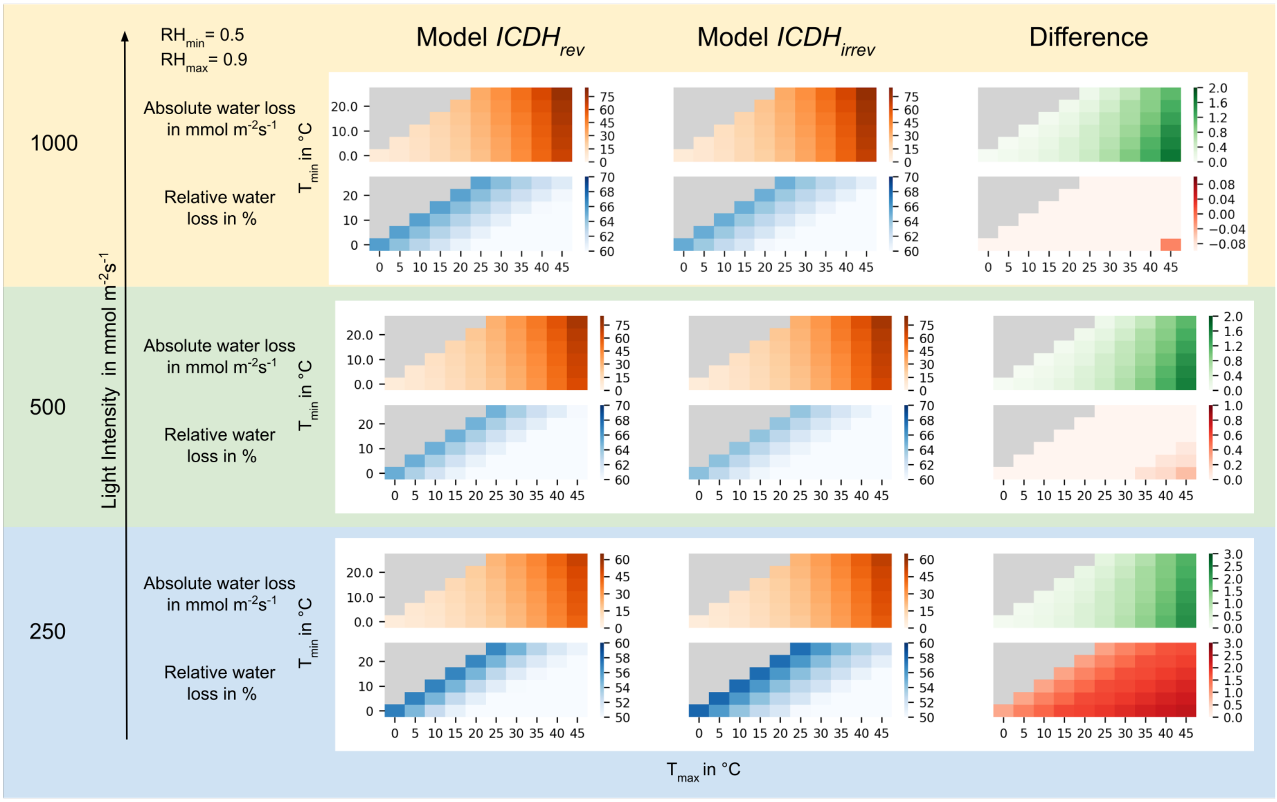


**Figure 3: Light intensity-dependent water loss for model $ICDH_{\mathrm{rev}}$ and model $ICDH_{\mathrm{irrev}}$.** Shown is the absolute and relative water loss in dependence of light intensity for 12 hours daylight, daytime relative humidity of $RH_{\min} = 0.5$ and night-time relative humidity of $RH_{\max} = 0.9$.

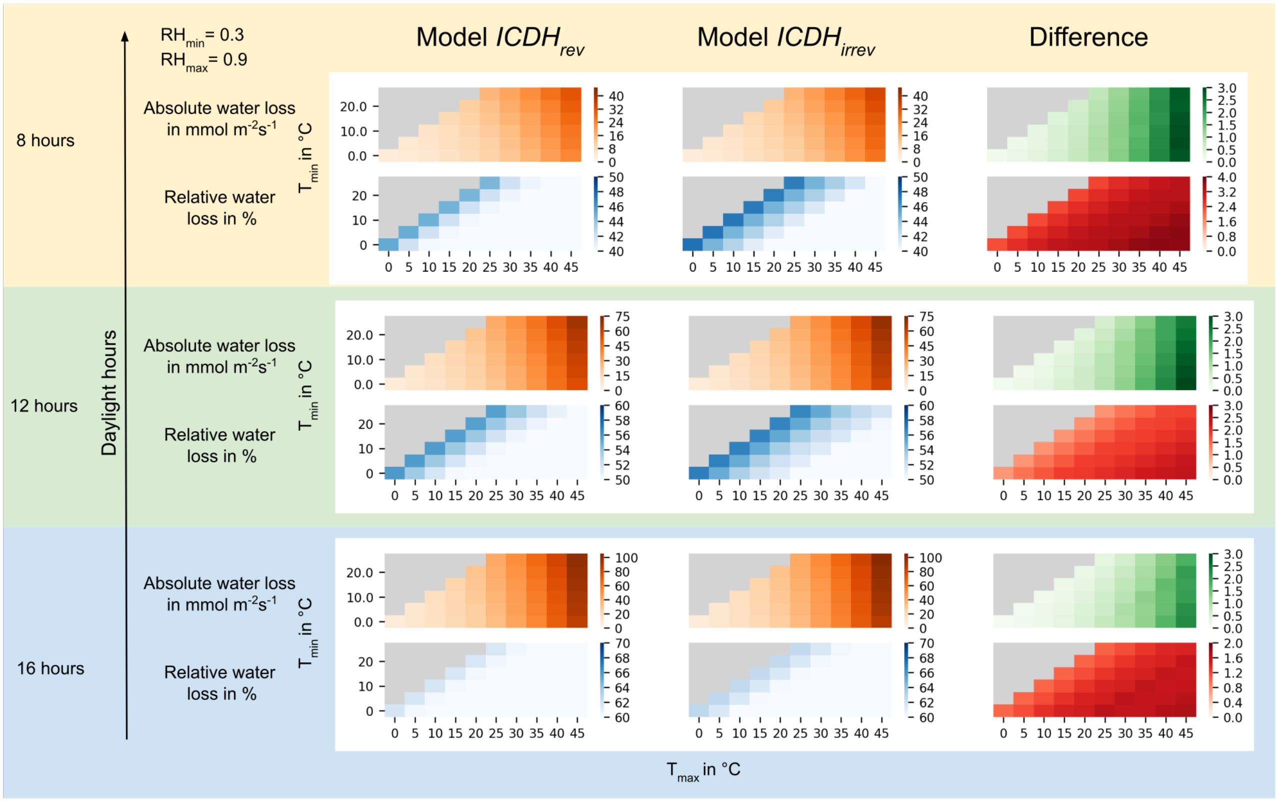


**Figure 4: Daylight hours-dependent water loss for model $ICDH_{\mathrm{rev}}$ and model $ICDH_{\mathrm{irrev}}$.** Shown is the absolute and relative water loss in dependence of daylight hours for a light intensity of 250$\mu$mol m$^{-2}$s$^{-1}$, a daytime relative humidity of $RH_{\mathrm{min}} = 0.5$ and a nighttime relative humidity of $RH_{\mathrm{max}} = 0.9$.
