## Supplementary material for "CAM emerges in a leaf metabolic model under water-saving constraints in different environments": Code: add_specific_phloem_output.py

```
#!/usr/bin/env python2
# -*- coding: utf-8 -*-

"""
    Initiated by MPM_main()
"""

import cobra

def add_phloem_composition(model,species_phloem_output):
    
    reaction = cobra.Reaction('Phloem_output_tx')
    reaction.name = 'Phloem_output_tx'
    reaction.id = 'Phloem_output_tx'
    reaction.subsystem = ''
    reaction.lower_bound = 0. 
    reaction.upper_bound = 1000. 
    model.add_reactions([reaction])
        
    if species_phloem_output == "Arabidopsis":        
        reaction.add_metabolites({
                'sSUCROSE_b':- 0.6893085734,
                'GLC_c': - 0.0,
                'FRU_c': - 0.0,
                'sASP_b': - 0.02976559749,
                'sGLU_b': - 0.03571871698,
                'ASN_c': - 0.03164552996,
                'SER_c': - 0.02412580007,
                'sGLN_b': - 0.1040229302,
                'GLY_c': - 0.002193254552,
                'THR_c': - 0.01754603641,
                'sALA_b': - 0.03039224164,
                'sGABA_b': - 0.0,
                'VAL_c': - 0.007519729891,
                'ILE_c': - 0.003759864946,
                'PHE_c': - 0.003446542867,
                'LEU_c': - 0.003759864946,
                'LYS_c': - 0.005639797418,
                'ARG_c': - 0.005013153261,
                'HIS_c': - 0.001253288315,
                'CYS_c': -0.0,
                'PRO_c': -0.0,
                'MET_c': - 0.002193254552,
                'TYR_c': - 0.001879932473,
                'TRP_c': - 0.002819898709,
                'MAL_c': - 0.0,
                'PROTON_e':-1.0 ,
                'PROTON_c': 1.0})
    
    elif  species_phloem_output == "Opuntia":    
        reaction.add_metabolites({
            'sSUCROSE_b':  - 0.703517588,
            'GLC_c': - 0.001546192518,
            'FRU_c': - 0.001546192518,
            'sASP_b': - 0.003146501699,
            'sGLU_b': - 0.009153459563,
            'ASN_c': - 0.01129880174,
            'SER_c': - 0.008009277117,
            'sGLN_b': - 0.03604174718,
            'GLY_c': - 0.01129880174,
            'THR_c': - 0.0,
            'sALA_b': - 0.004862775418,
            'sGABA_b': - 0.0,
            'VAL_c': - 0.03890220335,
            'ILE_c': - 0.03947429452,
            'PHE_c': - 0.02002319284,
            'LEU_c': - 0.03604174718,
            'LYS_c': - 0.02431387709,
            'ARG_c': - 0.002002319254,
            'HIS_c': - 0.0,
            'CYS_c': - 0.0,
            'PRO_c': - 0.001144182445,
            'MET_c': - 0.008009277117,
            'TYR_c': - 0.02946269815,
            'TRP_c': - 0.002860456164,
            'MAL_c': - 0.007344414335,
            'PROTON_e': - 1.0,
            'PROTON_c': 1.0})

    elif  species_phloem_output == "Tomato":
        reaction.add_metabolites({
            'sSUCROSE_b':  - 0.7628865979,
            'GLC_c': - 0.07216494845,
            'FRU_c': - 0.0824742268,
            'sASP_b': - 0.006268041237,
            'sGLU_b':- 0.01294845361 ,
            'ASN_c': - 0.001567010309,
            'SER_c': - 0.00412371134,
            'sGLN_b': - 0.02507216495,
            'GLY_c': - 0.0007422680412,
            'THR_c': - 0.007175257732,
            'sALA_b': - 0.004041237113,
            'sGABA_b': - 0.002391752577,
            'VAL_c': - 0.002886597938,
            'ILE_c': - 0.00181443299,
            'PHE_c': - 0.00593814433,
            'LEU_c': - 0.002144329897,
            'LYS_c': - 0.002309278351,
            'ARG_c': - 0.0004359351988,
            'HIS_c': - 0.0004359351988,
            'CYS_c': - 0.0004359351988,
            'PRO_c': - 0.0004359351988,
            'MET_c': - 0.0004359351988,
            'TYR_c': - 0.0004359351988,
            'TRP_c': - 0.0004359351988,
            'MAL_c': - 0.0,
            'PROTON_e':-1.0 ,
            'PROTON_c': 1.0})

        
    return model
```
