## Supplementary material for "CAM emerges in a leaf metabolic model under water-saving constraints in different environments": Code: helping_functions.py

```
#!/usr/bin/env python2
# -*- coding: utf-8 -*-

import numpy as np

def gaussian(x, level, delta, mean, sigma, shift):
    
    """

    Initiated by MPM_main()    

    - generates normal distributed curves to simulate light, RH, and temperature
    - curves are shifted so they form a continuous curve over the 24h simulation period
    
    parameters:
    ---------
    - level: baseline value for the curve
    - delta: heights of the curve
    - mean: mid-day
    - shit of curve from mid day towards afternoon
    - sigma: widths of the curve - calculated to match sunrise 
 
    """

    if x > (24 - (mean - shift)):
        value = level + delta * np.exp(-np.power(x - 24 - (mean + shift), 2.) / (2 * np.power(sigma, 2.)))
    else:
        value = level + delta * np.exp(-np.power(x - (mean + shift), 2.) / (2 * np.power(sigma, 2.)))
    return value
```
