## Supplementary material for "CAM emerges in a leaf metabolic model under water-saving constraints in different environments": Code: plotting_functions.py

```
#!/usr/bin/env python2
# -*- coding: utf-8 -*-

import numpy as np
import os
import matplotlib.pyplot as plt

import helping_functions as hf


def plot_generated_data(parameter_file, th_dict, sigma,constants):
    
    """
    Initiated by MPM_main()
    
    parameters:
    ------
     - parameter file
     - th_dict
     - sigma (parameter for Gauss curve that makes the curve match a certain day length)
   
    """

    # light curve
    min_light = 0
    mid_day = float(constants['mid_day'])
    Delta_light = float(th_dict['MaxLight'])
    shift_light = 0
    
    # temperature curve
    minT_celsius_theoretical = float(th_dict['MinT'])
    maxT_celsius = float(th_dict['MaxT'])
    day_length = float(th_dict['DayLength'])
    shift_temperature = float(th_dict['Shift_temperature'])
    temp_adjust = float(th_dict['TCurveAdjustment'])  # if below zero cuve gets narrower
   
    # humidity curve
    max_humidity = float(th_dict['MaxHumidity'])
    min_humidity = float(th_dict['MinHumidity'])
    shift_humidity = float(th_dict['Shift_humidity'])
    humidity_adjust = float(th_dict['RHCurveAdjustment']) # if below zero cuve gets narrower
 
    # calculations
    minT_theoretical = minT_celsius_theoretical #+ 273.15
    maxT = maxT_celsius# + 273.15
    deltaT  = maxT-minT_theoretical
    deltaH = min_humidity-max_humidity
    
    xx=np.linspace(0, 23, 100)
    
    g_humidity=list()
    for i in xx:
        g_humidity.append(hf.gaussian(i,max_humidity, deltaH, mid_day, humidity_adjust * day_length, shift_humidity))
    
    g_temperature=list()
    for i in xx:
        g_temperature.append(hf.gaussian(i, minT_theoretical, deltaT, mid_day, temp_adjust * day_length, shift_temperature)) 
    
    g_light=list()        
    for i in xx:
        g_light.append(hf.gaussian(i,min_light, Delta_light, mid_day, sigma, shift_light))
      
    
    fig, ax1 = plt.subplots()
    
    ln1=ax1.plot(xx, g_light,'y',label='Light',linewidth=2.8)
    ax1.set_ylim([0,1.1* Delta_light])
    ax1.set_ylabel('Light intensiy in $\mu mol/m^2/s$', fontsize=14)
    lns = ln1
    labs = [l.get_label() for l in lns]
    ax1.legend(lns, labs, loc=0,frameon=False, fontsize=13)
    fig.tight_layout()
    plt.show()
    fig.savefig(os.path.join("Results/Light.png"),dpi=600)

    
    fig, ax1 = plt.subplots()
    
    ax2 = ax1.twinx()
    ax1.set_ylim([minT_theoretical-2,minT_theoretical+deltaT+2])
    ax1.set_ylabel('Temperature in $\circ$C',fontsize=14)
    ax2.set_ylim([min_humidity-0.1,1])
    ax2.set_ylabel('Relative Humidity', fontsize=14)
    
    ln1=ax1.plot(xx, g_temperature,'orangered',label='Temperature',linewidth=2.8)
    ln2=ax2.plot(xx,g_humidity,'steelblue',label='Relative Humidity',linewidth=2.8)
    
    lns = ln1+ln2
    labs = [l.get_label() for l in lns]
    ax2.legend(lns, labs, loc=0,frameon=False, fontsize=13)
    fig.tight_layout()
    plt.show()
    fig.savefig(os.path.join("Results/Temp_Hum.png"),dpi=600)

  
```
