## Supplementary material for "CAM emerges in a leaf metabolic model under water-saving constraints in different environments": Code: writing_functions.py

```
#!/usr/bin/env python2
# -*- coding: utf-8 -*-

import csv

def write_results(csv_file, new_model, solution, pareto=1, write_method='wb'):
    
    """
    Initiated by MPM_main()
        
    Writes mpm_solve results to .csv file

    parameters:
    ------
    - csv_file: file which is opened
    - new_model: new model
    - solution: FBA/FVA solution in dictionary
    - pareto: Current Pareto number (1 for FBA or FVA analysis)
    - write_method:
        
    """

    reaction_pathway, pathway_priority = read_pathways()

    writer = csv.writer(csv_file, delimiter='\t')
    
    if write_method == 'wb':
        writer.writerow(['ID', 'Name', 'Flux', 'Pareto', 'Equation', 'Compartment', 'Pathway', 'Priority', 'EC', 'Original_ID', 'Phase', 'Reverse', 'lb', 'ub'])

    for key, value in solution.fluxes.items():
        key_orig, reverse, pathway, protein_class, compartments, original_id, phase, lb, ub = parse_results_for_csv(new_model, key, reaction_pathway)

        writer.writerow([key,new_model.reactions.get_by_id(key_orig).name, value,pareto, new_model.reactions.get_by_id(key).build_reaction_string(),
                         compartments,pathway,pathway_priority[pathway.lower()],protein_class,original_id,phase,reverse,lb,ub])
    
    
def read_pathways():
    
    """
    Instantiated by write_results, finds a pathway for every reaction in the model to add to the exported results

    returns:
    ------
    reaction_pathway: dictionary of reaction:pathway
    pathway_priority: dictionary of pathway:number assignment (just for ordering them in an excel sheet).
    
    """

    reaction_pathway = {}
    with open('Parameters/rxnpathwaydict.csv', 'r') as csv_file:
        reader = csv.reader(csv_file, delimiter=',')
        for k, v in reader:
            reaction_pathway[k] = v
    pathway_priority = {}
    with open('Parameters/pathwayprioritizer.csv', 'r') as csv_file:
        reader = csv.reader(csv_file, delimiter='\t')
        for k, v in reader:
            pathway_priority[k.lower()] = v

    return reaction_pathway, pathway_priority


def parse_results_for_csv(new_model, key, reaction_pathway):
    
    """
    Catch-all parsing for writing solver results to .csv. Just a few hacks/query's.

    returns:
    ------
    a bunch
    
    """
    if 'reverse' in key:
        reverse = True
        key_orig = key[:-8]
    else:
        reverse = False
        key_orig = key

    try:
        pathway = reaction_pathway[key_orig[:-3]]
    except KeyError:
        if "_L_" in key:
            pathway = 'Linker'
        else:
            pathway = ''

    try:
        protein_class = new_model.reactions.get_by_id(key_orig).notes['PROTEIN CLASS'][0]
    except KeyError:
        protein_class = 'NA'
    try:
        compartments = " + ".join(new_model.reactions.get_by_id(key_orig).get_compartments())
    except TypeError:
        compartments = "None"
    original_id = 'NA'
    try:
        original_id = new_model.reactions.get_by_id(key_orig).notes['Original_ID']
        if reverse:
            original_id += '_reverse'
    except KeyError:
        if "_L_" in key:
            original_id = key[:-6]
    try:
        phase = new_model.reactions.get_by_id(key_orig).notes['phase']
    except KeyError:
        if "_L_" in key_orig:
            phase = key_orig[-5:-3]
        else:
            phase = new_model.reactions.get_by_id(key_orig).id[-13:-11]

    lb = new_model.reactions.get_by_id(key_orig).lower_bound
    ub = new_model.reactions.get_by_id(key_orig).upper_bound

    return key_orig, reverse, pathway, protein_class, compartments, original_id, phase, lb, ub
```
