## Supplementary material for "CAM emerges in a leaf metabolic model under water-saving constraints in different environments": Code: MPM_main.py

```
#!/usr/bin/env python
"""
MPM_main for CobraPy:

"""

from __future__ import division, print_function, absolute_import
import csv
import os
import xml.etree.ElementTree as etree
import cobra
import numpy as np
from itertools import chain
from cobra.util import solver as sutil
from cobra.core.solution import get_solution
from optlang.symbolics import add, Zero
import pandas as pd

# read own files
from functions import plotting_functions as pf
from functions import helping_functions as hf
from functions import writing_functions as wf
from functions import add_specific_phloem_output as phloem

def get_generated_data(climate_file, Light_Comp):
 
            wv_file_name = 'Temperature_dependent_WaterVapour.csv'
            wv_file='Parameters/' + wv_file_name 
        
            reader = csv.reader(filter(lambda row: row[0]!='#', open(wv_file,'r')))
            wv_dict = {}
            for row in reader:
                k, v = row
                wv_dict[k] = v
                
                
            th_file_name = climate_file # 'Temperature_and_Humidity_Curves.csv'
            th_file='Parameters/' + th_file_name 
        
            reader = csv.reader(filter(lambda row: row[0]!='#', open(th_file,'r')))
            th_dict = {}
            for row in reader:
                k, v = row
                th_dict[k] = v  
            
            # generate light_dark list based on given day length             
            light_dark = list(['dark']) * 24
            day_length=float(th_dict['DayLength'])
            
            if day_length > 24:
                raise ValueError('The day length should be between 0 and 24 hours.')

            mid_day = float(constants['mid_day'])           
            sunrise = int(round(mid_day - day_length/2))
            sunset = int(round(mid_day + day_length/2)) # this makes sure that the last hour of sunlight is counted at light
            
            if sunrise < 0:    
                for i in xrange(0, sunset + 1):
                    light_dark[i]='light'
                for i in xrange(24 + sunrise, 24):
                    light_dark[i]='light'
            else:
                for i in xrange(sunrise, sunset + 1):
                    light_dark[i]='light'
            
            print('Day lenght was rounded to ' + str(light_dark.count('light')) + ' hours.')

            # calculate width (sigma) of the Gauss curve from maxLight and sunset             
            max_light = float(th_dict['MaxLight'])
            
            sigma = 0.1
            while hf.gaussian(sunset, 0, max_light, mid_day, sigma, 0) < Light_Comp: # below the light compensation point we consider it to be dark
                sigma = sigma + 0.1
                             
            # calculate mean light intensity    
            total_light=0
            for x in range(24):
                light=hf.gaussian(x, 0, max_light, mid_day, sigma, 0)
                total_light+=light
            
            mean_light=total_light/float(light_dark.count("light"))
            
            return wv_dict, th_dict, light_dark, sigma, mean_light   

           
def get_generated_data_parameter_scan( th_dict, Light_Comp):            
          
            # generate light_dark list based on given day length             
            light_dark = list(['dark']) * 24
            day_length=float(th_dict['DayLength'])
            
            if day_length > 24:
                raise ValueError('The day length should be between 0 and 24 hours.')

            mid_day = float(constants['mid_day'])           
            sunrise = int(round(mid_day - day_length/2))
            sunset = int(round(mid_day + day_length/2)) # this makes sure that the last hour of sunlight is counted at light
            
            if sunrise < 0:    
                for i in xrange(0, sunset + 1):
                    light_dark[i]='light'
                for i in xrange(24 + sunrise, 24):
                    light_dark[i]='light'
            else:
                for i in xrange(sunrise, sunset + 1):
                    light_dark[i]='light'
            
            print('Day lenght was rounded to ' + str(light_dark.count('light')) + ' hours.')

            # calculate width (sigma) of the Gauss curve from maxLight and sunset             
            max_light = float(th_dict['MaxLight'])
            
            sigma = 0.1
            while hf.gaussian(sunset, 0, max_light, mid_day, sigma, 0) < Light_Comp: # below the light compensation point we consider it to be dark
                sigma = sigma + 0.1
                             
            # calculate mean light intensity    
            total_light=0
            for x in range(24):
                light=hf.gaussian(x, 0, max_light, mid_day, sigma, 0)
                total_light+=light
            
            mean_light=total_light/float(light_dark.count("light"))
            
            return  light_dark, sigma, mean_light 

        

def mpm_create(c_model, name, species_phloem_output, xml , fixed_light, Light_Comp, wv_dict, th_dict, light_dark, mean_light, **kwargs):
    
    """
    Create a Multi-Phase-Model fom a COBRA model

    parameters:
    ------
    - c_model: input model filename
    - name: output model name (and filename)
    - xml: parameter xml file
    - wv_dict: temperture dependent water vapour pressure dictionary 
    - th_dict: temperature and humidity curve generating parameters file

    return:
    --------
    - model: MPM model
    - nr_phases: number of phases in model (inferred from parameter file)
    
    """

    model = cobra.io.read_sbml_model('Models/' + c_model) 
    
    # remove old biomass reaction and add new one
    model.reactions.Phloem_output_tx.remove_from_model()
    model = phloem.add_phloem_composition(model,species_phloem_output)
    
#    # add isoctrate storage in vacuole
#    aISO_CITv = cobra.Metabolite('aTHREO_DS_ISO_CITRATE_v', formula='C6H5O7', name='aTHREO_DS_ISO_CITRATE', compartment='v')
#    ISO_CITv = cobra.Metabolite('THREO_DS_ISO_CITRATE_v', formula='C6H5O7' , name='THREO_DS_ISO_CITRATE', compartment='v')
#    
#    model.add_metabolites([aISO_CITv])
#    model.add_metabolites([ISO_CITv])
#
#    reaction = cobra.Reaction('THREO_DS_ISO_CITRATE_PROTON_rev_vc')
#    reaction.name = 'THREO_DS_ISO_CITRATE_PROTON_rev_vc'
#    reaction.id = 'THREO_DS_ISO_CITRATE_PROTON_rev_vc'
#    reaction.subsystem = ''
#    reaction.lower_bound = 0. 
#    reaction.upper_bound = 1000. 
#    model.add_reactions([reaction])
#    reaction.add_metabolites({'THREO_DS_ISO_CITRATE_v': -0.5, 'PROTON_v': -2.5,'aTHREO_DS_ISO_CITRATE_v': -0.5, 'THREO_DS_ISO_CITRATE_c': 1.0, 'PROTON_c': 3.0})
#
#    reaction = cobra.Reaction('THREO_DS_ISO_CITRATE_PROTON_vc')
#    reaction.name = 'THREO_DS_ISO_CITRATE_PROTON_vc'
#    reaction.id = 'THREO_DS_ISO_CITRATE_PROTON_vc'
#    reaction.subsystem = ''
#    reaction.lower_bound = 0. 
#    reaction.upper_bound = 1000. 
#    model.add_reactions([reaction])
#    reaction.add_metabolites({'THREO_DS_ISO_CITRATE_c': -1.0, 'PROTON_v': -0.5,'aTHREO_DS_ISO_CITRATE_v': 0.5, 'THREO_DS_ISO_CITRATE_v': 0.5})
#  
    ### adjust model     
    #block backwards reaction of reactions.ISOCITDEH_RXN_m - change to simlate model ICDHrev and model ICDHirrev    
    model.reactions.ISOCITDEH_RXN_m.lower_bound = 0
    model.reactions.ISOCITDEH_RXN_c.lower_bound = 0
    model.reactions.ISOCITDEH_RXN_x.lower_bound = 0
    
    # block putative mal/cit antiporter
    model.reactions.MAL_CIT_rev_vc.lower_bound = 0
    model.reactions.MAL_CIT_rev_vc.upper_bound = 0
    model.reactions.MAL_CIT_vc.lower_bound = 0
    model.reactions.MAL_CIT_vc.upper_bound = 0
        
    # generate 'water loss' metabolite
    WATER_trans_e = cobra.Metabolite('WATER_trans_e',formula='H2O1', name='WATER_trans', compartment='e')
    model.add_metabolites(WATER_trans_e)
    
    # add Co2 backwards reaction to model water loss whenever stomata are open
    reaction = cobra.Reaction('CO2_tx_backwards')
    reaction.name = 'CO2_tx_backwards'
    reaction.id = 'CO2_tx_backwards'
    reaction.subsystem = ''
    reaction.lower_bound = 0. 
    reaction.upper_bound = 1000. 
    model.add_reactions([reaction])
   
    reaction.add_metabolites({'CARBON_DIOXIDE_e': -1.0})    
    
    model.reactions.Plastoquinol_Oxidase_p.upper_bound = 0.0
    
    model.id = name
    xml_tree = etree.parse('Parameters/' + xml)
    xml_root = xml_tree.getroot() 
    nr_phases = len(xml_root.find('phases'))
    if light_dark == None:
        light_dark = [x.attrib['light_dark'] for x in xml_root.find('phases')]  
        print('light_dark calculated')

    # multipy model and add respective compartment names for each phase of the day
    # this generates a model with 24 x nr comps compartments    
    original_compartments = model.compartments.copy()
    
    print("\nCreating compartments...")

    for key, value in model.compartments.items():    
        model.compartments["{k}_01".format(k=key)] = ("{v} [01]".format(v=model.compartments.pop(key)))

    # Compartments: create for phases 2+ 
    for phase in range(2, nr_phases + 1):
        for key, value in original_compartments.items():
            model.compartments["{k}_{p}".format(k=key, p=str(phase).zfill(2))] = ("{v} [{p}]".format(v=value, p=str(phase).zfill(2)))
    
    # Metabolites: create for phases 2+ (before phase 1, because they will be renamed)
    print("\nCreating metabolites...")
    original_metabolites = [i for i in model.metabolites]
    for phase in range(2, nr_phases + 1):
        for metabolite in original_metabolites: 
            new_metabolite = metabolite.copy()
            new_metabolite.notes['Original_ID'] = metabolite.id
            new_metabolite.notes['phase'] = str(phase).zfill(2)
            new_metabolite.compartment = "{m}_{p}".format(m=metabolite.compartment,p=str(phase).zfill(2))
            new_metabolite.id = "{m}_{p}".format(m=metabolite.id,p=str(phase).zfill(2))
            new_metabolite.name = "{m} [{p}]".format(m=metabolite.name,p=str(phase).zfill(2))
            new_metabolite.charge = metabolite.charge
            model.add_metabolites(new_metabolite)

    # Metabolites: rename originals for phase 1
    for metabolite in original_metabolites:    
        metabolite.notes['Original_ID'] = metabolite.id
        metabolite.notes['phase'] = '01'
        metabolite.id = "{m}_01".format(m=metabolite.id)
        metabolite.compartment = "{m}_01".format(m=metabolite.compartment)
        metabolite.name = "{m} [01]".format(m=metabolite.name)
    
    # make sure all new IDs are sbml compatible        
    cobra.manipulation.modify.escape_ID(model)  

    # Reactions: create for all phases
    print("\nCreating reactions...")
    original_reactions = [i for i in model.reactions]  
    for phase in range(1, nr_phases + 1):
        for reaction in original_reactions: 
            new_reaction = reaction.copy()
            new_reaction.notes['Original_ID'] = reaction.id
            new_reaction.notes['phase'] = str(phase).zfill(2)
            new_reaction.id = "{r}_{p}".format(r=reaction.id,p=str(phase).zfill(2))
            new_reaction.name = "{r} [{p}]".format(r=reaction.name,p=str(phase).zfill(2))

    # add new metabolite IDs to reactions
            new_reactants = {}
            for key, value in new_reaction.metabolites.iteritems():
                new_reactants["{k}_{p}".format(k=key.id[:-3],p=str(phase).zfill(2))] = value               
                
            remove_reactants = {}
            for key, value in new_reaction.metabolites.iteritems():
                remove_reactants[key] = -value
            model.add_reaction(new_reaction) 
            new_reaction.add_metabolites(new_reactants)             
            new_reaction.add_metabolites(remove_reactants)    

    for reaction in original_reactions:
        reaction.remove_from_model()
        
    # Correct Metabolites' ID if necessary    
    cobra.manipulation.modify.escape_ID(model)  
    
    print("\nAdding Linker reactions...")
    for phase_num in range(1, nr_phases + 1):
        for linker in xml_root.find('linkage'):
            mpm_set_linkers(model, phase_num, linker, nr_phases) 

    print("Adding Water Vapour Stoichiometry...")
    for p, phase in enumerate(xml_root.find('phases')):  
        mpm_set_watervapour(model, p, phase,  wv_dict,th_dict) 
            
    if mean_light == None:
        mean_light=calculate_mean_light(xml_root,nr_phases, light_dark)
        
    if xml_root.find('phases')[0].find('reaction') is not None: 
        print("Setting Phase-specific constraints...")
        mpm_set_phase_constraints(model, xml_root, fixed_light, th_dict, light_dark, mean_light, Light_Comp)
    else:
        print("\nNo Phase-specific constraints found.")

    if len(list(xml_root.find('balances'))) > 0:       
        print('Adding Balances...')
        for balance in xml_root.find('balances'):
            mpm_set_balances(model, balance, nr_phases, fixed_light, th_dict, light_dark) 
    else:
        print('No Balance constraints found')


    mpm_create_common_water_vapour(model, nr_phases)
    
    print("\nSaving Model...")
    cobra.io.write_sbml_model(model,"Models/" + "{}_{}.xml".format(name, metric), use_fbc_package=False)
    
    print("Done building the multi-phase model!")
    
    print("Setting model solver to "+str(solver))
    model.solver = 'gurobi'
  
    return model, nr_phases


def mpm_set_linkers(new_model, phase, linker, nr_phases):
    
    """
    Initiated by mpm_create()
    Create MPM Linker reactions in model

    parameters:
    ------
    - new_model: MPM model
    - phase: current phase number
    - linker: current linker metabolite
    - nr_phases: total number of phases in model
    
    """

    # Retrieve bounds
    if 'lower' in linker.keys():
        lower_bound = float(linker.attrib['lower'])
    else:
        lower_bound = 0

    if 'upper' in linker.keys():
        upper_bound = float(linker.attrib['upper'])
    else:
        upper_bound = 1000

    if phase == nr_phases:  # Link last phase back into the first phase
        if int(linker.attrib['uni']) == 1: 
            lower_bound = 0
            upper_bound = 0
        phase1 = phase # if last phase set string for transitions back to 1
        phase2 = 1
    else:
        phase1 = phase # else simpy add 1
        phase2 = phase + 1

    # Create reaction
    new_reaction = cobra.Reaction("{l}_L_{p1}_{p2}".format(l=linker.attrib['id'],p1=str(phase1).zfill(2),p2=str(phase2).zfill(2)),name="Linker {l} [{p1}] to {l} [{p2}]".format(l=linker.attrib['id'],p1=str(phase1).zfill(2),p2=str(phase2).zfill(2)),subsystem="Linker")
    new_reaction.lower_bound = lower_bound
    new_reaction.upper_bound = upper_bound

    # Set stoichiometry
    metabolite_query1 = "{l}_{p}".format(l=linker.attrib['id'],p=str(phase1).zfill(2))
    metabolite_query2 = "{l}_{p}".format(l=linker.attrib['id'],p=str(phase2).zfill(2))
    
    metabolite_linker1 = new_model.metabolites.get_by_id(metabolite_query1)
    metabolite_linker2 = new_model.metabolites.get_by_id(metabolite_query2)
    
    reactants = {metabolite_linker1: -1, metabolite_linker2: 1}
    new_reaction.add_metabolites(reactants)
    new_model.add_reaction(new_reaction)


def mpm_set_watervapour(new_model, p, phase, wv_dict, th_dict):
    
    """
    Initiated by mpm_create()
    Create Gas Diffusion constraint from data in parameter file or from normal distribution
    
    parameters:
    ------
    - new_model: MPM model
    - phase: phase parameter data (holds gas diffusion parameters)
    - wv_dict: temperture dependent water vapour pressure dictionary 
    - th_dict: temperature and humidity curve generating parameters file

    """

    if th_dict==None:
      
        temp = phase.attrib['temp']
        humidity = phase.attrib['RH']
        conc_H2O_in = phase.attrib['C_H2O_in']
        conc_H2O_out = phase.attrib['C_H2O_out']
   
        delta_C_H2O = eval("{C_H2O_in}-({C_H2O_out}*{RH})".format(C_H2O_in=conc_H2O_in,C_H2O_out = conc_H2O_out, RH=humidity)) 
                
    else:
        
        # temperature curve
        min_T_celsius_theoretical = float(th_dict['MinT'])
        max_T_celsius = float(th_dict['MaxT'])
        shift_temperature = float(th_dict['Shift_temperature'])
        temp_adjust = float(th_dict['TCurveAdjustment'])  # if below zero cuve gets narrower
   
        # humidity curve
        max_humidity = float(th_dict['MaxHumidity'])
        min_humidity = float(th_dict['MinHumidity'])
        shift_humidity = float(th_dict['Shift_humidity'])
        humidity_adjust = float(th_dict['RHCurveAdjustment']) # if below zero cuve gets narrower

        mid_day = float(constants['mid_day'])              
        day_length = float(th_dict['DayLength'])

        # calculations        
        min_T_theoretical = min_T_celsius_theoretical + 273.15
        max_T = max_T_celsius + 273.15
        delta_T = max_T - min_T_theoretical
        delta_H = min_humidity - max_humidity
        
        temp = hf.gaussian(p, min_T_theoretical, delta_T, mid_day, temp_adjust * day_length, shift_temperature)
        humidity = hf.gaussian(p, max_humidity, delta_H, mid_day, humidity_adjust * day_length, shift_humidity)

        delta_C_H2O = calculate_delta_C_H2O(float(temp), float(humidity), wv_dict)  

    delta_C_CO2 = float(constants['deltaC_CO2']) # CO2 difference between inside and outside of the leaf - fixed value
    D_H2O_0 = float(constants['D_H2O_0']) # diffusion coefficient H2O_0 
    D_CO2_0 = float(constants['D_CO2_0']) # diffusion coefficient CO2_0 
        
    # equations 
    D_H2O = eval("{dH2O_0}*({T}/273) * 10**1.8".format(dH2O_0=D_H2O_0, T=temp))
    D_CO2 = eval("{dCO2_0}*({T}/273) * 10**1.8".format(dCO2_0=D_CO2_0, T=temp))

    Stoichiometry=eval("({D_H2O}*{deltaC_H2O})/({D_CO2}*{deltaC_CO2})".format(D_H2O=D_H2O, deltaC_H2O=delta_C_H2O, D_CO2=D_CO2, deltaC_CO2=delta_C_CO2)) 
  
    # get co2 transport from model and add water loss with pre-calculated stoichiometry forward
    co2_reaction = new_model.reactions.get_by_id("CO2_tx_{}".format(phase.attrib['num'].zfill(2))) 
    co2_reaction.add_metabolites({new_model.metabolites.get_by_id("WATER_trans_e_{}".format(phase.attrib['num'].zfill(2))): Stoichiometry})
    co2_reaction.lower_bound=0

    co2_reaction = new_model.reactions.get_by_id("CO2_tx_backwards_{}".format(phase.attrib['num'].zfill(2)))
    co2_reaction.add_metabolites({new_model.metabolites.get_by_id("WATER_trans_e_{}".format(phase.attrib['num'].zfill(2))): Stoichiometry})
    co2_reaction.lower_bound=0
 

def calculate_delta_C_H2O(T, RH, wv_dict):
    
    """
    Initiated by mpm_set_watervapour()
    
    this function looks up the temperature-dependent sturation water concentration
        in mol/m^3 and calculates delta H20 (outside and inside the leaf) by assuming 
        that the leaf temperature is 2 degrees more than outside and RH inside  = 1
    
    table extracted from "Physicochemical and Environmental Plant Physiology
        FOURTH EDITION"
        
    parameters:
    ------
    - T: temperature in K
    - RH: relative humidity of the air
    - wv_dict: temperture dependent water vapour pressure dictionary 
   
    """
    deltaT = float(constants['deltaT'])
    
    c_H2O_sat_out=float(wv_dict[str(int(round(T-273.15)))]) # convert from Kelvin to Celsius for lookup table
    c_H2O_sat_in=float(wv_dict[str(int(round(T-273.15+ deltaT)))])
    
    deltaC_H2O = c_H2O_sat_in - (c_H2O_sat_out * RH) 
    
    return deltaC_H2O


def calculate_mean_light(xml_root,nr_phases, light_dark): 
    
    """
    Initiated by mpm_create()
    
    this functions is called if use_real_data = 1
    - it sums up the light fractions over the course of the day, normalizes it by the max light intensity and defides by day length
    
    """ 
    total_light=0 
    
    for p, phases in enumerate(xml_root.find('phases')): 
        if phases.attrib['num']=='7': # midday
            for reaction in phases.findall('reaction'):
                if reaction.attrib['id']=='Photon_tx':
                    max_light = float(reaction.attrib['upper'])

    if len(list(xml_root.find('balances'))) > 0:         
        for balance in xml_root.find('balances'):
            if balance.attrib['type']  == 'light':
                reaction = balance[0]
                for phase in range(1, int(nr_phases + 1)):
                    total_light += float(reaction[phase - 1].attrib['b'])
    
        mean_light=total_light * max_light /float(light_dark.count("light"))
        
    else:
        raise AttributeError('No Light Balance recognized.') 
    
    return mean_light  
  
    
def mpm_set_phase_constraints(new_model, xml_root, fixed_light, th_dict, light_dark, mean_light, Light_Comp):
    
    """
    Initiated by mpm_create()
    Create phase constraints according to parameter file
    - for ATPase (maintenance) calculate lower and upper bounds based on max/average light intensity and day - night pattern 
    
    parameters:
    ------
    - new_model: MPM model
    - xml_root: parameter xml data
    - light_dark: information about light or dark periode to calculate maintenance cost
    - th_dict: temperature and humidity curve generating parameters file
        
    """

    if th_dict==None:  
        
        for p, phases in enumerate(xml_root.find('phases')):  
            for reaction in phases.findall('reaction'):
                reaction_query = "{r}_{p}".format(r=reaction.attrib['id'],p=str(p + 1).zfill(2))
                new_reaction = new_model.reactions.get_by_id(reaction_query)                  
                if reaction.attrib['id']=='ATPase_tx':     
                    
                            if light_dark[p]=='light':        
                               new_reaction.lower_bound = estimate_maintenance(mean_light)  
                               new_reaction.upper_bound = estimate_maintenance(mean_light) 
                            else:
                               new_reaction.lower_bound = 1/3 * estimate_maintenance(mean_light)
                               new_reaction.upper_bound = 1/3 * estimate_maintenance(mean_light)                       
                else:            
                    if 'lower' in reaction.keys():
                        new_reaction.lower_bound = float(reaction.attrib['lower'])
                    if 'upper' in reaction.keys():
                        new_reaction.upper_bound = float(reaction.attrib['upper'])      
    else:
  
        max_light = float(th_dict['MaxLight'])
        min_light = float(0)
        shift_light = float(0) 

        mid_day = float(constants['mid_day'])
        
        for p, phases in enumerate(xml_root.find('phases')): 
            for reaction in phases.findall('reaction'):
                reaction_query = "{r}_{p}".format(r=reaction.attrib['id'],p=str(p + 1).zfill(2))
                new_reaction = new_model.reactions.get_by_id(reaction_query)  
                if reaction.attrib['id']=='ATPase_tx':     

                            if light_dark[p]=='light':
                               new_reaction.lower_bound =  estimate_maintenance(mean_light) 
                               new_reaction.upper_bound =  estimate_maintenance(mean_light)
                            else:
                               new_reaction.lower_bound = 1/3 * estimate_maintenance(mean_light) 
                               new_reaction.upper_bound = 1/3 * estimate_maintenance(mean_light)
               
       
                elif reaction.attrib['id']=='Photon_tx': 
                    
                    light_intensity = float(hf.gaussian(p, min_light, max_light, mid_day, sigma, shift_light))
                    
                    if light_intensity > Light_Comp: # above light compensation point
                        light_in = light_intensity
                    else:
                        light_in = 0
                        
                    if  fixed_light == 1:                    
                        new_reaction.lower_bound = light_in
                    else:
                       new_reaction.lower_bound = 0 
                       
                    new_reaction.upper_bound = light_in
                    
                ### block RuBisCO at night    
                elif reaction.attrib['id']=='RIBULOSE_BISPHOSPHATE_CARBOXYLASE_RXN_p': 
                    
                    if 'lower' in reaction.keys():
                        new_reaction.lower_bound = float(reaction.attrib['lower'])
                    if 'upper' in reaction.keys():                      
                        if light_dark[p]=='dark':
                            new_reaction.upper_bound = 0
                        else:
                            new_reaction.upper_bound = float(reaction.attrib['upper'])                  
        
                else:            
                    if 'lower' in reaction.keys():
                        new_reaction.lower_bound = float(reaction.attrib['lower'])
                    if 'upper' in reaction.keys():
                        new_reaction.upper_bound = float(reaction.attrib['upper'])                           
                        
                 
def estimate_maintenance(light_intensity):
    
    """
    Initiated by mpm_set_phase_constraints()
    Calculates light-dependent maintenance cost based on max light intensity

    parameters:
    ------
    light_intensitiy extreacted from xml_rool element or th_dict
 
    """
    
    ATPase = (0.0049 * light_intensity) + 2.7851 
    return float(ATPase)


def mpm_set_balances(new_model, balance, nr_phases, fixed_light, th_dict,  light_dark):
    
    """
    Initiated by mpm_create()
    Create balance constraints according to parameter file

    parameters:
    ------
    - new_model: MPM model
    - balance: current balance xml data
    - nr_phases: total number of phases in model
    - light_dark: list of light/dark phases
    
    example phase:
        
        Enzyme_1 generates - > 1/3 fake A
        Enzyme_2 generates -> 1 fake B 
        Enzyme_3 generates -> 1 fake B
        Enzyme_4 generates -> 1 fake B
        
        Day constraint uses 1 fake A + 1 Fake B -> 1 Light_balance (balance)
        Night constraint uses 1 fake A + 1 Fake B -> 1 Night_balance (balance) 
        
        2 Day_balance + 1 Night_balance =>  are consumed (maintenance constraint)
        
        this example sets day-night ratio to 3:1 (from set_phase constraints) and the ratio 
        of A to B to 3:1 whereby B can come from any of the 3 reactions
    """
    
    if balance.attrib['type'] == 'carbon_storage_mets':
        
        for phase in range(1, int(nr_phases + 1)):

            m_common_name = '{m}_common_{p}'.format(m=balance.attrib['id'], p=phase)
            m_common = cobra.Metabolite(m_common_name, name=m_common_name)                
            new_model.add_metabolites(m_common)            
            r_commonizer_name = '{m}_commonizer_{p}'.format(m=balance.attrib['id'], p=phase)
            r_commonizer = cobra.Reaction(r_commonizer_name, name=r_commonizer_name, subsystem="Balance")
            r_commonizer.notes = {'phase': str(phase).zfill(2)}
            r_commonizer.lower_bound = 0
            r_commonizer.upper_bound = float(balance.attrib['upper'])
            reactants = {}
            
            r_commonizer.add_metabolites({m_common: -1})
            new_model.add_reaction(r_commonizer)
                    
            for reaction in balance:

                if phase < 24:
                    r_select = new_model.reactions.get_by_id('{r}_{p}'.format(r=reaction.attrib['id'], p = str(phase).zfill(2) +'_'+ str(phase+1).zfill(2)))
                else:  
                    r_select = new_model.reactions.get_by_id('{r}_{p}'.format(r=reaction.attrib['id'], p = str(phase).zfill(2) +'_'+ str(1).zfill(2)))
   
                r_select.add_metabolites({m_common: 1 })   
                    

    elif balance.attrib['type'] == 'aa_storage':
        
        for phase in range(1, int(nr_phases + 1)):

            m_common_name = '{m}_common_{p}'.format(m=balance.attrib['id'], p=phase)
            m_common = cobra.Metabolite(m_common_name, name=m_common_name)                
            new_model.add_metabolites(m_common)
            r_commonizer_name = '{m}_commonizer_{p}'.format(m=balance.attrib['id'], p=phase)
            r_commonizer = cobra.Reaction(r_commonizer_name, name=r_commonizer_name, subsystem="Balance")
            r_commonizer.notes = {'phase': str(phase).zfill(2)}
            r_commonizer.lower_bound = 0
            r_commonizer.upper_bound = float(balance.attrib['upper'])
            reactants = {}
            
            r_commonizer.add_metabolites({m_common: -1})
            new_model.add_reaction(r_commonizer)
                    
            for reaction in balance:

                if phase < 24:
                    r_select = new_model.reactions.get_by_id('{r}_{p}'.format(r=reaction.attrib['id'], p = str(phase).zfill(2) +'_'+ str(phase+1).zfill(2)))
                else:  
                    r_select = new_model.reactions.get_by_id('{r}_{p}'.format(r=reaction.attrib['id'], p = str(phase).zfill(2) +'_'+ str(1).zfill(2)))
   
                r_select.add_metabolites({m_common: 1 })   

    elif balance.attrib['type'] == 'maintenance':
        
        reaction_unique = []
        for reaction in balance:
            reaction_unique.append(reaction.attrib['n'])
        reaction_unique = set(reaction_unique)
        for phase in range(1, int(nr_phases + 1)):
            m_common_name = {}
            m_common = {}
            for n in reaction_unique:
                m_common_name[n] = '{m}_common_{n}_{p}'.format(m=balance.attrib['id'], n=n, p=phase)
                m_common[n] = cobra.Metabolite(m_common_name[n], name=m_common_name[n])
                
            new_model.add_metabolites(m_common.values())
            
            r_commonizer_name = '{m}_commonizer_{p}'.format(m=balance.attrib['id'], p=phase)
            r_commonizer = cobra.Reaction(r_commonizer_name, name=r_commonizer_name, subsystem="Balance")
            r_commonizer.notes = {'phase': str(phase).zfill(2)}
            r_commonizer.lower_bound = -1000
            r_commonizer.upper_bound = 1000
            reactants = {}
            
            for n in m_common.values():
                reactants[n] = -1
            r_commonizer.add_metabolites(reactants)
            new_model.add_reaction(r_commonizer)
                    
            for reaction in balance:
                r_select = new_model.reactions.get_by_id('{r}_{p}'.format(r=reaction.attrib['id'], p=str(phase).zfill(2)))
                r_select.add_metabolites({m_common[reaction.attrib['n']]: 1 / float(reaction.attrib['b'])})             

    elif balance.attrib['type'] == 'rubisco':
            
            reaction_unique = []
            for reaction in balance:
                reaction_unique.append(reaction.attrib['n'])
            reaction_unique = set(reaction_unique)
            for phase in range(1, int(nr_phases + 1)):
                m_common_name = {}
                m_common = {}
                for n in reaction_unique:
                    m_common_name[n] = '{m}_common_{n}_{p}'.format(m=balance.attrib['id'], n=n, p=phase)
                    m_common[n] = cobra.Metabolite(m_common_name[n], name=m_common_name[n])
                    
                new_model.add_metabolites(m_common.values())
                
                r_commonizer_name = '{m}_commonizer_{p}'.format(m=balance.attrib['id'], p=phase)
                r_commonizer = cobra.Reaction(r_commonizer_name, name=r_commonizer_name, subsystem="Balance")
                r_commonizer.notes = {'phase': str(phase).zfill(2)}
                r_commonizer.lower_bound = -1000
                r_commonizer.upper_bound = 1000
                reactants = {}
                
                for n in m_common.values():
                    reactants[n] = -1
                r_commonizer.add_metabolites(reactants)
                new_model.add_reaction(r_commonizer)
                        
                for reaction in balance:
                    r_select = new_model.reactions.get_by_id('{r}_{p}'.format(r=reaction.attrib['id'], p=str(phase).zfill(2)))
                    r_select.add_metabolites({m_common[reaction.attrib['n']]: 1 / float(reaction.attrib['b'])})             

    elif balance.attrib['type'] == 'phase':
        
            reaction = balance[0]
              
            r_balance_name = '{m}_balance'.format(m=balance.attrib['id'])
            r_balance = cobra.Reaction(r_balance_name, name=r_balance_name, subsystem="Balance")
            r_balance.notes = {'phase': nr_phases}
            new_model.add_reaction(r_balance)
    
            for phase in range(1, int(nr_phases + 1)):
                m_balance_name = '{m}_balance_{p}'.format(m=balance.attrib['id'], p=phase)
                m_balance = cobra.Metabolite(m_balance_name, name=m_balance_name)
                new_model.add_metabolites(m_balance)
    
                r_select = new_model.reactions.get_by_id('{r}_{p}'.format(r=reaction.attrib['id'],p=str(phase).zfill(2)))
                r_select.add_metabolites({m_balance: 1})               
                if light_dark[phase-1] == 'light':
                    stoichiometry = float(balance.attrib['light'])
                else:
                    stoichiometry = float(balance.attrib['dark'])
                r_balance.add_metabolites({m_balance: -stoichiometry})
                           
    
    elif balance.attrib['type'] == 'light':
        
        if th_dict==None: 
            reaction = balance[0]
            
            if fixed_light == 1:
            
                r_balance_name = '{m}_balance'.format(m=balance.attrib['id'])
                r_balance = cobra.Reaction(r_balance_name, name=r_balance_name, subsystem="Balance")
                r_balance.notes = {'phase': nr_phases}
                new_model.add_reaction(r_balance)
        
                for phase in range(1, int(nr_phases + 1)):
                    m_balance_name = '{m}_balance_{p}'.format(m=balance.attrib['id'], p=phase)
                    m_balance = cobra.Metabolite(m_balance_name, name=m_balance_name)
                    new_model.add_metabolites(m_balance)
        
                    r_select = new_model.reactions.get_by_id('{r}_{p}'.format(r=reaction.attrib['id'],p=str(phase).zfill(2)))
                    r_select.add_metabolites({m_balance: 1})
                    stoichiometry = float(reaction[phase - 1].attrib['b'])
                    r_balance.add_metabolites({m_balance: -stoichiometry})
               
            else:    

                for phase in range(1, int(nr_phases + 1)):
                    r_select = new_model.reactions.get_by_id('{r}_{p}'.format(r=reaction.attrib['id'],p=str(phase).zfill(2)))
                    r_select.upper_bound = r_select.upper_bound * float(reaction[phase - 1].attrib['b'])
                    r_select.lower_bound = 0
                
    else:
        raise AttributeError('Balance type not recognized, should be "maintenance", "phase" or "light".') 


     
def mpm_create_common_water_vapour(new_model, nr_phases):
    """
    Creates a common water vapour flux so it can be a single optimized reaction (optimize minimal flux does not
    support minimization of multiple reactions).
    Initiated by mpm_create()

    parameters:
    ------
    - new_model: MPM model
    - nr_phases: total number of phases in model
    """
    
    new_model.add_metabolites([cobra.Metabolite('WATER_trans_e_common', name='WATER_trans_e_common')])
    
    for phase in range(1, nr_phases + 1): # for all phases generate water vapor reaction and then put together
        reaction = cobra.Reaction('H2O_transpiration_tx_commonizer_{}'.format(str(phase).zfill(2)),
                                  name='H2O_transpiration_tx_commonizer_{}'.format(str(phase).zfill(2)),subsystem='Balance',lower_bound=-1000)
       
        new_model.add_reaction(reaction)
        reaction.add_metabolites({"WATER_trans_e_{}".format(str(phase).zfill(2)): -1,'WATER_trans_e_common': 1})

    common_reaction = cobra.Reaction('H2O_transpiration_tx_common', name='H2O_transpiration_tx_common',subsystem='Balance')
    new_model.add_reaction(common_reaction)
    common_reaction.add_metabolites({'WATER_trans_e_common': -1})
    

def mpm_solve(new_model, rxn2avoid, objective1, analysis_type, constants = None, reversible=None,
                        objective2=None, pareto_range=None, pareto_step_size=None,
                        constraint_reaction=None, constraint_range=None, constraint_step_size=None, upper_lower=None, metric=None):
    
    """
    Solves any of the solving methods:
    - FBA
    - Pareto
    - Pareto + Constraint scan

    parameters:
    ------
    - new_model: MPM model
    - rxn2avoid: Reactions not to include in the minimization of flux optimization (excludes linkers)
    - objective1: First objective (ie. max Phloem output)
    - analysis_type: Any of the above types

    For FBA:
    - reversible: If set to false, the reactions will not be split (faster optimization)

    For Pareto or Constraint Scan:
    - objective2: Second objective (ie. minimize water vapour loss)
    - pareto range: If for whatever reason you don't want the full (0, 1 range)
    - pareto_step_size: Pareto range step size

    For Constraint Scan:
    - constraint_reaction: Reaction you want to scan
    - constraint_range: Range of values for reaction
    - constraint_step_size: Step size for range
    """

    print("\n       Starting {}         ".format(analysis_type))

    for reaction in new_model.reactions:  # Changes default 1000 bounds to a number that glpk interprets as infinite, this increases the solver's efficiency.
        if reaction.upper_bound >= 1000:
            reaction.upper_bound = 9999999999999999999999999
        if reaction.lower_bound <= -1000:
            reaction.lower_bound = -999999999999999999999999

    new_model.objective = {}
    new_model.reactions.get_by_id(objective1).objective_coefficient = 1

    print("Performing pFBA ...")

    if analysis_type.lower() == 'fba' and reversible: 
        
        """
        pFBA
        """
            
        solution = cobra.flux_analysis.pfba(new_model)
        solution.fluxes.to_csv('Results/{}_FBA.csv'.format(new_model.id, metric), sep=',')
    else:
    
        solution = pfba_rxns2avoid(new_model,rxn2avoid)
        # write to file
        #solution.fluxes.to_csv('Results/{}_FBA.csv'.format(new_model.id, metric), sep=',')
      
    if analysis_type.lower() == 'pareto':
             
        """
        Pareto
        """
        
        reaction_obj1 = new_model.reactions.get_by_id(objective1)
        reaction_obj2 = new_model.reactions.get_by_id(objective2)

        new_model.objective = {}
        reaction_obj1.objective_coefficient = 1

        solution=new_model.optimize()
       
        print("\nSolving model (FBA) for determining max Phloem Output...")
        
        max_obj1 = dict(solution.fluxes)[objective1] 
        print("Max {0}: {1}".format(objective1, max_obj1))

        # change objective
        reaction_obj1.objective_coefficient = 0 
        reaction_obj2.objective_coefficient = 1

        print("\nSolving all iterations for Pareto frontier (FBA)...")
        write_method = 'wb'
        
        for pareto in np.arange(pareto_range[0], pareto_range[1], pareto_step_size):
            
            if pareto == 1:
                reaction_obj1.lower_bound = max_obj1 * pareto# * 0.999 # we might need to add a bit of slack as the quadratic optimization is less accurate than the linear couterpart
            else:
                reaction_obj1.lower_bound = max_obj1 * pareto #* 0.999

            # minimise water loss 
            sol=new_model.optimize(objective_sense='minimize') 
            
            # fix this minimal water loss value
            reaction_obj2.bounds = (sol.get_primal_by_id(objective2), sol.get_primal_by_id(objective2))

            if metric=='manhattan':
                
                solution = pfba_rxns2avoid(new_model, rxn2avoid)
           
            elif metric=='euclidean':
                                  
                # make copy because that is easier than reverting all the solver settings
                copy_model = new_model.copy()
                
                FeasTol = float(constants['FeasTol'])
                OptTol = float(constants['OptTol'])

                copy_model.solver.configuration.tolerances.feasibility = FeasTol
                copy_model.solver.configuration.tolerances.optimality = OptTol
                
                rxnlist = [r for r in copy_model.reactions if r.id  not in rxn2avoid]  
    
                obj_vars = list(chain.from_iterable([r.flux_expression**2] for r in rxnlist))            
                copy_model.objective = copy_model.problem.Objective(add(obj_vars), direction='min')
                                
                print('\nSolving quadratic minimisation of sum of fluxes for pareto step ' + str(round(pareto,1)) +'.')
                solution = copy_model.optimize(objective_sense=None)
                
            reaction_obj2.bounds = (0, 9999999999999999999999999) 

            with open(os.path.join('Results', '{}_pareto.csv'.format(new_model.id)),
            write_method) as csv_file:
                wf.write_results(csv_file, new_model, solution, pareto, write_method)
            write_method = 'ab'


def mpm_solve_parameter_scan(new_model, rxn2avoid, objective1, analysis_type, constants = None, reversible=None,
                        objective2=None, pareto_range=None, pareto_step_size=None,
                        constraint_reaction=None, constraint_range=None, constraint_step_size=None, upper_lower=None, metric=None):
    
    """
    Solves any of the solving methods:
    - FBA
    - FVA
    - Pareto
    - Pareto + Constraint scan
    - Customized

    parameters:
    ------
    - new_model: MPM model
    - rxn2avoid: Reactions not to include in the minimization of flux optimization (excludes linkers)
    - objective1: First objective (ie. max Phloem output)
    - analysis_type: Any of the above types

    For FBA:
    - reversible: If set to false, the reactions will not be split (faster optimization)

    For Pareto or Constraint Scan:
    - objective2: Second objective (ie. minimize water vapour loss)
    - pareto range: If for whatever reason you don't want the full (0, 1 range)
    - pareto_step_size: Pareto range step size

    For Constraint Scan:
    - constraint_reaction: Reaction you want to scan
    - constraint_range: Range of values for reaction
    - constraint_step_size: Step size for range
    """

    print("\n       Starting {}         ".format(analysis_type))

    for reaction in new_model.reactions:  # Changes default 1000 bounds to a number that glpk interprets as infinite, this increases the solver's efficiency.
        if reaction.upper_bound >= 1000:
            reaction.upper_bound = 9999999999999999999999999
        if reaction.lower_bound <= -1000:
            reaction.lower_bound = -999999999999999999999999

    new_model.objective = {}
    new_model.reactions.get_by_id(objective1).objective_coefficient = 1

    print("Performing pFBA ...")

    solution = pfba_rxns2avoid(new_model,rxn2avoid)
       
    if analysis_type.lower() == 'pareto':
             
        """
        Pareto
        """
        
        reaction_obj1 = new_model.reactions.get_by_id(objective1)
        reaction_obj2 = new_model.reactions.get_by_id(objective2)

        new_model.objective = {}
        reaction_obj1.objective_coefficient = 1

        solution=new_model.optimize()
       
        print("\nSolving model (FBA) for determining max Phloem Output...")
        
        max_obj1 = dict(solution.fluxes)[objective1] 
        print("Max {0}: {1}".format(objective1, max_obj1))

        # change objective
        reaction_obj1.objective_coefficient = 0 
        reaction_obj2.objective_coefficient = 1

        print("\nSolving all iterations for Pareto frontier (FBA)...")

        ob1_list = []
        ob2_list = []
        ob3_list = []
        p_step_list = []
        
        for pareto in np.arange(pareto_range[0], pareto_range[1], pareto_step_size):
            
            p_step_list.append(pareto)

            try:
                val = 1
                reaction_obj1.lower_bound = max_obj1 * pareto * val                                     
                sol=new_model.optimize(objective_sense='minimize')    
                ob1_list.append(max_obj1 * pareto) 
                ob3_list.append(val)

            except:         
                try:
                    val = 0.999
                    reaction_obj1.lower_bound = max_obj1 * val                                      
                    sol=new_model.optimize(objective_sense='minimize')  
                    ob1_list.append(max_obj1 * pareto)
                    ob3_list.append(val)

                except: 
                     try:
                        val = 0.99 
                        reaction_obj1.lower_bound = max_obj1 * val                            
                        sol=new_model.optimize(objective_sense='minimize')
                        ob1_list.append(max_obj1 * pareto)  
                        ob3_list.append(val)

                     except: 
                        val = 0.95 
                        reaction_obj1.lower_bound = max_obj1 * pareto * val                          
                        sol=new_model.optimize(objective_sense='minimize') 
                        ob1_list.append(max_obj1 * pareto)
                        ob3_list.append(val)
                
            
            reaction_obj2.bounds = (sol.get_primal_by_id(objective2), sol.get_primal_by_id(objective2))
            print('water loss')
            print(sol.get_primal_by_id(objective2))

            ob2_list.append(sol.get_primal_by_id(objective2))
            
            if metric=='manhattan':
                print('done')
                #solution = pfba_rxns2avoid(new_model, rxn2avoid)
            
            elif metric=='euclidean':

                # make copy because that is easier that reverting all the solver settings
                copy_model = new_model.copy()
                
                FeasTol = float(constants['FeasTol'])
                OptTol = float(constants['OptTol'])

                copy_model.solver.configuration.tolerances.feasibility = FeasTol
                copy_model.solver.configuration.tolerances.optimality = OptTol

                rxnlist = [r for r in copy_model.reactions if r.id  not in rxn2avoid]  
    
                obj_vars = list(chain.from_iterable([r.flux_expression**2] for r in rxnlist))            
                copy_model.objective = copy_model.problem.Objective(add(obj_vars), direction='min')
                                
                print('\nSolving quadratic minimisation of sum of fluxes')
                solution = copy_model.optimize(objective_sense=None)
                
            reaction_obj2.bounds = (0, 9999999999999999999999999) 
       
    return p_step_list, ob1_list, ob2_list, ob3_list
        

 

def pfba_rxns2avoid(model,rxns2avoid, fraction_of_optimum=1.0, objective=None, reactions=None):
    
    """Perform basic pFBA (parsimonious Enzyme Usage Flux Balance Analysis)
    to minimize total flux.

    Parameters
    ----------
    model : cobra.Model
        The model
    fraction_of_optimum : float, optional
        Fraction of optimum which must be maintained. The original objective
        reaction is constrained to be greater than maximal_value *
        fraction_of_optimum.
    objective : dict or model.problem.Objective
        A desired objective to use during optimization in addition to the
        pFBA objective. Dictionaries (reaction as key, coefficient as value)
        can be used for linear objectives.
    reactions : iterable
        List of reactions or reaction identifiers. Implies `return_frame` to
        be true. Only return fluxes for the given reactions. Faster than
        fetching all fluxes if only a few are needed.

    Returns
    -------
    cobra.Solution

    """
    reactions = model.reactions if reactions is None \
        else model.reactions.get_by_any(reactions)
    with model as m:
        add_pfba_rxns2avoid(m, rxns2avoid, objective=objective,
                 fraction_of_optimum=fraction_of_optimum)
        m.slim_optimize(error_value=None)
        solution = get_solution(m, reactions=reactions)
    return solution


def add_pfba_rxns2avoid(model, rxns2avoid, objective=None, fraction_of_optimum=1.0):
    
    """Add pFBA objective

    Add objective to minimize the summed flux of all reactions to the
    current objective.

    See Also
    -------
    pfba

    Parameters
    ----------
    model : cobra.Model
        The model to add the objective to
        
    rxns2avoid : reactions not to be included on flux minimisation    
    objective :
        An objective to set in combination with the pFBA objective.
    fraction_of_optimum : float
        Fraction of optimum which must be maintained. The original objective
        reaction is constrained to be greater than maximal_value *
        fraction_of_optimum.
        
    """
    
    if objective is not None:
        model.objective = objective
    if model.solver.objective.name == '_pfba_objective':
        raise ValueError('The model already has a pFBA objective.')
        
    sutil.fix_objective_as_constraint(model, fraction=fraction_of_optimum)   
    rxnlist = [r for r in model.reactions if r.id  not in rxn2avoid]  
    reaction_variables = ((rxn.forward_variable, rxn.reverse_variable)
                         for rxn in rxnlist)

    variables = chain(*reaction_variables)
    model.objective = model.problem.Objective(
        Zero, direction='min', sloppy=True, name="_pfba_objective")
    model.objective.set_linear_coefficients({v: 1.0 for v in variables})


if __name__ == "__main__":

    """    
    # ######################### Settings #############################

    # ### metabolic model and modelling settings
    
    """

    objective_species = 'Tomato' # 'Tomato' , 'Opunita' , 'Arabidopsis'
    cobra_model = 'PlantCoreMetabolism_v1_2_3.xml'
    
    parameter_file = 'model_parameters.xml' 
    climate_file = 'Temperature_and_Humidity_Curves.csv'
    
    new_model_name = 'S_24_' + cobra_model[:-4] + '_'+  objective_species 
     
    constants = { 'deltaC_CO2' : 0.004 , 'D_H2O_0' : 2.13E-05 , 'D_CO2_0' : 1.33E-05 , 'mid_day' : 6, 'deltaT' : 2 , 'FeasTol' : 1e-03 , 'OptTol' : 1e-03  } 
    mid_day = float(constants['mid_day'])
    
    analysis_type = 'pareto' # 'FBA', 'Pareto' 
    metric =  'euclidean' # 'euclidean' or 'manhattan' 
 
    reversible = False
    rxn2avoid =  'Commonizers' #'Linkers and Commonizers' 
    objective1 = 'Phloem_output_tx_01' 

    # Pareto:
    objective2 = 'H2O_transpiration_tx_common'
    pareto_range = (0.1, 1.001)  #  pick a number higher than 1).
    pareto_step_size = 0.1
 
    Light_Comp = 30 # light compensation point

    solver = 'gurobi'    
    
    # parameter_scan 
    parameter_scan = 0
    
    """
    # 
    # ### input data settings and environment-coupled, temporal model construction
    # 
    """
    
    use_real_data = 0 # 1: use real temperature and humidity curve, 0: use generated normal distribution curves    
    fixed_light = 1 # 1 means that the system needs take a bell shaped light absorbtion curve, 0 sets this curve as an upper value
   
    

    # use this part when performing an individual simulation
    
    if parameter_scan==0:
    
        if use_real_data==0:
            wv_dict, th_dict, light_dark, sigma, mean_light = get_generated_data(climate_file,Light_Comp)         
            new_model, new_nr_phases = mpm_create(cobra_model, new_model_name, objective_species, parameter_file, fixed_light, Light_Comp,  wv_dict, th_dict = th_dict, light_dark=light_dark, mid_day = mid_day, sigma = sigma, mean_light = mean_light, solver = solver)
            pf.plot_generated_data(parameter_file, th_dict, sigma, constants) 
            
        elif use_real_data == 1:        
            new_model, new_nr_phases = mpm_create(cobra_model, new_model_name, objective_species, parameter_file, fixed_light, Light_Comp, wv_dict = None, th_dict = None, light_dark = None, mean_light = None, solver = solver)
            
        else:
            print('Data type is invalid')
                
        if rxn2avoid == 'Commonizers':
            rxn2avoid = [x.id for x in new_model.reactions if  'common' in x.id]
            
        if rxn2avoid == 'Linkers and Commonizers':
            rxn2avoid = [x.id for x in new_model.reactions if '_L_' in x.id  or  'common' in x.id]
    
        if rxn2avoid == 'Linkers, Commonizers and gas exchange': 
            rxn2avoid = [x.id for x in new_model.reactions if '_L_' in x.id or 'H2O_tx' in x.id or 'CO2_tx' in x.id or  'common' in x.id ]   
    
        """
        #
        # ### Analysis type
        #
        """
    
        if analysis_type.lower() == 'fba':
            mpm_solve(new_model, rxn2avoid, objective1, analysis_type, reversible=reversible)
            
        elif analysis_type.lower() == 'pareto':             
            mpm_solve(new_model, rxn2avoid, objective1, analysis_type,objective2=objective2,
                      pareto_range=pareto_range, pareto_step_size=pareto_step_size, constants = constants, metric = metric)   
    
        else:
            print('Analysis type is invalid')
        
        
        
        
    #  use this part when performing a parameter scan          
    elif parameter_scan==1: 
                  
        wv_file_name = 'Temperature_dependent_WaterVapour.csv'
        wv_file='Parameters/' + wv_file_name 
        
        reader = csv.reader(filter(lambda row: row[0]!='#', open(wv_file,'r')))
        wv_dict = {}
        for row in reader:
            k, v = row
            wv_dict[k] = v
                
                
        th_file_name = climate_file
        th_file='Parameters/' + th_file_name 
        
        reader = csv.reader(filter(lambda row: row[0]!='#', open(th_file,'r')))
        th_dict = {}
        for row in reader:
            k, v = row
            th_dict[k] = v  
        
        
        iterator = 0
        scan_dict={}
        
        r = 0.3
        th_dict['MinHumidity'] = r               
        s = 0.9 
        th_dict['MaxHumidity'] = s
        
        for p in range(0,30,5): # T min
        
            th_dict['MinT'] = p
            
                     
            for q in range(0,50,5): # T max
                
                 if p <= q:          
            
                    th_dict['MaxT'] = float(q)
                                      
                    light_dark, sigma, mean_light = get_generated_data_parameter_scan(th_dict, Light_Comp) 
                    
                    new_model, new_nr_phases = mpm_create(cobra_model,new_model_name, objective_species, parameter_file, fixed_light, Light_Comp,
                                                          wv_dict, th_dict = th_dict, light_dark=light_dark, mid_day = mid_day, sigma = sigma,
                                                          mean_light = mean_light, solver = solver)
            
                    
                    if rxn2avoid == 'Commonizers':
                        rxn2avoid = [x.id for x in new_model.reactions if  'common' in x.id]
                        
                    if rxn2avoid == 'Linkers and Commonizers':
                        rxn2avoid = [x.id for x in new_model.reactions if '_L_' in x.id  or  'common' in x.id]
        
                    if rxn2avoid == 'Linkers, Commonizers and gas exchange': # H2O_transpiration_tx_common # H2O_transpiration_tx_commonizer
                        rxn2avoid = [x.id for x in new_model.reactions if '_L_' in x.id or 'H2O_tx' in x.id or 'CO2_tx' in x.id or  'common' in x.id ]   
                    
                    p_steps, ob1s, ob2s , ob3s = mpm_solve_parameter_scan(new_model, rxn2avoid, objective1, analysis_type,objective2=objective2, pareto_range=pareto_range, 
                                                    pareto_step_size=pareto_step_size, constants = constants, metric = metric)   
            
        
                    scan_dict[iterator]={} 
                    scan_dict[iterator]['MinHumidity'] = float(r)
                    scan_dict[iterator]['MaxHumidity'] = float(s)
                    scan_dict[iterator]['MinT'] = int(th_dict['MinT'])
                    scan_dict[iterator]['MaxT'] = int(th_dict['MaxT'])
                   
                    scan_dict[iterator]['Pareto Step_'+ str(round(p_steps[0],2))] = p_steps[0]
                    scan_dict[iterator]['Pareto Step_'+ str(round(p_steps[1],2))] = p_steps[1]
                    scan_dict[iterator]['Pareto Step_'+ str(round(p_steps[2],2))] = p_steps[2]
                    scan_dict[iterator]['Pareto Step_'+ str(round(p_steps[3],2))] = p_steps[3]
                    
                    scan_dict[iterator]['Phloem output_' + str(round(p_steps[0],2))] = ob1s[0]
                    scan_dict[iterator]['Phloem output_' + str(round(p_steps[1],2))] = ob1s[1]
                    scan_dict[iterator]['Phloem output_' + str(round(p_steps[2],2))] = ob1s[2]
                    scan_dict[iterator]['Phloem output_' + str(round(p_steps[3],2))] = ob1s[3]
                    
                    scan_dict[iterator]['Water loss_' + str(round(p_steps[0],2))] = ob2s[0]
                    scan_dict[iterator]['Water loss_' + str(round(p_steps[1],2))] = ob2s[1]
                    scan_dict[iterator]['Water loss_' + str(round(p_steps[2],2))] = ob2s[2]
                    scan_dict[iterator]['Water loss_' + str(round(p_steps[3],2))] = ob2s[3]
                    iterator += 1
        
                    if iterator % 10 == 0: 
                        df = pd.DataFrame(scan_dict).T
                        df.to_csv('Results/Parameter_scan/Parameter_scan_results_dict_noICDH_ML_MD_RH_min_'+ str(r)+'_RH_max_'+ str(s) +'.csv', sep=',')
                        
        df = pd.DataFrame(scan_dict).T
        df.to_csv('Results/Parameter_scan/Parameter_scan_results_dict_noICDH_ML_MD_RH_min_'+ str(r)+'_RH_max_'+ str(s) +'.csv', sep=',')
                        
                
            
            
            
            
```
