## Supplementary material for "CAM emerges in a leaf metabolic model under water-saving constraints in different environments": Code: plot_budgets.py

```
#!/usr/bin/env python2
# -*- coding: utf-8 -*-
'''
this file generates budget plots and cummulative plots for the 24 hour cycle for individual compartments or all compartments

'''

import cobra
import pandas as pd
import matplotlib.pyplot as plt
import json
import numpy as np


'''
functions
'''

def resort_data(dframe, pareto):
    
    reframe = dframe[dframe['Pareto'] == pareto] 
    reframe = reframe.sort_values(['Original_ID', 'Phase'])
    try:
        reframe['Flux'] = reframe['Flux'].replace(reframe[reframe['Flux'] < 0.00001]['Flux'].values, 0.0)
    except AttributeError:
        pass       
    reframe = reframe[pd.notnull(reframe['Original_ID'])] # only reactions with original ID i.e., no commonizers
    reframe['Pathway'] = reframe['Pathway'].fillna('NA')
    reframe = reframe[['Phase', 'Original_ID', 'Flux']]
    
    return reframe


def generate_budget(model, data_frame, budget_type, colors, col_dict, i, string, comp, min_flux, show_plot = True): 

    pareto_range = df['Pareto'].unique()
    
    for pareto in pareto_range[6:7]: # here only 70% of the max phloem output
        #  bring data in right order        
        df1 = df[df['Pareto'] == pareto] 
        df1 = df1.sort_values(['Original_ID', 'Phase']) 
        df1 = df1[pd.notnull(df1['Original_ID'])] 
        df1['Pathway'] = df1['Pathway'].fillna('NA') 
        df2 = df1[['Phase', 'Original_ID', 'Flux']]  
        df3 = df2.pivot('Phase','Original_ID')
                
        Budget_dict = dict()
        time_tag = '01' # does not matter - just to find reaction name
        
        #for p in ('c', 'p', 'm', 'x'): # compartment specific or across all compartments
        for p in (comp):  
            
            if budget_type == 'ATP':

                  met = model.metabolites.get_by_id('ATP_' + p + '_' + time_tag)
                  met_a = model.metabolites.get_by_id('aATP_' + p + '_'+ time_tag)     
                  
                  for rxn in met.reactions:        
                      # exclude transporters
                       if rxn.id.__contains__('ATP_AMP_mc') or rxn.id.__contains__('ATP_ADP_mc') or rxn.id.__contains__('ATP_pc') or rxn.id.__contains__('AMP_ATP_xc') or rxn.id.__contains__('ATP_ADP_Pi_pc'): 
                           continue      
                       stoiCoeff = rxn.metabolites.get(met)
                       stoiCoeff_a = rxn.metabolites.get(met_a)
                       Budget_dict[rxn.id] =  stoiCoeff + stoiCoeff_a
              
            elif budget_type == 'CO2':      
   
                  co2 = model.metabolites.get_by_id('CARBON_DIOXIDE_' + p + '_' + time_tag)
                  if p !='x': # no hco3 in x
                      hco3 = model.metabolites.get_by_id('HCO3_' + p + '_' + time_tag)
                      mets=[co2,hco3]
                  else:
                      mets=[co2] 
                      
                  for met in mets:  
                      for rxn in met.reactions:
                          # exclude transporters
                           if rxn.id.__contains__('CO2_xc') or rxn.id.__contains__('CO2_pc') or rxn.id.__contains__('CO2_mc') or rxn.id.__contains__('CO2_ec') or rxn.id.__contains__('RXN0_5224_c'): #  CO2 to Hco3
                               continue                             
                           stoiCoeff = rxn.metabolites.get(met)
                           Budget_dict[rxn.id] =  stoiCoeff 
                       
            elif budget_type == 'NAD(P)H':          

                  nadh = model.metabolites.get_by_id('NADH_' + p + '_' + time_tag)
                  nadph = model.metabolites.get_by_id('NADPH_' + p + '_' + time_tag)
                  mets=[nadh,nadph]
                  
                  for met in mets:               
                      for rxn in met.reactions:                           
                           stoiCoeff = rxn.metabolites.get(met)
                           Budget_dict[rxn.id] =  stoiCoeff 
                           
            elif budget_type == 'Dummy':          

                  met = model.metabolites.get_by_id(string + p + '_' + time_tag)
                  print(met)
            
                  for rxn in met.reactions:                            
                       stoiCoeff = rxn.metabolites.get(met)
                       Budget_dict[rxn.id] =  stoiCoeff                

        # now extract the relevant fluxes     
        plot_mat = pd.DataFrame()
        other_pos = pd.DataFrame()
        other_neg = pd.DataFrame()
        
        for key in Budget_dict:
            df4 = df3['Flux',key[:-3]] * Budget_dict[key] 
            df4.name = key[:-3]
    
            if max(abs(df4)) < min_flux  : 
                # split positive and negtive contribution and add to other
                df_neg = df4.copy()
                df_pos = df4.copy()
                df_neg[df_neg > 0] = 0
                df_pos[df_pos < 0] = 0
                other_pos = other_pos.append(df_pos) 
                other_neg = other_neg.append(df_neg)
           
            else:
                plot_mat = plot_mat.append(df4)
                 # if key does not already have a color
                if key[:-3] not in col_dict.keys():
                     col_dict[key[:-3]] = col[i]
                     i+=1
        
        # split plot mat in positive and negative matrix
        plot_mat_pos = plot_mat.copy()
        plot_mat_neg = plot_mat.copy()
        plot_mat_pos[plot_mat_pos < 0] = 0
        plot_mat_neg[plot_mat_neg > 0] = 0
        
        #kick out empty lines
        plot_mat_pos = plot_mat_pos.loc[~(plot_mat_pos==0).all(axis=1)]
        plot_mat_neg = plot_mat_neg.loc[~(plot_mat_neg==0).all(axis=1)]

        other_pos = other_pos.sum()
        other_pos.name = 'Other_pos'
        
        other_neg = other_neg.sum()
        other_neg.name = 'Other_neg'
        
        plot_mat_pos = plot_mat_pos.append(other_pos)   
        plot_mat_neg = plot_mat_neg.append(other_neg)     

        plot_mat = plot_mat_pos.append(plot_mat_neg) 
        plot_mat.to_csv('Results/' + budget_type +'_budget_' + str(round(pareto,2)) + '.csv')
       
        #  plot
        x=range(1,25)
        y_pos = plot_mat_pos.values
        y_neg = plot_mat_neg.values
        

        plot_mat_pos.sum(1)
        plot_mat_neg.sum(1)

        rxns_pos = plot_mat_pos.axes[0]        
        rxns_neg = plot_mat_neg.axes[0]
        
        rxnsNames_pos = rxns_pos.tolist()
        rxnsNames_neg = rxns_neg.tolist()
        
        # combine
        cols_pos = [col_dict[j] for j in rxnsNames_pos]
        cols_neg = [col_dict[j] for j in rxnsNames_neg]
        
        # combine 24-SOF of each reaction and reaction name
        list_pos1 = plot_mat_pos.sum(1).round(2).tolist()
        list_pos1 = [str(j) for j in list_pos1]
        list_pos2 = rxnsNames_pos        
        val_name_pos = [z + ":       " + y for z,y in zip(list_pos1,list_pos2)]
        
        list_neg1 = plot_mat_neg.sum(1).round(2).tolist()
        list_neg1 = [str(j) for j in list_neg1]
        list_neg2 = rxnsNames_neg       
        val_name_neg = [z + ":       " + y for z,y in zip(list_neg1,list_neg2)]
        
        # budget plots
        fig = plt.figure()
        plt.stackplot(x,y_pos,labels = val_name_pos, colors = cols_pos, baseline='zero') 
        plt.stackplot(x,y_neg,labels = val_name_neg, colors = cols_neg, baseline='zero')     
        plt.title(budget_type + ' Budget - '+ 'Pareto Step '+ str(round(pareto,2)) ,fontsize= 14)  
        
        plt.rcParams['xtick.labelsize']=12
        plt.rcParams['ytick.labelsize']=12
        plt.legend(loc='center left', bbox_to_anchor=(1, 0.5),frameon = False, fontsize = 8)    
        fig.savefig('Results/Plots/' + budget_type + '_'+ p +  '_budget_' + str(round(pareto,2)) + '.png', bbox_inches = 'tight',dpi=600)
        
        # cummulative plot
        fig = plt.figure()       
        vec = plot_mat_pos.sum(0)
        cum_vec = np.cumsum(vec)
        plt.plot(x,cum_vec)
        plt.title(budget_type + ' Cumulative Production - '+ 'Pareto Step '+ str(round(pareto,2)) ,fontsize= 14)      
        plt.ylabel(budget_type + ' cumulative Production in $\mu mol/m^{2}/s$')
        plt.grid()
        
    return col_dict, i
        


'''
Main
'''

new_model = cobra.io.read_sbml_model('Models/S_24_PlantCoreMetabolism_v1_2_3_Tomato_manhattan.xml')
df = pd.read_csv('Results/S_24_PlantCoreMetabolism_v1_2_3_Tomato_pareto.csv', header=0, sep='\t')
df = df[pd.notnull(df['Original_ID'])]

# for unique colorscheme
col = [
'#1f78b4',
'#b2df8a',
'#8ae1ed',
'#bebada',
'#fdb462',
'#d68aed',
'#f5737a',
'#fee08b',
'#e69595',
'#66c2a5',
'#d690b2',
'#3288bd',
'#ffff99',
'#a6cee3',
'#ffffb3',
'#6a3d9a',
'#ff7f00',
'#d9d9d9',
'#9e0142',
'#b3de69',
'#33a02c',
'#f46d43',
'#abdda4',
'#d53e4f',
'#fccde5',
'#fb8072',
'#ccebc5',
'#ffffbf',
'#e6f598',
'#bc80bd',
'#5e4fa2',
'#e89f82',
'#80b1d3',
'#ffed6f']
# initiate color dict so all colors assigned are the same
# prepare reaction-specific color dict
col_dict = dict()
col_dict['Other_neg'] = col[0]
col_dict['Other_pos'] = col[1] 
# color index
i = 2
# later just this one
#col_dict = json.load(file(r'Results/Plots/Color_Dictionary_Stack_Plots.txt'))
#i=40
#i=1

budget_types = ('Dummy','CO2','NAD(P)H','ATP')  # Dummy as place holder for other mets
# here replace the Dummy with metabolite and compartment
string = 'GAP_'
comp = 'c'

for budget_type in budget_types:    
    generate_budget(new_model, df, budget_type, col, col_dict, int(i), string,comp, min_flux = 0.50)

json.dump(col_dict, file(r'Results/Plots/Color_Dictionary_Stack_Plots.txt', 'w'))
```
