## Supplementary material for "CAM emerges in a leaf metabolic model under water-saving constraints in different environments": Code: plot_linker_fluxes_and_pareto_frontier.py

```
#!/usr/bin/env python2
# -*- coding: utf-8 -*-
"""
this file reads pareto analysis data 

plots: 

- CO2 uptake
- malate storage
- starch storage
- rubisco activity
- and other linker fluxes of interest

over a 24 hour periode given certain constraints on relative humidity and temperature

"""

import pandas as pd
import matplotlib.pyplot as plt


file_name = "S_24_PlantCoreMetabolism_v1_2_3_Tomato_pareto.csv"
th_file_name='Temperature_and_Humidity_Curves.csv'


'''
Plot reactions of interest only
'''

df = pd.read_csv('Results/' + file_name, header=0, sep='\t')

reactions_high_flux = ['Photon_tx']
reactions_low_flux = ('RIBULOSE_BISPHOSPHATE_CARBOXYLASE_RXN_p' ,'STARCH_p_L','CIT_v_L','MAL_v_L','cAcids_v_L','CO2_tx')
#reactions_low_flux = ('ISOCITDEH_RXN_m' ,'PEPCARBOX_RXN_c') 

pareto_range = df['Pareto'].unique()

high_flux_color = {'Photon_tx':'gold' }
low_flux_color = {'RIBULOSE_BISPHOSPHATE_CARBOXYLASE_RXN_p':'sandybrown','CO2_tx':'#7c7c7cff',
                  'aMAL_v_L': 'palevioletred','MAL_v_L':'mediumvioletred','CIT_v_L':'purple', 'aCIT_v_L':'mediumpurple',
                  'STARCH_p_L':'seagreen','SUCROSE_v_L':'peachpuff', 'cAcids_v_L': 'mediumvioletred','PRO_v_L':'navy',
                  'ISOCITDEH_RXN_c':'orchid','ISOCITDEH_RXN_m':'mediumseagreen',  'PEPCARBOX_RXN_c': 'lightcoral'}

high_flux_label = {'Photon_tx':'Photon uptake'}
low_flux_label = {'CO2_tx':'CO$_2$ uptake','RIBULOSE_BISPHOSPHATE_CARBOXYLASE_RXN_p':'RuBisCO','MAL_v_L':'Malate Linker Vacuole',
                  'aMAL_v_L': 'aMalate Linker Vacuole','CIT_v_L':'Citrate Linker Vacuole', 'aCIT_v_L':'aCitrate Linker Vacuole',
                  'STARCH_p_L': 'Starch','SUCROSE_v_L': 'Sucrose Linker', 'cAcids_v_L': 'Carboxylic acids                   ','PRO_v_L':'Proline',
                  'ISOCITDEH_RXN_c':'ISOCITDEH_RXN_c','ISOCITDEH_RXN_m':'reverse ISOCITDEH_RXN_m','PEPCARBOX_RXN_c':'PEPCARBOX_RXN_c                              '}


total_transpiration_list = []
total_phloem_output_list = []
total_CO2_tx_list = []

for pareto in pareto_range:
  
    plt.rcParams["figure.figsize"] = [5,2]
    fig, ax1 = plt.subplots()    
    ax1.set_ylim([0,300])
    plt.rcParams['xtick.labelsize']=12
    ax1.set_ylabel(r'Light intensity [$\mu$mol/m$^2$/s]', fontsize = 14)
    ax1.set_xlabel('Hour of the Day (Starting from Sunrise)', fontsize = 14)

    df_1 = df[df['Pareto'] == pareto] 
    df_1 = df_1[['Phase', 'Original_ID','ID','Flux']] 
    
    lines = []    
    low_fluxes={}
    
    for reaction in reactions_high_flux:
        
        flux = df_1[df_1['ID'].str.contains(reaction)] 
        high_flux = flux['Flux'].values.tolist()
           
    ax1.fill_between(range(0,24), high_flux, color= '#FFF483FF', alpha = 0.3)
        
    
    ax2 = ax1.twinx()
    ax2.set_ylabel('Fluxes [$\mu$mol/m$^2$/s]', fontsize = 14)
    ax2.set_ylim([0,35])  
    plt.rcParams['xtick.labelsize']=12
    plt.rcParams['ytick.labelsize']=12
  
    for reaction in reactions_low_flux:
        flux = df_1[df_1['ID'].str.contains(reaction)]
        
        if len(flux) > 24:
            flux = flux[~flux['ID'].str.contains('a')]   # kick out the aForm (different protonation states (see Shameer at al, 2018))
   
        if len(flux) > 24:
            flux = flux[~flux['ID'].str.contains('backwards')]
             
        low_fluxes[reaction] = flux['Flux'].values.tolist()
        
        if reaction == 'cAcids_v_L':

            new_flux_1  = [i * 2 for i in low_fluxes['CIT_v_L']] # Cit and aCit come at a ratio of 1:1 
            new_flux_2  = [i * 10/7 for i in low_fluxes['MAL_v_L']] # Mal and aMal come at a ratio of 7:3       
            low_fluxes['cAcids_v_L'] = [sum(x) for x in zip(new_flux_1, new_flux_2)]       
            ax2.fill_between(range(0,24), low_fluxes['cAcids_v_L'], color= low_flux_color['cAcids_v_L'],alpha=0.3) 
    
        if reaction == 'STARCH_p_L':
            ax2.fill_between(range(0,24), low_fluxes['STARCH_p_L'], color= low_flux_color['STARCH_p_L'],alpha=0.3) 
    
        if reaction == 'ISOCITDEH_RXN_m':     
           new_flux_3  = [i * -1 for i in low_fluxes['ISOCITDEH_RXN_m']]
           low_fluxes['ISOCITDEH_RXN_m'] = new_flux_3
           
        if reaction != 'CIT_v_L' and reaction != 'MAL_v_L':
            
            if reaction == 'CO2_tx':
                lw = 4
            else:
                lw = 1
                
            l = ax2.plot(low_fluxes[reaction], low_flux_color[reaction],label = low_flux_label[reaction],linewidth=lw)  
            lines.append(l[0])  
            labs = [li.get_label() for li in lines]    
            
            ax2.legend(lines, labs, bbox_to_anchor=(1.2, 1), loc=0, frameon=False, borderaxespad=0.0, fontsize=12)

    plt.show()
     
    transpiration_flux = df_1[df_1['ID'].str.contains('H2O_transpiration_tx_commonizer')] 
    total_transpiration = sum(transpiration_flux['Flux'].values.tolist())    
    total_transpiration_list.append(total_transpiration)
    
    phloem_flux = df_1[df_1['ID'].str.contains('Phloem_output_tx')] 
    total_phloem_output = sum(phloem_flux['Flux'].values.tolist())   
    total_phloem_output_list.append(total_phloem_output)
    
    CO2_flux = df_1[df_1['ID'].str.contains('CO2_tx')] 
    total_CO2_tx = sum(CO2_flux['Flux'].values.tolist())   
    total_CO2_tx_list.append(total_CO2_tx)
    
    fig.savefig('Results/Plots/'+ file_name +'_'+ str(round(pareto,1)) + '.png',dpi=600, bbox_inches='tight') 
 


"""
# pareto frontier plot

"""

fig = plt.figure()
plt.rcParams["figure.figsize"] = [4,3]
    
maxVal = total_phloem_output_list[-1]
total_phloem_output_percent  = [str(int(round(p * 1/maxVal * 100,0))) for p in total_phloem_output_list]
total_transpiration_list_scaled = [round(t*1/100) for t in total_transpiration_list]     

MaxTrans = max(total_transpiration_list_scaled)
total_transpiration_list_percent = [t*100/MaxTrans for t in total_transpiration_list_scaled]    

total_phloem_output_percent.reverse()
total_transpiration_list_percent.reverse()

plt.bar(total_phloem_output_percent, total_transpiration_list_percent, color ='#4adbffff') 
plt.axhline(max(total_transpiration_list_percent)/2, color="blue")
plt.ylabel('Water loss in % of maximum',fontsize = 14)
plt.xlabel('Phloem output in % of maximum',fontsize = 14)

plt.rcParams['xtick.labelsize']=14
plt.rcParams['ytick.labelsize']=14
plt.show()

fig.savefig("Results/Plots/Pareto_frontier_"+file_name +"_paper.jpg", dpi=600) 
```
