## Supplementary material for "CAM emerges in a leaf metabolic model under water-saving constraints in different environments": Code: plot_parameter_scan_results.py

```
#!/usr/bin/env python2
# -*- coding: utf-8 -*-
"""
this file plots the results of a parameter scan

"""
import matplotlib.pyplot as plt
import pandas as pd
import math
import seaborn as sns


def roundup(x):
    return int(math.ceil(x / 10.0)) * 10

def rounddown(x):
    return int(math.floor(x / 10.0)) * 10

# define parameters

RH_max = 0.9
RH_min = 0.3

pareto = 'Water loss_0.7'
pareto_1 = 'Water loss_1.0'

# read results file with and without ICDH

file_name1 = 'Parameter_scan_results_dict_ML_MD_RH_min_'+ str(RH_min) +'_RH_max_'+ str(RH_max) +'.csv'
file_name2 = 'Parameter_scan_results_dict_noICDH_ML_MD_RH_min_'+ str(RH_min) +'_RH_max_'+ str(RH_max) +'.csv'

#Pareto_Step = (1,2,3)

df_1 = pd.read_csv('Results/Parameter_scan/' + file_name1, header=0, sep=',')
df_2 = pd.read_csv('Results/Parameter_scan/' + file_name2, header=0, sep=',')


# select only data with same min humidity
df_p1 = df_1[df_1['MinHumidity'] == RH_min] 
df_p2 = df_2[df_2['MinHumidity'] == RH_min] 

# loop over all tested temperatures 
iter_list_all = list(df_2['MaxHumidity'].unique())
# just in case
iter_list =  [i for i in iter_list_all if i <= 1]


for i in range(len(iter_list)):
    print(i)    

    plt.subplot(3, 1, 1)

    RH_max = iter_list[i]

    # we already selected on RH min and now we also select one specific RH max
    df_p1 = df_1[df_1['MaxHumidity']==iter_list[i]]
    df_p2 = df_2[df_2['MaxHumidity']==iter_list[i]]
    
    df_p1.sort_values(by=['MinT', 'MaxT'])
    df_p2.sort_values(by=['MinT', 'MaxT'])
       
    #### 
    # first plot acutual water loss for both scenarios w/wo ICDH
    ####

    plt.plot(df_p1[pareto]/1000, label = 'ICDH', color = '#3da4ab') # from micro to milli
    plt.plot(df_p2[pareto]/1000, label = 'no ICDH', color = '#f6cd61') 
                
    a = math.ceil(max(max(df_p1[pareto]/1000),max(df_p2[pareto]/1000)))

    if a%10 > 5:
        b = 5- (a%10-5)
    else:
        b = 5- a%10
    plt.ylim([0,a + b]) 
   
    plt.rcParams["figure.figsize"] = [10,3]
    #plt.rcParams['xtick.labelsize']=12
    plt.legend(frameon=False, fontsize = 8)
    plt.grid(color='lightgray', linestyle='-', linewidth=0.1)     
    plt.tick_params(axis='x',which='both', bottom=False, labelbottom=False)
    
    #### 
    # second plot realative water loss for both scenarios w/wo ICDH wrt C3
    ####
  
    H2O_loss_percent_df_p1 =  df_p1[pareto].divide(df_p1[pareto_1], fill_value=0) * 100
    H2O_loss_percent_df_p2 =  df_p2[pareto].divide(df_p2[pareto_1], fill_value=0) * 100


#### matrix for absolute values   
M1=df_p1[['MinT', 'MaxT', pareto]]
M1_abs = M1.pivot(index = 'MinT', columns ='MaxT',values = pareto )

M2=df_p2[['MinT', 'MaxT', pareto]]
M2_abs = M2.pivot(index = 'MinT', columns ='MaxT',values = pareto )

a = math.ceil(max(max(M1[pareto]/1000),max(M2[pareto]/1000)))

if a%10 > 5:
    b = 5- (a%10-5)
else:
    b = 5- a%10
plt.ylim([0,a + b]) 

Diff_abs = M2_abs.subtract(M1_abs, fill_value=0) 


### matrix for relative values with respect to C3

df_p1['Water_loss_relative'] =  df_p1[pareto].divide(df_p1[pareto_1], fill_value=0) * 100
df_p2['Water_loss_relative'] =  df_p2[pareto].divide(df_p2[pareto_1], fill_value=0) * 100

M1=df_p1[['MinT', 'MaxT', 'Water_loss_relative']]
M1_rel = M1.pivot(index = 'MinT', columns ='MaxT', values = 'Water_loss_relative')

M2=df_p2[['MinT', 'MaxT', 'Water_loss_relative']]
M2[['MinT','MaxT']] = M2[['MinT','MaxT']].astype(int)

M2_rel = M2.pivot(index = 'MinT', columns ='MaxT', values = 'Water_loss_relative')

c = math.ceil(max(M1['Water_loss_relative']))
d = math.floor(min(M1['Water_loss_relative']))
    
Diff_rel = M2_rel.subtract(M1_rel, fill_value=0) 

# define parameters for plot range
a = roundup(M1_abs.max().max()/1000)# M1 should always have higher water loss than M2 
b = 5
c = roundup(M1_rel.max().max())
d = rounddown(M1_rel.max().max())
e = 0
f = math.ceil(Diff_abs.max().max()/1000)
g = 0
h =  math.ceil(Diff_rel.max().max())

# plot absolute water loss and difference for both scenarios w/wo ICDH rev

plt.subplot(3, 3, 1) # pos 1 in 3x3 pplot
ax = sns.heatmap(M1_abs/1000, cmap="Oranges", vmin=0, vmax=a +b,  xticklabels=False)# cbar = False,
cbar = ax.collections[0].colorbar
cbar.ax.tick_params(labelsize=8)
plt.setp(ax.get_yticklabels(), rotation='horizontal', fontsize=8)
ax.invert_yaxis()
ax.set_xlabel('')
ax.set_ylabel('')
ax.tick_params(bottom=False)
ax.set_facecolor('lightgray')

plt.subplot(3, 3, 2)
ax =sns.heatmap(M2_abs/1000 ,cmap="Oranges", vmin=0, vmax=a +b, yticklabels=False, xticklabels=False)
cbar = ax.collections[0].colorbar
cbar.ax.tick_params(labelsize=8)
ax.invert_yaxis()
ax.set_ylabel('')
ax.set_xlabel('')
ax.tick_params(left=False,  bottom=False)
ax.set_facecolor('lightgray')

plt.subplot(3, 3, 3)
ax = sns.heatmap(Diff_abs/1000,  cmap="Greens", yticklabels=False, xticklabels=False, vmin=e, vmax=f)
cbar = ax.collections[0].colorbar
cbar.ax.tick_params(labelsize=8)
ax.invert_yaxis()
ax.set_ylabel('')
ax.set_xlabel('')
ax.tick_params(left=False, bottom=False)
ax.set_facecolor('lightgray')


# plot relative water loss and difference for both scenarios w/wo ICDH rev

plt.subplot(3, 3, 4) # pos 1 in 3x3 pplot
ax = sns.heatmap(M1_rel, cmap="Blues",  vmin=d, vmax=c )
plt.setp(ax.get_xticklabels(), rotation='horizontal', fontsize=8)
plt.setp(ax.get_yticklabels(), rotation='horizontal', fontsize=8)
cbar = ax.collections[0].colorbar
cbar.ax.tick_params(labelsize=8)
ax.invert_yaxis()
ax.set_xlabel('')
ax.set_ylabel('')
ax.set_facecolor('lightgray')


plt.subplot(3, 3, 5)
ax = sns.heatmap(M2_rel, cmap="Blues",  vmin=d, vmax=c, yticklabels=False )
plt.setp(ax.get_xticklabels(), rotation='horizontal', fontsize=8)
cbar = ax.collections[0].colorbar
cbar.ax.tick_params(labelsize=8)
ax.invert_yaxis()
ax.set_xlabel('')
ax.set_ylabel('')
ax.tick_params(left=False)
ax.set_facecolor('lightgray')

plt.subplot(3, 3, 6)
ax = sns.heatmap(Diff_rel, cmap="Reds", yticklabels=False, vmin=g, vmax=h)
plt.setp(ax.get_xticklabels(), rotation='horizontal', fontsize=8)
cbar = ax.collections[0].colorbar
cbar.ax.tick_params(labelsize=8)
ax.invert_yaxis()
ax.set_ylabel('')
ax.set_xlabel('')
ax.tick_params(left=False)
ax.set_facecolor('lightgray')

plt.savefig('Results/Plots/Heatmaps_'+'RH_max_' + str(RH_max) +' - RH_min_' +  str(RH_min) + '.png',dpi=300, bbox_inches='tight')     

print("Max water loss")
print(round(M1_abs.max().max(),1))

print("Min water loss")
print(round(M1_rel.min().min(),1))

print("Max add. water saving")
print(round(Diff_rel.max().max(),1))

print("Min add water saving")
print(round(Diff_rel.min().min(),1))
```
