## Supplementary material for "CAM emerges in a leaf metabolic model under water-saving constraints in different environments": Code: requirements.txt

We suggest to run the code in a virtual environment, such as miniconda ([https://docs.conda.io/en/latest/miniconda.html](https://docs.conda.io/en/latest/miniconda.html)) using spyder or any other python editor. 

To solve any quadratic optimization problems we used an academic gurobi solver licence (https://www.gurobi.com/downloads/end-user-license-agreement-academic/)

The code was tested with the following specifications:

conda create -n <Environment_name> python=2.7
source activate <Environment_name>

conda install cobra==0.13.4
conda install gurobi (==8.1.0, this is the version we obtained a licence for)
conda install matplotlib
conda install seaborn
